# Uncovering bacterial pseudaminylation with pan-specific antibody tools

**DOI:** 10.1101/2025.07.13.664564

**Authors:** Kristian I. Karlic, Arthur H. Tang, Niccolay Madiedo Soler, Leo Corcilius, Christopher Lehmann, Aleksandra Debowski, Ashleigh L. Dale, Lauren Zavan, Michelle Cielesh, Adedunmola P. Adewale, Karen D. Moulton, Lucy Li, Chenzheng Guan, Maria Kaparakis-Liaskos, Benjamin P. Howden, Ruohan Wei, Xuechen Li, Danielle H. Dube, Stuart J. Cordwell, Mark Larance, Keith A. Stubbs, Glen P. Carter, Nichollas E. Scott, Ethan D. Goddard-Borger, Richard J. Payne

**Affiliations:** Department of Microbiology and Immunology, University of Melbourne at the Peter Doherty Institute for Infection and Immunity, Melbourne 3000, Australia; School of Chemistry, The University of Sydney, Sydney, NSW 2006, Australia; Australian Research Council Centre of Excellence for Innovations in Peptide and Protein Science, The University of Sydney, Sydney, NSW 2006, Australia; The Walter and Eliza Hall Institute of Medical Research, Parkville, Victoria 3052, Australia; Department of Medical Biology, University of Melbourne, Parkville, Victoria 3010, Australia; Marshall Centre for Infectious Disease Research and Training, School of Biomedical Sciences, The University of Western Australia, Nedlands, Western Australia, Australia; School of Molecular Sciences, The University of Western Australia, Crawley, Western Australia, Australia; School of Life and Environmental Sciences, The University of Sydney, Sydney, NSW 2006, Australia; Charles Perkins Centre, The University of Sydney, Sydney, NSW 2006, Australia; School of Medical Sciences, The University of Sydney, Sydney, NSW 2006, Australia; Department of Chemistry and Biochemistry, Bowdoin College 6600 College Station Brunswick, ME 04011, USA; Centre for Pathogen Genomics, The University of Melbourne, Melbourne, VIC, Australia; Department of Chemistry, State Key Laboratory of Synthetic Chemistry, The University of Hong Kong, Pokfulam Road, Hong Kong SAR, P. R. China; Sydney Mass Spectrometry, The University of Sydney, Sydney, NSW 2006, Australia

## Abstract

Pseudaminic acids (Pse) are a family of carbohydrates found within bacterial lipopolysaccharides, capsular polysaccharides and glycoproteins that are critical for the virulence of human pathogens. However, a dearth of effective tools for detecting and enriching Pse has restricted study to only the most abundant Pse-containing glycoconjugates. Here, we devise a synthesis of α- and β-O-pseudaminylated glycopeptides to generate ‘pan-specific’ monoclonal antibodies (mAbs) that recognise α- and β-configured Pse and its C8 epimer (8ePse) presented within glycans or directly linked to polypeptide backbones. Structural characterisation reveals the molecular basis of Pse recognition across a range of diverse chemical contexts. Using these mAbs, we establish a glycoproteomic platform to provide unprecedented depth in mapping the Pse glycome of *Helicobacter pylori*, *Campylobacter jejuni*, and *Acinetobacter baumannii* strains. Finally, we demonstrate that the mAbs recognise diverse capsule types in multidrug-resistant *Acinetobacter baumannii* and enhance phagocytosis to eliminate infections in mice.

## Introduction

Pseudaminic acids (Pse) are nine-carbon ketose sugars (nonulosonic acids) found exclusively in bacteria and archaea. They are structurally similar to neuraminic acid (Neu), the most common sialic acid in humans, but differ in stereochemistry at three carbon atoms (C5, C7 and C8), are deoxygenated at C9, and possess an additional amide functionality at C7 (**Figure 1A**)^1^. While the most commonly encountered Pse derivative is Pse5Ac7Ac, which possesses two N-acetyl groups, a diverse array of N-acyl groups including N-acetamidyl, N-dihydroxypropionyl, N-formyl and N-hydroxybutyryl moieties can adorn the N5 and N7 positions^2^ to tune the physicochemical properties and functions of bacterial glycoconjugates.^3^

**Figure 1.**
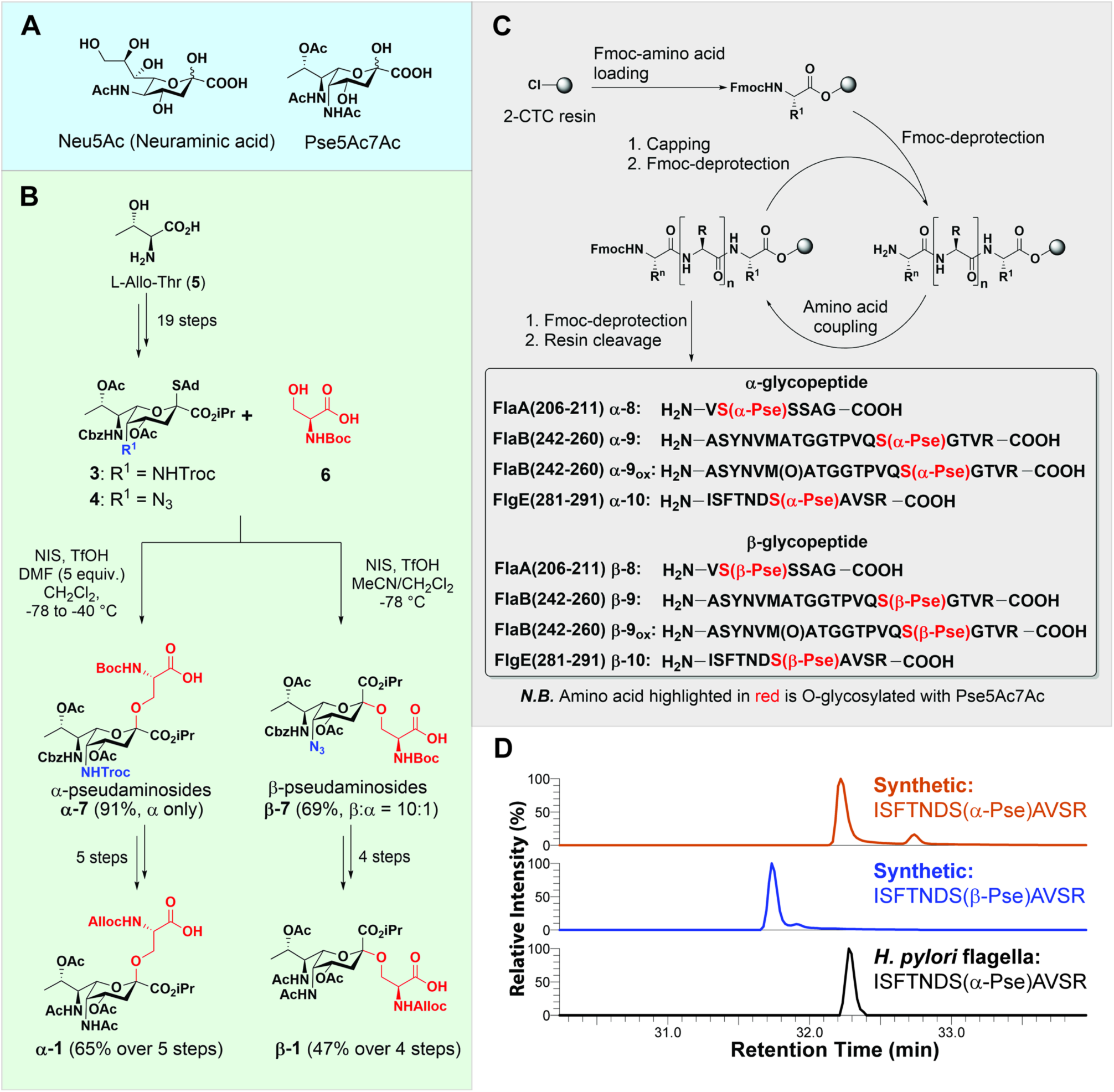
(A) Structure of the common sialic acid derivative Neu5Ac and the common pseudaminic acid derivative – Pse5Ac7Ac. (B) Scheme outlining the synthesis of α-pseudaminylated serine building block **α-1** and β-pseudaminylated serine building block **β-1** from L-allo-threonine (**5**) (see Supplementary Methods for experimental details) (C) Synthesis of flagellin-based glycopeptides discovered by nanoLCMS/MS: α-pseudaminylated glycopeptide **α-8** and tryptic glycopeptides **α-9**, **α-9_ox_**, **α-10**; and β-pseudaminylated glycopeptide **β-8** and tryptic glycopeptides **β-9**, **β-9_ox_**, **β-10**. (D) Extracted ion chromatograms are shown for the singly charged precursor ion after Pse-loss (*z* = +1) at m/z = 1196.5835 ± 5 ppm from the HCD fragmentation of intact Pse-modified precursor peptide ^281^ISFTNDS(Pse)AVSR^291^ (m/z = 756.8632, *z* = +2). Comparison of the retention times for the major peak derived from synthetic tryptic peptide of either the α, or β-configured forms shows a distinct difference in retention time of >0.5 min. The elution time for the tryptic peptide derived from a *H. pylori* flagella extract shows a retention time consistent with the presence of the α-pseudaminylated form of the glycopeptide.

The structural similarity between Pse and Neu is intriguing from both an evolutionary and functional perspective, especially within the context of host-pathogen interactions.^1^ Both molecules are biosynthesised through similar enzymatic cascades and it has been suggested that Pse plays a role in pathogenesis through mimicry of host Neu.^4,5^ For example, bacterial glycoconjugates containing Pse interact with the sialic acid-binding immunoglobulin-type lectin (SIGLEC)-10 receptor on immune cells to dampen immune responses and facilitate prolonged survival and persistence within the host^6,7^.

Pse can be incorporated into lipopolysaccharide (LPS)^8^, capsular polysaccharides (CPS),^9,10^ and glycoproteins in a number of important human pathogens, including the ESKAPE bacteria *Acinetobacter baumannii*^10^ and *Pseudomonas aeruginosa*^8,11,12^; gastrointestinal pathogens *Helicobacter pylori*^13,14^, *Campylobacter coli* and *Campylobacter jejuni*^3,14^ and the periodontal pathogen *Tannerella forsythia*^15^, among others. Pseudaminylated glycoproteins are commonly modified with a single Pse directly O-linked to Ser and Thr residues of proteins in pili^16^ or flagella^13,14,17^. This post-translational modification (PTM) can comprise up to 10% of the mass of flagellin proteins and it is essential for assembly of flagellar filaments, motility and thus virulence in *C. jejuni* and *H. pylori.*^14^ In *H. pylori* and *C. jejuni*, flagellin O-pseudaminylation has been extensively studied, with several investigations suggesting the presence of additional pseudaminylated proteins in these organisms, although these have yet to be identified^18–20^. This alludes to protein pseudaminylation playing a broader functional role than is currently appreciated.

To date, targeted exploration of pseudaminylation has been restricted to the use of metabolic labelling techniques^18,21^. While these methods enable detection of Pse on abundant substrates like flagellin, they have not facilitated the site-specific identification of O-pseudaminylation sites on less abundant substrates. An alternative approach to chemical labelling is the use of modification-specific affinity reagents, such as glycan-binding proteins or antibodies. While Pse-reactive polyclonal antibodies have been raised^22,23^, and an engineered Pse-binding bacteriophage protein has been recently reported^24^, neither tool has delivered new insights into Pse biology. Identifying the full gamut of Pse-modified glycoproteins in bacteria is a crucial next step in refining our understanding of the functional roles that pseudaminylation plays in bacterial physiology. Such insights could underpin future diagnostic and antimicrobial drug discovery efforts to tackle the growing AMR crisis.

Here, we describe the development of ‘pan-specific’ monoclonal antibodies (mAbs) against Pse that enable exploration of the recalcitrant Pse glycoproteome in unprecedented detail. We first developed methodology for the chemical synthesis of Ser and Thr-linked Pse glycopeptide fragments of *H. pylori* flagellin proteins and, through comparisons with native samples, revealed that the glycosidic linkage is α-configured in this organism. These glycopeptides were used to generate mAbs and, through a series of extensive biophysical, immunochemical and structural studies, we show that these tools can recognise both α- and β-configured Pse, as well as its C-8 epimer (8ePse), linked either to a polypeptide or to glycans. These remarkably versatile mAbs were used to enrich glycopeptides from trypsinized protein extracts of *H. pylori*, *C. jejuni* and *A. baumannii*, which enabled the discovery of hitherto unknown Pse-modified proteins in each organism. Finally, we used *in vitro* and *in vivo* infection models to demonstrate the potent therapeutic potential of pan-specific Pse-recognising mAbs against multidrug resistant *A. baumannii*, which is a priority pathogen on the ‘WHO Priority Pathogens List’ and ‘CDC Urgent Threats’ list.

## Results

### Hapten synthesis and assignment of Pse anomeric configuration in H. pylori

Before chemically synthesising O-pseudaminylated peptides as haptens for the generation of Pse-specific antibodies, we first sought to re-evaluate O-pseudaminylation events within the *H. pylori* strain P12 using LC-MS analysis^25^. This analysis identified eight distinct O-linked Pse5Ac7Ac sites on FlaA, six on FlaB as well as three previously unrecognised sites on the flagellin hook protein FlgE (**Supplementary Table 1**). The anomeric configuration of Pse on these peptides remains unknown. As such, we decided to synthesise matched α- and β-configured glycopeptides that could be used both as standards to assign the Pse anomeric configuration in *H. pylori* and as haptens for the selection of pan-specific Pse mAbs.

We chose three glycopeptide targets for synthesis: FlaA(206-211), and the tryptic sequences FlaB(242-260) and FlgE(281-291) identified by the LC-MS analysis above. They possessed single O-linked Pse modifications at Ser207, Ser256 and Ser287, respectively, as confirmed from native flagellin samples (**Extended Data** Figure 1). To access the target glycopeptides by solid-phase peptide synthesis (SPPS), suitably protected α- and β-configured pseudaminylated serine building blocks **α-1** and **β-1** required preparation. Synthesis began with adamantyl thioglycoside donors (**3**) and (**4**) (for **α-1** and **β-1**, respectively), which were prepared in 19-steps from L-*allo*-threonine **5** (**Figure 1B**)^26^. Exclusive α-selectivity was achieved by glycosylating Boc-Ser-O*t*Bu **6** with PseNHTroc donor **3** using 5 equivalents of DMF as an additive in CH_2_Cl_2_ at -78 °C before activation of the thioglycoside donor with NIS and TfOH to afford intermediate glycosylamino acid **α-7** in excellent yield.^26^ Completion of the target pseudaminylated serine building block was achieved via a series of five protecting group removal and functional group manipulation steps (see Supplementary Methods for details) to generate **α-1** in excellent yield. Preparation of β-pseudaminylated serine building block **β-1** began with the glycosylation of Boc-Ser-O*t*Bu **6** with PseN_3_ donor **4** with NIS and TfOH as promoters in a co-solvent system of CH_2_Cl_2_ and MeCN providing a separable 10:1 β:α anomeric mixture. Following purification, **β-7** was generated in excellent yield as a single anomer.^26^ Following four protecting and functional group manipulations (see Supplementary Methods for details) the β-configured pseudaminylated serine building block **β-1** was generated. It should be noted that α- and β-pseudaminylated threonine building blocks were also prepared via a similar method (see Supplementary Methods for synthetic details) but these were not required for our initial experiments.

With **α-1** and **β-1** in hand, we embarked on the assembly of FlaA(206-211) and FlaB(242-260), with and without methionine oxidation, and FlgE(281-291) from the flagellar hook protein (**Figure 1C**). It was envisaged that each of the targets, bearing a single O-linked Pse site, could be synthesised with either α- or β-stereochemistry by installing the differentially configured building blocks **α-1** and **β-1** by solid-phase peptide synthesis (SPPS). Synthesis of each of the target glycopeptides was performed on 2-chlorotrityl chloride (2-CTC) resin and elongation of the target peptide sequence was performed with fluorenylmethoxycarbonyl (Fmoc)-strategy SPPS. Coupling of standard Fmoc-amino acids was performed using ethyl cyano(hydroxyimino)acetate (Oxyma) and *N*,*N*’-diisopropylcarbodiimide (DIC) in DMF and Fmoc deprotection steps were performed by treatment with 20% v/v piperidine in DMF. Coupling of pseudaminylated serine building blocks **α-1** and **β-1** was performed under specialised conditions, namely hexafluorophosphate azobenzotriazole tetramethyl uronium (HATU), 1-hydroxy-7-azabenzotriazole (HOAt) and 2,4,6-trimethylpyridine (TMP) in DMF, in order to suppress epimerisation during coupling of the sterically encumbered glycosylamino acid.^27^ Following the coupling of the pseudaminylated building blocks, the Alloc group was removed from the α-amine under neutral condition with Pd(PPh_3_)_4_ and PhSiH_3_ in CH_2_Cl_2_. After elongation of the glycopeptide sequences the glycopeptides were cleaved off the resin with concomitant deprotection of the side chain protecting groups by treating with a cocktail of TFA/*i*Pr_3_SiH/H_2_O (90:5:5 v/v/v). Finally, the *O*-acetyl groups and isopropyl ester of the Pse moiety on the peptide were removed by mild saponification by treating with 50 mM LiOH solution to obtain the target series of eight α- and β-configured pseudaminylated FlaA, FlaB and FlgE peptides in 1-23% isolated yields following reverse-phase HPLC purification and lyophilisation (over 11-39 steps).

Using the FlgE(281-291) synthetic glycopeptides bearing an α- or β-configured Pse residue we proceeded to perform comparative glycoproteomic analyses with tryptic peptides generated from *H. pylori* (**Figure 1D**). In this experiment, the α-configured synthetic FlgE(281-291) co-eluted with the native tryptic peptide and had a nearly identical fragmentation pattern after HCD fragmentation (Spectral angle 0.98, **Extended Data** Figure 2). In contrast, the β-configured synthetic FlgE(281-291) possessed a different retention time during LC separation and a divergent fragmentation pattern after HCD fragmentation (Spectral angle 0.88, **Extended Data** Figure 2). Taken together, these data unequivocally assign O-linked Pse in *H. pylori* as being α-configured.

### Generation of pan-specific mAbs against Pse

With the synthetic Pse glycopeptides in hand, our next goal was to generate ‘pan-specific’ mAbs capable of recognising O-linked Pse in a broad range of chemical contexts. To this end, we pooled the FlaA, FlaB and FlgE glycopeptides (**α-8**, **α-9** and **α-10**), activated them with Traut’s reagent (2-iminothiolane hydrochloride) and conjugated the resulting products to maleimide-activated KLH and BSA. We used the same conjugation chemistry to link the corresponding non-glycosylated (**8**, **9** and **10**) and β-configured glycopeptides (**β-8**, **β-9** and **β-10**) to maleimide-activated BSA for use in counter-screening. The pooled glycopeptide-KLH conjugates were used to immunise four mice, together with Freund’s complete adjuvant. After three boosters with this antigen and Freund’s incomplete adjuvant, the Pse-reactive antibody titres of these animals were assessed by ELISA with serial dilutions of serum and the pooled glycopeptide-BSA conjugates. The highest titre producing animal was selected for hybridoma generation and supernatants from ∼1000 hybridoma clones screened at low dilution (1:50) by ELISA using the α-Pse glycopeptide-BSA conjugate yielding 46 clones for further evaluation. Higher dilution (1:1000) ELISA against the α-Pse glycopeptide-BSA conjugate, the non-glycosylated peptide-BSA conjugate, and flagellin purified from *C. jejuni* 81-176, which is also extensively modified with O-linked Pse^25^, was used to identify 26 Pse reactive hybridoma candidate lines (**Extended Data** Figure 3A). These candidates were then screened against both α-Pse- and β-Pse-containing synthetic glycopeptides (1:2000) revealing that all clones could recognise both anomeric configurations of Pse (**Extended Data** Figure 3B).

Nine clones were selected for sub-cloning and sequencing with all candidates possessing light chains resulting from IGKV9-123*01 F–IGKJ5*01 F germline allele recombination. Seven clones (1A8, 1B7, 3E11, 4A11, 4H7, 5A7, 9C3) shared a heavy chain comprised of IGHV4-1*01, IGHD1-2*01 F and IGHJ1*03 F alleles, while the remaining two differed subtly in their use diversity (D) alleles: IGHD1-1*01 F for 1E8, and IGHD1-1*01 F for 6D2 (**Figure 2A**). To minimise redundancy and simplify subsequent experiments, we selected three clones: 1E8, 3E11 and 6D2 for further biophysical characterisation. Generation of monovalent Fab of 1E8, 3E11 and 6D2 allowed the determination of binding kinetics and affinities to synthetic N-terminally biotinylated Pse-modified FlgE peptide (Biotin-PEG2-ISFTNDS(α-Pse)AVSR-OH) using biolayer interferometry (BLI) (**Figure 2B**). Robust Pse binding was observed for all clones, with 1E8 possessing the highest affinity (K_D_ = 69 nM) for α-Pse. When reformatted as an scFv, 1E8 bound α-Pse with a similar affinity (K_D_ = 62 nM) and this was used in subsequent structural studies.

**Figure 2.**
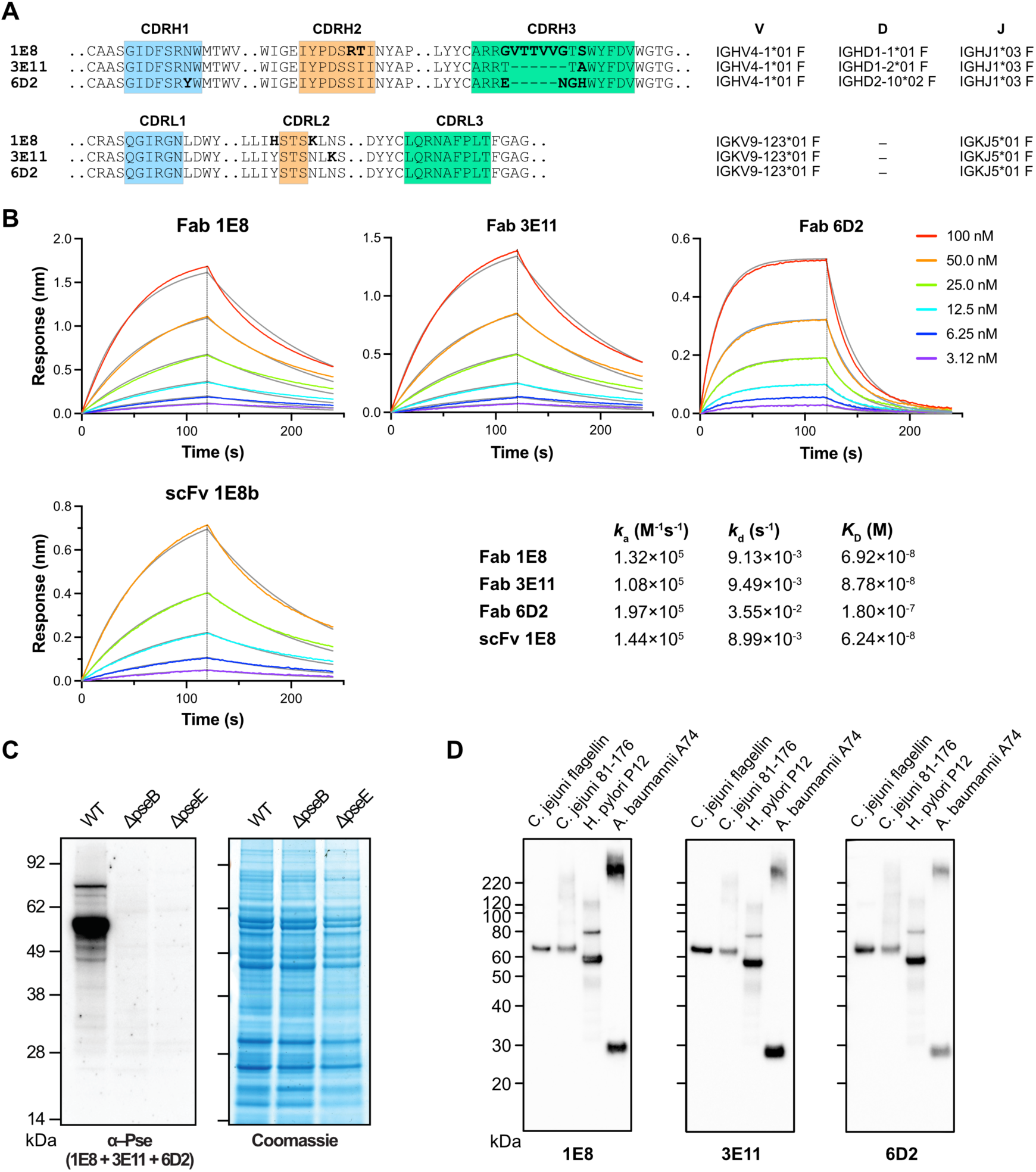
Generation of Pse-specific monoclonal antibodies. (A) Multiple sequence alignment of heavy and light chains from the three clonally distinct Pse-binding mAbs that were selected for further study, together with their IMGT-determined V(D)J allele usage. (B) Representative BLI sensorgrams for the binding of Fab (and an scFv) derived from 1E8, 3E11 and 6D2 to synthetic N-terminally biotinylated Pse-modified FlgE peptide (Biotin-PEG2-ISFTNDS(αPse)AVSR-OH) immobilised on streptavidin-coated BLI tips. The inset table summarises the kinetics and affinity of binding. (C) Western blot using pooled mAbs 1E8, 3E11 and 6D2 to probe cell lysate from wild-type *H. pylori* G27 and two mutant strains deficient in Pse biosynthesis to demonstrate the specificity of these mAbs for pseudaminylation. (D) Representative western blots demonstrating the ability of mAbs 1E8, 3E11 and 6D2 to recognise different forms of Pse in diverse bacterial cell lysates, including O-linked α-Pse5Ac7Ac in *H. pylori* P12 and *C. jejuni* 81-176, as well as glycan-linked α-Pse5Ac7Ac in *A. baumannii* A74.

To assess the specificity of these antibodies, we probed *H. pylori* G27 wild-type and Pse-deficient mutant strains Δ*pseB* and Δ*pseE* ^28,29^ using western blot analysis: this confirmed reactivity only with the wild-type strain (**Figure 2C**). Further western blots were performed with *C. jejuni* 81-176 purified flagellin and lysates, *H. pylori* P12 lysates, and *A. baumannii* A74 lysate (which contains a terminal glycan-linked α-Pse5Ac7Ac) (**Figure 2D**). All three mAbs revealed similar banding patterns across all samples, detecting masses consistent with known glycoconjugates, such as flagellin in *C. jejuni* (65kDa) and *H. pylori* (55kDa) and *A. baumannii* capsular polysaccharide (>250kDa) (**Figure 2D**). Together, these results reveal that these three mAbs are specific for Pse and appear indifferent to its anomeric configuration or aglycone.

### Structural basis of Pse recognition by pan-specific mAbs

To gain insights into Pse recognition, we sought to solve the structure of these antibodies in complex with antigen. We successfully obtained X-ray diffracting crystals of the 1E8 scFv with the synthetic building block Alloc-Ser(α-Pse)-OH (**α-1**), a ligand that we subsequently refer to as ‘Pse-Ser’. Structure determination was achieved by molecular replacement with an Alphafold2 prediction of scFv-1E8(b) serving as a search model. The complex was refined to a resolution of 2.06 Å with a final R_work_/R_free_ value of 0.1929/0.2287 (**Supplementary Table 2**) with all residues in the VL and VH domains modelled. Positive electron density was observed in a cavity formed by the CDR1 and CDR3 loops of the heavy and light chains that enabled placement of the Pse-Ser ligand. The overall structure is typical of an scFv, with VH and VL domains packed tightly against each other and antigen recognition being mediated by the CDRs (**Figure 3A**).

**Figure 3.**
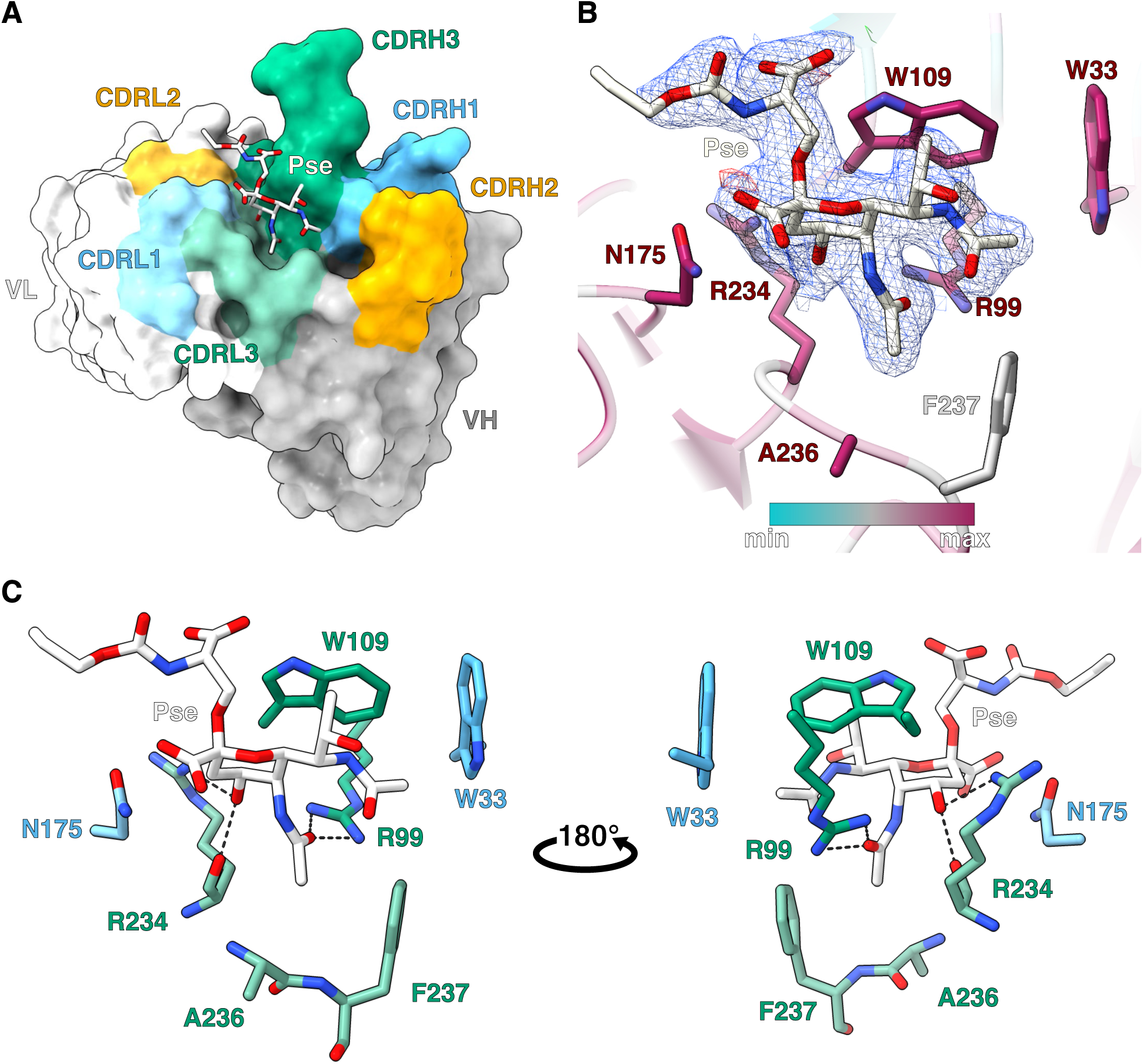
Structural insights into the recognition of Pse5Ac7Ac by mAb 1E8. (A) Overall structure of Alloc-Ser(α-Pse5Ac7Ac)-OH (‘Pse-Ser’, white sticks) bound to 1E8 formatted as an scFv. (B) Pse-Ser bound to 1E8b with a Fo-Fc omit map contoured at 1.5σ around the ligand. Residues in proximity to the pseudaminic acid are coloured by conservation amongst the nine unique mAbs sequenced in this study (maroon is highly conserved, teal is minimally conserved). (C) Key interactions made between Pse and 1E8.

This structure revealed that Pse recognition by 1E8 is mediated by steric fitting, with the anomeric position and aglycone of the Pse facing solvent (**Figure 3A**). These features explain why the anomeric configuration of Pse and the nature of its aglycone are irrelevant to antigen recognition by the mAb. Mapping the sequence conservation between our mAbs (1E8, 3E11 and 6D2) onto the structure demonstrate that the constellation of residues that comprise the antigen binding site are highly conserved (**Figure 3B**), implying that observations from this complex are applicable to all of the mAbs reported here. Key polar interactions between Pse and 1E8 include H-bonds between Arg234 (IMGT V_L_ R107) and O-4, as well as Arg99 (IMGT V_H_ R107) and the carbonyl oxygen of the N-5 acetyl group (**Figure 3C**). This acetyl group sits snugly in a hydrophobic pocket bounded by Ala236 and Phe237 (IMGT V_L_ A113 and F114) and it is unlikely that larger acyl groups, such as the hydroxybutyryl moiety found in *P. aeruginosa* strains^30^, could be accommodated on N-5 by the mAb. In contrast, an open groove near the N-7 acetyl group reveals that larger acyl groups, as is common at this position, could easily be accommodated (**Figure 3A**). Finally, Trp109 (IMGT V_H_ W113) makes CH-π interactions with the Pse pyranose ring and hydrophobic contacts with the C-9 methyl group. The C-8 hydroxyl group is solvent exposed, and its configuration appears to be irrelevant to epitope recognition (*vide infra*). Collectively, these structural data reveal how the pan-specific mAbs can recognise Pse in many different forms but also indicate that they are likely to be intolerant of bulkier modifications at N-5 or O-4.

### Enrichment and glycoproteomics of Pse-modified glycopeptides

Having established the Pse-specific binding features of our mAbs, we turned our attention to applying them as tools to expand knowledge of bacterial pseudaminylation. Previous studies had alluded to the presence of multiple pseudaminylated glycoproteins within *H. pylori*^19,31^, although the localisation of these glycosylation sites beyond FlaA and FlaB remained unknown. Using pooled 1E8, 3E11 and 6D2 mAbs, we performed an immunoenrichment of glycopeptides from trypsin-digested *H. pylori* P12 lysate to facilitate mapping of Pse glycosylation sites using mass spectrometry-based glycoproteomics (**Figure 4A and Supplementary Table 3**). While the majority of pseudaminylated glycopeptides detected were derived from FlaA, FlaB and the flagellar hook protein FlgE (corroborating our initial glycoproteomic analysis of *H. pylori*, **Supplementary Table 1**), the chaperone protein DnaK, glucose-6-phophate isomerase Pgi and HPP12_0570, which belongs to the CopG family of transcriptional regulators were also found to be pseudaminylated (**Figure 4A**). The O-pseudaminylation site within the HPP12_0570 glycopeptide ^65^KGTNQDSSINCDSSSRL^82^ was localised using EThcD to Ser78, further supporting the HCD assignment of this glycopeptide (**Figure 4B, Extended Data** Figure 4). Similar glycopeptide enrichments on trypsin-digested *H. pylori* 26695 lysate also revealed the presence of Pse-modified Pgi and HPP12_0570 (corresponding to HP_0564 within *H. pylori 26695*, **Extended Data** Figure 5 **and Supplementary Table 4**), demonstrating that these hitherto unknown glycosylation sites likely occur across *H. pylori* and are not a strain-specific phenomenon.

**Figure 4.**
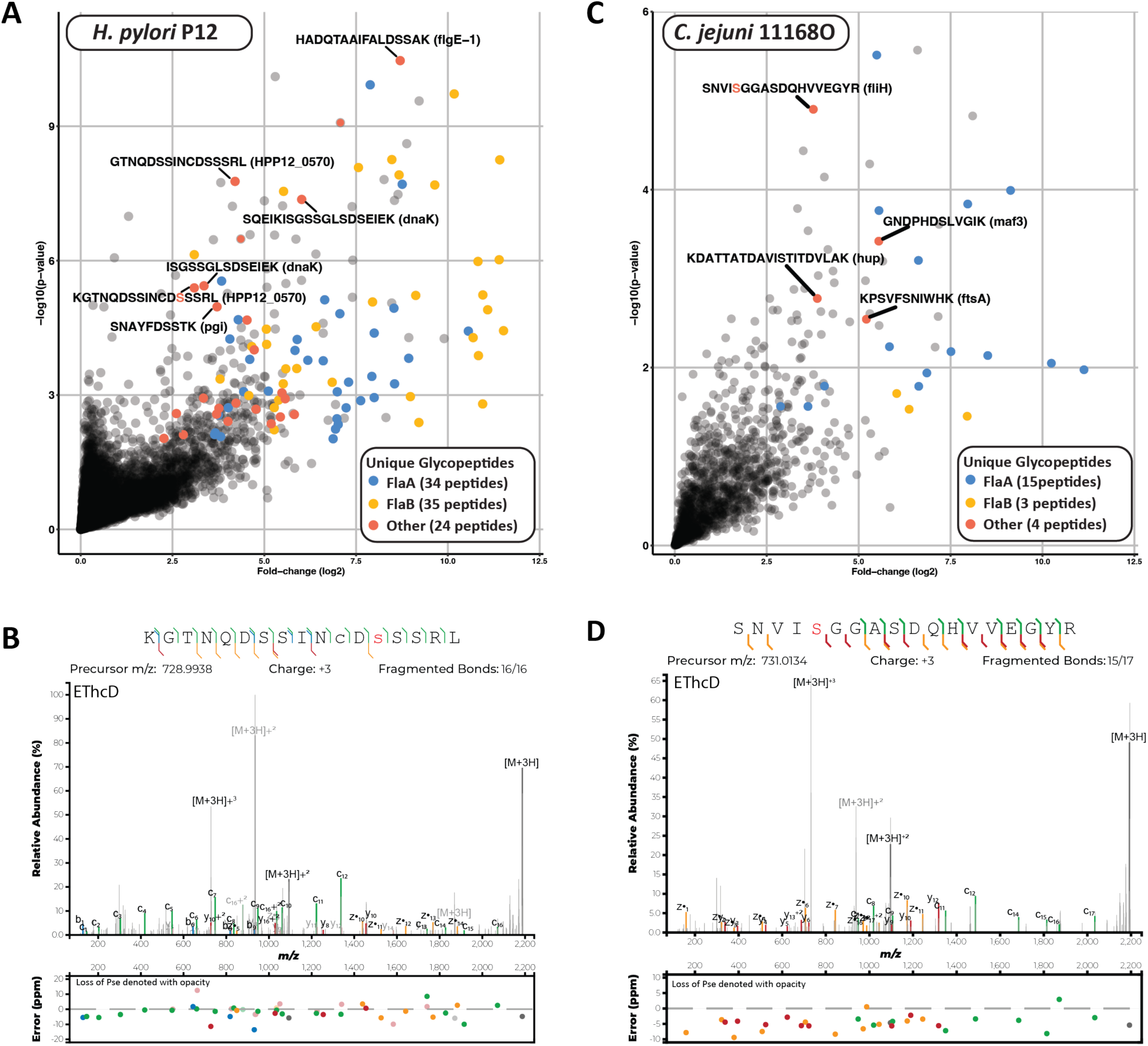
Enrichment of Pse-modified proteins in *H. pylori* and *C. jejuni.* (A) Volcano plot illustrating the enrichment of Pse5Ac7Ac-modified glycopeptides from FlaA/B and proteins that were hitherto unknown to be Pse-modified. (B) EThcD spectra of the novel glycopeptide ^65^KGTNQDSSINCDSSSRL^82^ from HPP12_0570 confirms Pse5Ac7Ac on Ser78. (C) Enrichments of Pse5Ac7Ac-modified glycopeptides reveal multiple FlaA/B derived glycopeptides as well as additional peptide from other proteins. (D) EThcD spectra of the novel glycopeptides ^5^SNVISGGASDQHVVEGYR^22^ from FliH confirming Pse5Ac7Ac on Ser9.

Following the discovery of these new Pse-modification proteins/sites in *H. pylori*, we applied a similar workflow to *C. jejuni*, which is also known to produce FlaA and FlaB that is extensively modified with Pse^32–34^. While it has been suggested that additional O-linked glycoproteins may exist in *C. jejuni*^35^, to our knowledge this has never been confirmed. Consistent with previous studies, *C. jejuni* FlaA and FlaB were readily enriched with the Pse antibody (**Figure 4C, Supplementary Table 5**). However, we also observed Pse sites on several other *C. jejuni* proteins, including flagellar assembly protein FliH (^5^SNVISGGASDQHVVEGYR^22^ **Figure 4D, Extended Data** Figure 6) as well as the putative pseudaminyl transferase (motility accessory factor) Maf3, histone-like DNA-binding protein Hup, and cell division protein FtsA (**Extended Data Figure 7)**. These data confirm the presence of pseudaminylation events beyond the previously reported FlaA/B substrates.

Having demonstrated that these mAbs can facilitate the discovery of glycoproteins with simple O-linked Pse on Ser/Thr residues, we next sought to assess their potential for the immunoenrichment of more complex glycan structures bearing Pse using *A. baumannii*. *A. baumannii* isolates generate different glycan monomers, referred to as K types^36^, which can contain varying nonulosonic acids including Pse ^36,37^ which are used to decorate both O-linked glycoproteins as well as polymerised to form CPS. Western blot analysis of *A. baumannii* strain ATCC19606, a K3 strain that does not generate Pse-containing glycans^37,38^, and *A. baumannii* strain A74^39^, a K2 strain known to produce O-linked glycans and CPS comprised of the α-Pse5Ac7Ac-β-D-Glcp(1→6)[3)-D-Galp-β-D-GalpNAc-(1→]^40^, revealed detection of both CPS and glycoprotein substrates (**Figure 5A**), with proteinase K treatment ablating glycoproteins but not CPS. Analysis of immunoenriched *A. baumannii* A74 glycopeptides using open searching revealed the presence of a 843.31 Da glycoform consistent with the mass of α-Pse5Ac7Ac-β-D-Glcp(1→6)[3)-D-Galp-β-D-GalpNAc-(1→] as the most abundant modification observed based on peptide-spectrum matches (PSM). Dimer masses (1686.62 Da), trimer masses (2529.92 Da), as well as Fe[III] adducts, were also observed supporting the detection of polymerised glycans on protein substrates as previously reported^37^ (**Figure 5B, Supplementary Table 6**). Quantitative analysis (**Figure 5C, Supplementary Table 6**) revealed the enrichment of known glycoproteins of *A. baumannii*^41^ as well as previously unidentified proteins, such as the HlyD family type I secretion periplasmic adaptor subunit (^184^GAASEVEVLR^193^, MDW5584234.1 **Figure 5D**) and the multidrug efflux RND transporter periplasmic adaptor subunit AdeA (^374^APAVANHASSVETK^387^, MDW5585830.1, **Extended Figure 8**) with a further two unique glycopeptides derived from the tetratricopeptide repeat protein (MDW5583480.1) and CvpA family protein (MDW5584485.1) identified in single mAb-mediated enrichment replicates **(Extended Figure 8, Supplementary Table 6)**.

**Figure 5.**
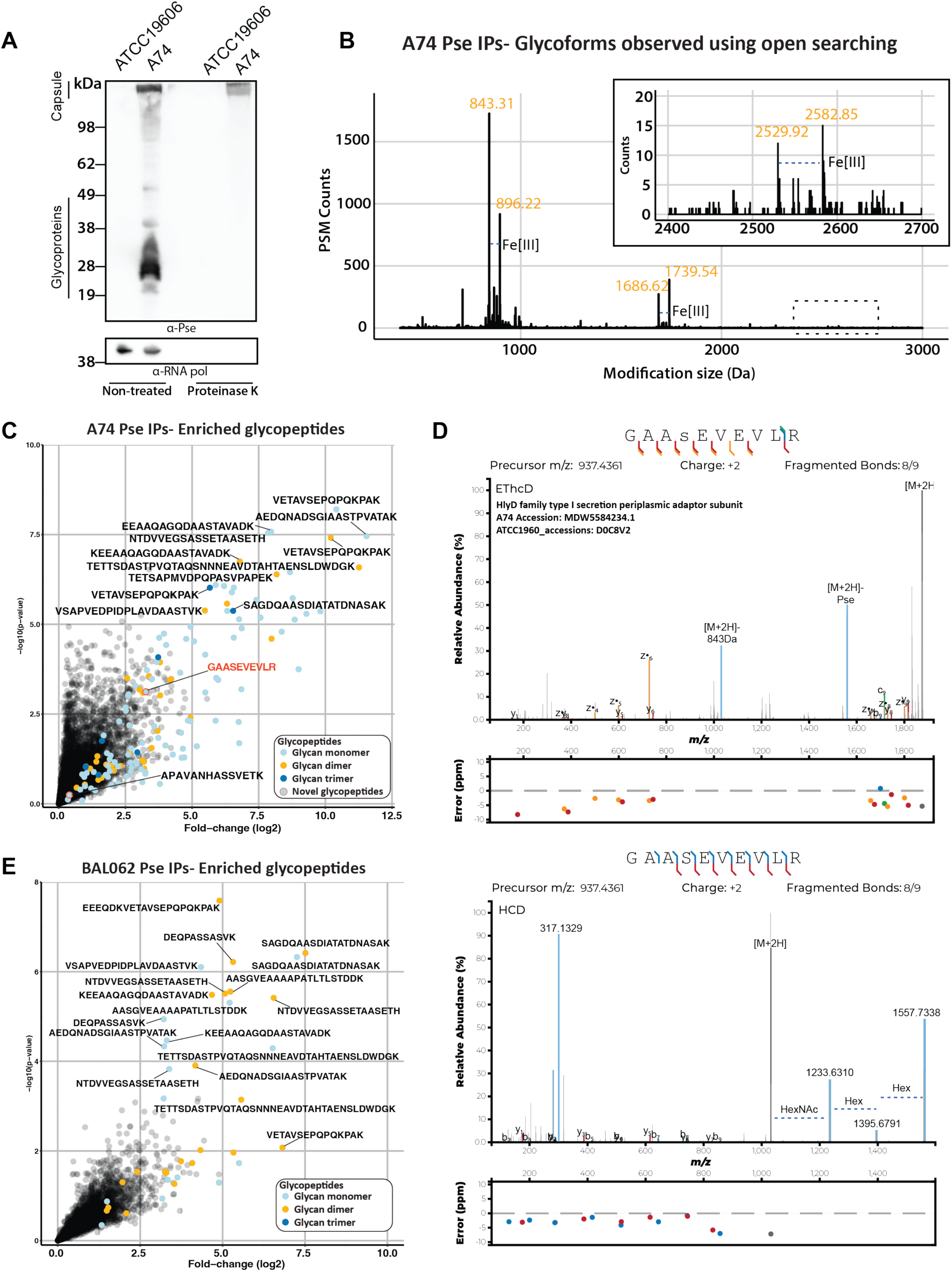
Detection and enrichment of Pse containing glycopeptides from *A. baumannii* strains. (A) Western blotting reveals the detection of both capsule and glycoproteins within the Pseudaminic acid generating strain *A. baumannii* A74. (B) Open searching of *A. baumannii* A74 results reveals multiple glycoforms corresponding to the masses of 843.31 Da, 1686.62 Da and 2529.92 Da as well as Fe[III] adducts of these glycoforms. (C) Volcano plot of pseudaminic acid containing glycopeptides identified from *A. baumannii* A74 revealing the enrichment of multiple glycopeptides, including from previously unreported glycoproteins (denoted with red outlines) using peptide-based enrichment. (D) HCD and EThcD spectra of the previously unrecognised glycopeptides ^184^GAASEVEVLR^193^ (MDW5584234.1) supporting the composition of the O-linked glycan as α-Pse5Ac7Ac-β-D-Glcp(1→6)[3)-D-Galp-β-D-GalpNAc- and localisation of this glycan to Ser187. (E) Volcano plot of glycopeptides identified from *A. baumannii* BAL062 reveals the preferential enrichment of dimer containing glycans using peptide-based enrichment.

To further demonstrate the broad applicability of these mAbs, we assessed if immunoenrichment of glycopeptides modified with 8-epi-pseudaminic acid (8ePse5Ac7Ac) was also possible. This was performed using *A. baumannii* BAL062^42^, which generates a modified K58 monomer corresponding to α-8ePse5Ac7Ac-(2→6)-α-D-Galp-(1→6)-[3-β-D-Glcp-(1→3)-β-D-GalpNAc-(1→]. As with *A. baumannii* A74, immunoenrichment enabled the identification of glycopeptides decorated with glycan monomers, dimers and trimers of known glycoproteins in *A. baumannii* BAL062 (**Figure 5E, Supplementary Table 7**). These results provide yet further evidence that the mAbs 1E8, 3E11 and 6D2 allow the immunoenrichment of structurally diverse glycoconjugates bearing Pse and its C-8 epimer.

### Pan-specific Pse mAbs recognise diverse A. baumannii capsule types

Given the ability of these mAbs to recognise Pse in multiple contexts, we explored their potential as a passive immunotherapy to treat bacterial infections. Multidrug resistant *A. baumannii* is a natural choice for such an intervention, since it is recalcitrant to most other therapies, is commonly acquired in a clinical setting, requires rapid intervention and previous reports have highlighted the effectiveness of CPS-reactive antibodies for controlling *A. baumannii* infections^43–45^. To begin with, we assembled a panel of *A. baumannii* strains with different K types containing Pse with either α- or β-anomeric linkages, as a branching group or part of the polymer backbone, with 4-O-acylation, and as the C-8 epimer. These strains include RBH4 (K6, 4-β-Pse5Ac7Ac-(2→6)-β-D-Galp-(1→6)-β-D-Galp-(1→3)-β-D-GalpNAc-(1→])^46^, NIPH-329 (K46, β-Psep5Ac7Ac4Ac-(2→6)-[3)-α-D-Galp-(1→6)-α-D-GalpNAc-(1→3)-β-D-GalpNAc-])^47^, BAL062 (K58, α-8ePse5Ac7Ac-(2→6)-α-D-Galp-(1→6)-[3-β-D-Glcp-(1→3)-β-D-GalpNAc-(1→])^42^ and A74 (K2, α-Pse5Ac7Ac-(2→6)-β-D-Glcp(1→6)[3)-D-Galp-β-D-GalpNAc-(1→])^48^, with ATCC19606 (KL3) lacking Pse, used as a negative control. A western blot was performed on lysate from these strains (**Figure 6A**).

**Figure 6.**
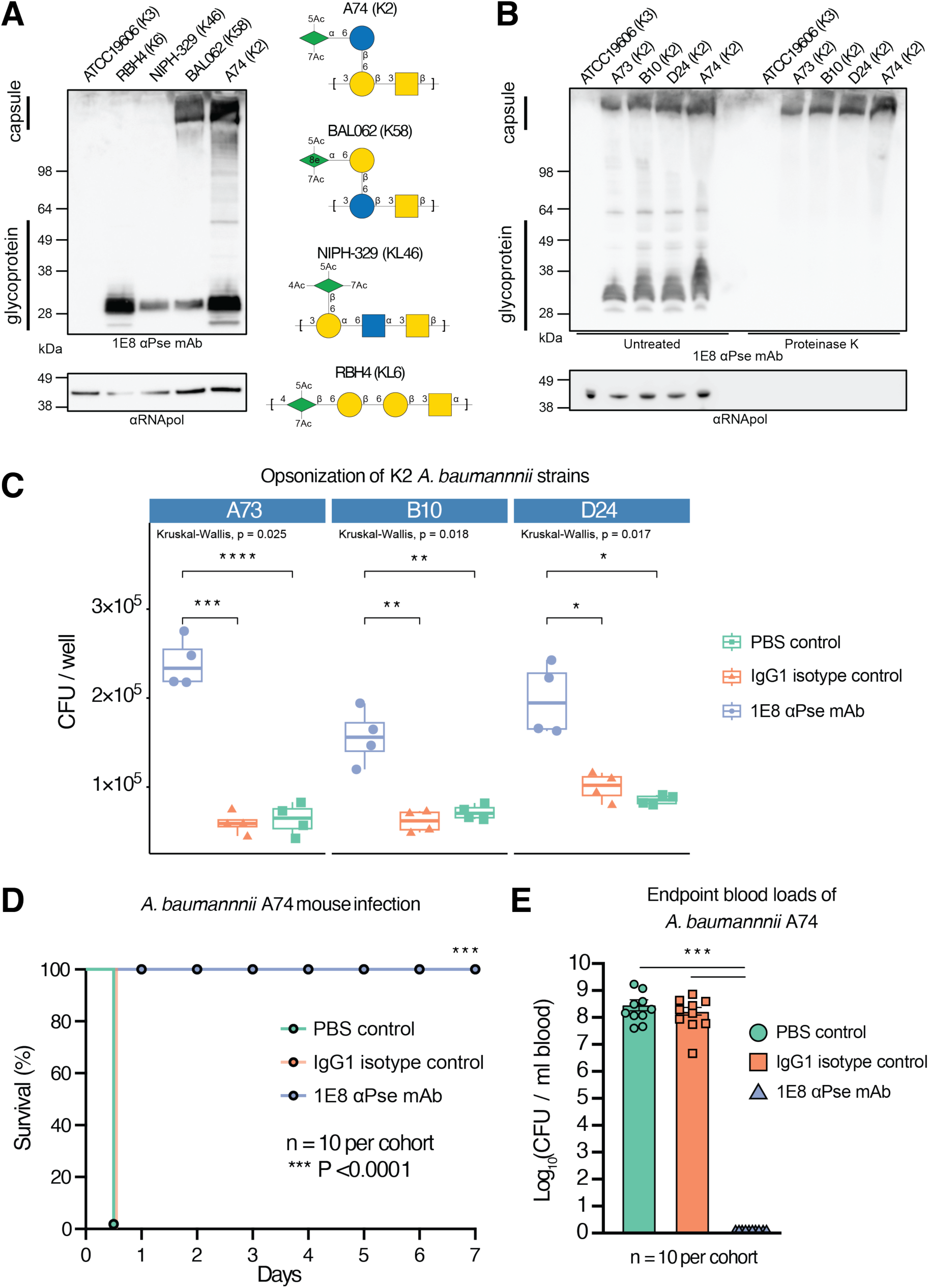
Pse antibodies as tools for detecting and treating *A. baumannii* infections. (A) Western blot probing lysate from *A. baumannii* strains ATCC19606 (K3), RBH4 (K6), NIPH-329 (K46), BAL062 (K58), and A74 (K2) with the Pse mAb 1E8. (B) Western blot analysis of K2 lysates with and without proteinase K treatment. (C) *In vitro* gentamycin protection assays of gentamycin-sensitive K2 strains in response to pretreatment with 1E8 Pse-specific antibody, isotype control, or PBS (n=4 independent experiments for each condition). Box and whisker plots indicate minimum, first quartile, median, third quartile, and maximum values. Kruskal–Wallis tests gave p-values <0.05 (*), <0.01 (**), <0.001 (***) and <0.0001 (****). (D) Kaplan–Meier survival curve of mice infected with 10^5^ cfu of *A. baumannii* A74 administered by intraperitoneal injection (t = 0). One-hour post-infection, mice received either 100 μg of Pse mAb 1E8 (n = 20), 100 µg of mouse IgG1 isotype control (n =10), or PBS (n = 10) via lateral tail vein injection. Animals were monitored out to 7 days with hourly monitoring from 10 to 24 hours followed by daily monitoring. Both the PBS and IgG1 isotype control cohorts reached humane endpoints at 12 hours: these and 10 Pse mAb-treated animals were euthanized and and used to determine 12-hour blood bacterial loads. The remaining 10 Pse mAb 1E8-treated animals were monitored out to 7 days. (E) Bacterial loads (t=12 h) in blood obtained by cardiac puncture of the PBS-treated, IgG1 isotype control-treated, and Pse mAb 1E8-treated mice (n = 10 per group). Error bars represent SD. Unpaired two-tailed t-test gave p-values <0.001 (***).

The mAb 1E8 was able to recognise glycoproteins in all Pse producing strains, yet only recognised the CPS of A74 (K2) and BAL062 (KL58). The reactivity of K2 and K58 CPS supports that the branching α-Pse5Ac7Ac or α-8ePse5Ac7Ac is highly accessible to mAb with analysis of three additional K2 isolates (A73, B10 and D24) demonstrating consistent reactivity of 1E8 across unrelated strains bearing the same capsule type (**Figure 6B**). In contrast, in the K6 strain the β-Pse5Ac7Ac is part of the polymer backbone, with O-4 being glycosylated, thereby preventing binding of the mAb to the capsular polysaccharide. Nonetheless, glycoproteins bearing O-linked K6 units of the capsule have a terminal β-Pse5Ac7Ac and are recognised by the mAb, reiterating the indifference of these antibodies toward anomeric configurations. For K46, the CPS has a branching β-Pse5Ac7Ac4Ac yet is not recognised by the mAb potentially due to the effect of polymerisation and acetylation at O-4, a position which makes important interactions with the protein (**Figure 3C**). Interestingly, the mAb does recognise O-linked K46 units on glycoproteins from this strain, suggesting potential variability in acetylation levels at the O-4 position enabling glycoprotein detection.

### Pan-specific mAbs are effective MDR A. baumannii anti-infectives

Having established that these mAbs recognise different *A. baumannii* isolates, we proceeded to assess their ability to enhance the *in vitro* opsonophagocytosis of gentamycin-sensitive K2 strains (strains A73, B10 and D24) using 5-hour gentamycin protection assays of differentiated THP-1 macrophages.^45,49^ Pre-incubation of bacterial cells from each strain with mAb 1E8 enhanced phagocytosis 2- to 3-fold, relative to a mouse IgG1 isotype control and untreated control (**Figure 6C)**. For the K2 gentamycin-resistant strain A74, enhanced phagocytosis was also observed using 1-hour infection assays, as assessed using immunofluorescence microscopy (**Extended Figure 9**).

Past demonstrations of the anti-infective properties of antibodies against *A. baumannii* CPS have been demonstrated in a prophylactic manner^43,44,50^. We sought to assess the therapeutic potential of our mAbs using a more challenging post-infection model. To this end, C57BL/6 mice were infected with *A. baumannii* A74 (10^5^ cfu per animal) then treated one hour later by IV delivery of either 100 μg of 1E8 mAb, 100 μg of a mouse IgG1 isotype control, or PBS. The 1E8 mAb provided a robust response with all mice surviving infection 7-day post *A. baumannii* infection, while mice administered either isotype control or PBS reached their humane endpoint after 12 hours post-infection. At this time point, both isotype and PBS controls had blood bacterial loads exceeding 10^8^ cfu/ml, while no viable *A. baumannii* A74 was detected in the blood of 1E8 mAb-treated animals (**Figure 6E**). Taken together, these data demonstrate the potent infection-clearing potential of pan-specific mAbs against Pse and further support the application of passive immunisation with capsular polysaccharide-reactive antibodies as an attractive therapeutic approach for combating MDR *A. baumanniii*.

## Discussion

Bacterial protein glycosylation systems are tremendously diverse, both in form and function. As the most prevalent nonulosonic acid used by prokaryotes, Pse is a common structure found on the glycoconjugates of a myriad species^1,2^. The biological roles of Pse are likely to be as diverse as the places it is found; there is no single definitive function for this sugar. As such, robust tools for interrogating the Pse-glycome are valuable assets for probing the glycobiology of many bacterial species, including important human pathogens. The present work has focussed on *H. pylori*, *C. jejuni* and *A. baumannii* as a well-studied test bed to validate our ‘pan-specific’ Pse mAbs as tools for discovery. Despite these systems being well-studied, our mAbs uncovered hitherto unknown glycosylation sites and glycoproteins, providing new avenues of investigation to understand pseudaminylation. This makes these defined reagents powerful assets for studying other bacterial species that produce Pse^1,2^.

The general utility of the mAbs reported here stems from their remarkable ‘pan-specificity’, i.e. their unique ability to recognise Pse in many chemical contexts. Our biochemical, biophysical, structural and glycoproteomic data collectively show that these tools are indifferent to the anomeric configuration, aglycone and C-8 stereochemistry of the Pse. While this broad reactivity is remarkable, there are still limitations to what epitopes the mAbs will recognise. Modifications at O-4 appear poorly compatible with these mAbs and acyl groups larger than an acetyl moiety are unlikely to be accommodated at N-5. This detailed understanding of epitope recognition will inform how best to deploy these mAbs in other bacterial systems.

Beyond interrogating the prevalence of pseudaminylation in bacteria, these pan-specific mAbs have demonstrated promise as anti-infectives against bacteria displaying surface Pse glycoconjugates. The rapid infection-clearing properties of mAb 1E8 against a MDR *A. baumannii* strain is notable because it might reasonably be expected to provide commensurately broad protection against the other capsule (K) types it recognises. Antibodies with similar properties would represent superior alternatives to emerging phage therapies for MDR *A. baumannii*. This suggests that efforts to develop similar pan-specific tools against other bacterial nonulosonic acids, like legionaminic acid, would not only be useful tools for discovery but might also have therapeutic uses.

## Methods

### Chemistry

All synthetic methods and characterisation data for pseudaminylated amino acid building blocks **α-1**, **α-2**, **β-1** and **β-2** and pseudaminylated FlaA, FlaB and FlgE peptide fragments can be found in the Supplementary Methods file.

### Bacterial strains and culture conditions

*H. pylori* P12 ^51^ were grown under microaerobic conditions at 37 °C for 48 h on Columbia blood agar plates containing 5% horse blood and Dent’s antibiotic supplement. *H. pylori* 26695 ^52^ was cultured on Horse Blood Agar (HBA: Blood Agar Base No. 2 (Oxoid), 8% (v/v) horse blood) supplemented with 0.2% Skirrow’s selective supplement (0.0155% (w/v) polymyxin B, 0.625% (w/v) vancomycin, 0.3125% (w/v) trimethoprim and 0.125% (w/v) amphotericin B 12.5% (w/v). *H. pylori* G27 ^49^ wild type, Δ*pseB* and Δ*pseE* strains were grown on Columbia agar base and 5% (v/v) horse blood under microaerobic conditions at 37 °C for 48 h. *H. pylori* mutants bearing the chloramphenicol acetyl transferase (*cat*) cassette (Δ*pseB* and Δ*pseE*) were grown under selection with horse blood agar plates supplemented with 34 µg/mL of chloramphenicol. Cells were collected by centrifugation (4,000×g, 10 min, 4 °C), washed twice with ice-cold PBS (4,000×g, 10 min, 4°C) then flash frozen and stored at -80 °C for proteome sample preparation.

*C. jejuni* NCTC11168^53^ and 81-176^54^ were grown at 37 °C under microaerobic conditions in Mueller-Hinton (MH) broth with 25 µg/mL trimethoprim for 48 h. Cells were subcultured into fresh MH broth at an initial OD_600_ of 0.1 and grown until late exponential phase before being harvested (4,000×g, 10 min, 4 °C), washed twice with ice-cold PBS (4,000×g, 5 min, 4°C) then flash frozen and stored at -80 °C for the isolation of flagellin or for proteome sample preparation.

*A. baumannii strains* ATCC19606 (KL3)^55^, A74 (KL2)^39,40^, A73 (KL2)^48^, B10 (KL2)^48^, D24 (KL2)^48^, RBH4 (KL6)^46^, NIPH-329 (KL46)^47,56^ and BAL062 (KL58)^42,57^ were grown at 37 °C in Lysogeny Broth (LB) overnight with shaking at 185 rpm. Cells were collected by centrifugation (4,000×g, 10 min, 4 °C), washed twice with ice-cold PBS (4,000×g, 10 min, 4 °C) then flash frozen and stored at -80 °C for use in proteome sample preparation.

### Purification of H. pylori P12 flagellin

Flagellar filaments were isolated according to previously described methods. ^58,59^ All manipulations were done on ice or at 4 °C. In brief, cells were harvested in 30 mL cold PBS and washed twice with PBS by pelleting (4000 × *g*; 5 min) and resuspending the cells. The cell pellet was then resuspended in distilled water. Protease Inhibitor Cocktail (Roche) was added to the solution and flagella were removed from the cells by mechanical shearing in a blender (3 × 15s pulses). The sheared cell suspension was centrifuged twice (4,000 × *g*; 20 min) to remove bulk cell material, then centrifuged again (12,000 × *g*; 1 h) to remove residual cell debris. Released flagella was pelleted by ultracentrifugation (100,000 × *g*; 1 h), the flagella pellet resuspended in distilled water and subjected again to ultracentrifugation (100,000 × *g* for 1 h). The resulting flagellum pellet was then collected in distilled water and lyophilized.

### Proteomics of H. pylori flagellin preparations

The flagellum pellet was resuspended in water and subjected to chloroform/methanol precipitation as described ^60^. The protein pellet was resuspended in 50 mL of 4% SDC, 100 mM Tris-HCl (pH 8.5), 10 mM TCEP, 40 mM chloroacetamide prior to heating to 95 °C for 10 min with vortexing at 1,000 rpm. The solution was cooled and flagellar protein (10 μg) was then diluted to 1% SDC with water and trypsin digested (1:20 ratio) overnight at 37 °C. Peptides were purified using SDB-RPS Stagetips as previously described ^61^. Peptides were reconstituted with 5% formic acid in MS-grade water, sealed and stored at 4 °C until LC-MS/MS acquisition. Peptide samples were injected onto a 50 cm × 75 μm C18 (Dr Maisch, 1.9 μm) fused silica analytical column with a 10 μm pulled tip, coupled to a nanospray electrospray ionization source. Peptides were resolved over a gradient from 5% to 35% acetonitrile (MeCN) over 70 min with a flow rate of 300 nL/min using a Dionex Ultimate 3000 UPLC coupled to a Lumos™ Mass Spectrometer. Lumos was operated in a data-dependent mode switching between the collection of a single Orbitrap MS scan (350-1400 *m/z*, maximal injection time of 50 ms, an Automated Gain Control (AGC) of 100% and a resolution of 120k) every second followed by Orbitrap MS/MS HCD scans (NCE 28%, maximal injection time of 54 ms, an a AGC of 100% ions and a resolution of 30k). HCD scans containing the associated Pse oxonium ion (299.1246 *m/z*) triggered an additional product-dependent ^62^ Orbitrap EThcD scan (NCE 15%, maximal injection time of 54 ms, a AGC of 200% and a resolution of 30k). Localisation of FlgE ^281^ISFTNDSAVSR^291^ was undertaken using electron activated dissociation (EAD) PRM on a SCIEX ZenoTOF 7600 targeting the +2-charge state (756.8632 m/z, EAD with 6eV) coupled to an identical LC set up and gradient as outlined above.

Parallel reaction monitoring (PRM) assays to assess the fragmentation of α- or β-configured ^281^ISFTNDS(Pse)AVSR^291^ compared to native glycopeptide was performed using an Exploris 480 mass spectrometer (Thermo Fisher Scientific). The Exploris MS was operated in a data-independent mode switching between the collection of a single Orbitrap MS scan (350-1400 *m/z*, maximal injection time of 50 ms, an Automated Gain Control (AGC) of maximum of 4*10^5^ ions and a resolution of 60k), followed by Orbitrap MS/MS HCD scans of the +2 charge state of ^281^ISFTNDS(Pse)AVSR^291^ (756.8632 m/z, isolation width 0.7 Th, NCE 30%, maximal injection time of 54 ms, and a AGC of 400% ions and a resolution of 30k). The resulting MS2 was manually extracted using the Freestyle Viewer (1.7 SP1, Thermo Fisher Scientific) and MS/MS spectral angle calculated using the Universal Spectrum Explorer^63^. Raw MS data have been deposited to the ProteomeXchange Consortium (http://proteomecentral.proteomexhange.org) *via* the PRIDE partner repository with the dataset identifier PXD056875 (Username, Password: 8cZfCmVbZFQK).

### Conjugation of peptides to carrier proteins

A 260 μM stock of the pseudaminylated peptides (**α-8**, **α-9** and **α-10**) were prepared in 50 mM NaP_i_, 150 mM NaCl, 5 mM EDTA, pH 8. 500 µL of this stock was treated with 32 µL of 14 mM Traut’s reagent (2-iminothiolane hydrochloride) and the solution kept at 22 °C for 1 h. The activated peptide was exchanged into conjugation buffer (100 mM NaP_i_, 150 mM NaCl, 5 mM EDTA, pH 7.2) using a PD MiniTrap G-10 desalting column (GE Healthcare). For conjugation to KLH, 60 µL of 10 mg/mL maleimide-activated KLH (ThermoFisher) resuspended in MilliQ H_2_O was combined with 600 µL of the activated peptide and the reaction incubated at 22 °C for 2 h. The reaction was quenched by the addition of reduced glutathione (final concentration 570 µM) and incubation at 22 °C for 20 min. The KLH conjugate was exchanged into DMPBS using a Zeba spin column (7K MWCO, ThermoFisher). For conjugation to BSA, 40 µL of 10 mg/mL maleimide-activated BSA (ThermoFisher) resuspended in MilliQ H_2_O was combined with 400 µL of the activated peptide and the reaction incubated at 22 °C for 2 h. The reaction was quenched by the addition of cysteine (final concentration 540 µM) and incubation at 22 °C for 20 min. The BSA conjugate was exchanged into DMPBS using a Zeba spin column (7K MWCO, ThermoFisher).

### Antibody generation

Animal experiments complied with the regulatory standards of, and were approved by, The Walter and Eliza Hall Institute of Medical Research Animal Ethics Committee. Mice were kept on a 12/12 light/dark cycle at 20 °C and 50% humidity. At day 0, four mice were immunised with 45 μg of Pse-peptide-KLH conjugate with complete Freund’s adjuvant. Boosters of 45 μg Pse-peptide-KLH conjugate and Freund’s incomplete adjuvant were provided at days 28 and 56. On day 68, the mice were bled and serum titres determined by ELISA using the Pse-peptide BSA conjugate. The mouse with the highest serum titre was provided a 15 μg booster of Pse-peptide KLH conjugate with Freund’s incomplete adjuvant on day 84. On day 88 splenocytes were harvested, fused with a mouse myeloma cell line, and cloned by serial dilution. Supernatants from the resulting hybridomas were screened by ELISA for their ability to bind the Pse-peptide-BSA conjugate. Positive hybridoma supernatants were further screened by ELISA for their ability to bind unglycosylated peptide-BSA conjugate (antigen lacking the pseudaminic acid) and lysate from *C. jejuni* 81-176, which possesses unrelated pseudaminylated proteins. From these experiments, we chose nine hybridoma lines (1B7, 4A11, 1E8, 3E11, 4H7, 6D2, 5A7, 9C3, 1A8) for subcloning by serial dilution, sequencing, and production of purified IgG.

### Purification of C. jejuni flagellin

Flagellin was isolated as described above with minor modification. Cells were re-suspended in water with Protease Inhibitor Cocktail (Roche) and flagella material released by mechanical shearing. The sheared cell suspension was centrifuged twice at 4000×g (20 min, 4 °C) followed by 12,000×g (1 h, 4 °C) to remove bulk cell material. The supernatant was then subjected to ultra-centrifugation (100,000×g, 1 h, 4 °C) and the supernatant discarded. The pellet was re-suspended in water and material recollected by ultra-centrifugation. To obtain pure flagellin, the crude flagella material was pH adjusted to 2.0 (using 0.1 M HCl), and the sample held at 0 °C for 15 min to ensure dissociation of the flagellin proteins. Insoluble material was removed by ultracentrifugation (100,000×g, 1 h, 4 °C). The supernatant was adjusted to pH 7.0 (using 0.1 M NaOH), and the sample held at 0 °C for 15 min to enable re-aggregation of the flagellin protein, which was collected by ultracentrifugation (100,000×g, 1 h, 4 °C). The resulting pellet was resuspended in water then stored at -80 °C.

### Biolayer interferometry

Biolayer interferometry was performed on an Octet RED96 instrument (Sartorius). Kinetic assays were performed at 25 °C with a data acquisition rate of 5.0 Hz and a plate shake speed of 1000 rpm. Biotinylated glycopeptide [Biotin-ISFTNDS(α-Pse)AVSR-OH] was diluted to 10 nM in buffer (50 mM NaPi, 150 mM NaCl, pH 7.4, 0.1% (w/v) BSA, 0.02% Tween 20) then immobilised on an Octet Streptavidin (SA) biosensor (Sartorius) for 10 min at 25 °C. Once a stable baseline was established, the biosensor was dipped into wells containing Fabs (1E8, 3E11, and 6D2) and scFv 1E8b at various concentrations (100, 50.0, 25.0, 12.5, 6.25 and 3.12 nM) and the association observed for 2 min. The biosensor was then transfer to blank buffer and the dissociation of bound Fab or scFv was observed for 2 min. Non-specific interactions of the Fab or scFv with the biosensor were measured and subtracted from the data prior to curve fitting to 1:1 binding model. This provided association (*k*_on_) and dissociation (*k*_off_) rates, as well as the dissociation constant (K_D_). All plots were produced using Prism v10.

### Cloning and expression of Fab and scFv

The mAbs 1E8, 3E11 and 6D2 were reformatted as Fab, and mAb 1E8 also reformatted as an scFv, for periplasmic expression in *E. coli.* A series of dsDNA oligonucleotides encoding these constructs (**Supplementary Table 8**) were synthesised (IDT) and cloned into the pET29b(+) vector (Novagen) at the *NdeI/XhoI* restriction sites. The resulting plasmids were sequence-verified using Sanger sequencing (AGRF).

Plasmids encoding Fab or scFv were transformed into chemically competent NEB T7 Express^®^ *E. coli* cells (NEB), plated onto LB agar + 2% glucose + 50 µg/mL Kan, and incubated at 37 °C for 16 h. Single colonies were picked to grow overnight cultures, which were used to inoculate SB media + 0.2% glucose + 50 µg/ml Kan. Cultures were incubated at 37 °C and 220 rpm until OD_600_ reached 0.8, then IPTG was added to a final concentration of 0.4 mM and the cultures were incubated at 25 °C and 220 rpm for a further 16 h. Cells were harvested by centrifugation (13,800×g, 30 min, 4 °C) then resuspended in ice-cold high osmolyte buffer (0.5 M sucrose, 0.2 M Tris-HCl, pH 8.0) and incubated on ice for 30 min. Three volumes of ice-cold milli-Q water were added and the cells were incubated at 4 °C for 30 min with gentle agitation. Sodium chloride and imidazole were added to a final concentration of 300 mM and 20 mM, respectively. The samples were centrifugated (38,000×g, 45 min, 4 °C) and the supernatants were collected then filtered (0.22 µm).

Filtered supernatants were loaded onto IMAC columns (HisTrap HP columns, GE Healthcare) pre-equilibrated in purification buffer (20mM Tris, 300 mM, 20 mM imidazole, pH 7.4). Columns were washed with 10 CV of purification buffer then eluted with elution buffer (20 mM Tris, 300 mM NaCl, 400 mM imidazole, pH 7.4). Fractions containing the Fab or scFvs (as determined by SDS-PAGE) were further purified by size exclusion chromatography (Superdex® 75 10/300, GE Healthcare) in 25 mM Tris, 150 mM NaCl, pH 7.4, using an ÄKTA Pure system (GE Healthcare).

### Crystallization and structural determination of an scFv-Pse complex

Crystals of the scFv 1E8b complexed to Pse grew over 5 days in sitting drops at 20 °C after mixing 100 nL well solution [2 M (NH_4_)_2_SO_4_, 0.1 M BIS-TRIS pH 5.5] with 100 nL scFv-Pse solution [1:3 molar ratio, 10 mg/ml]. The crystal was cryoprotected in mother liquor supplemented with 30% ethylene glycol and was flash frozen using liquid nitrogen. Diffraction data was collected at the Australian Synchrotron (MX2 beamline) and processed using XDS.^64^ The structure was solved by molecular replacement using PHASER^65^ and a search model generated using AlphaFold2.^65^ The final model was built in Coot ^66^ and refined with Phenix^67^ to a resolution of 2.06 Å. Data collection and refinement statistics are summarized in **Supplementary Table 2**. The R_work_ and R_free_ after the final refinement was 0.1929 and 0.2287, respectively. The coordinates have been deposited in the Protein Data Bank (accession code: 9BEN). Figures were prepared using ChimeraX v1.6.1.^68^

### Inactivation of Pse biosynthesis in H. pylori

Mutagenesis constructs were generated using Gibson assembly^69^ by assembling fragments flanking genes of interest with the chloramphenicol acetyl transferase (*cat*) cassette.^70^ PCR fragments corresponding to the flanking regions of *HpG27_798* (*pseB*) and *HpG27_106* (*pseE*) were amplified by PCR using the oligonucleotides outlined in **Supplementary Table 9** with OneTaq 2x Master Mix (New England Biolabs) from wildtype *H. pylori* strain G27. Amplicons were isolated using a QIAquick Gel Extraction Kit (QIAGEN) and assembled into linear fragments via Gibson assembly before being transformed into *H. pylori* G27 through natural transformation and potential mutants selected for with 25 μg/mL chloramphenicol horse blood agar plates, containing 4% Columbia agar base and 5% laked horse blood. The successful insertion of the resistance cassette was confirmed by PCR and sequencing of PCR products confirming the insertional inactivation of target genes.

### Proteome sample preparation

Lyophilized/flash-frozen cells were resuspended in 4% sodium dodecyl sulfate (SDS), 100 mM Tris pH 8.5, and boiled at 95 °C for 10 minutes with shaking (2000 rpm; Eppendorf ThermoMixer®). ∼1 mg of protein, as determined by bicinchoninic acid protein assays (Thermo Fisher Scientific) was then precipitated by mixing with 4x volume of ice-cold acetone and incubating overnight at -20 °C. Protein samples were then pelleted at 0 °C, the acetone discarded and samples air dried. Precipitated proteome samples were prepared using S-trap mini columns (Protifi, USA) according to the manufacturer’s instructions. Briefly, samples were resuspended in 4% SDS by boiling before being reduced with 10mM DTT at 95 °C for 10 minutes, allowed to cool to room temperature, then alkylated with 40 mM of iodoacetamide for 30 minutes in the dark before being divided across four S-traps (∼250μg each S-trap mini column) per replicate. Samples were then acidified with phosphoric acid to a final concentration of 1.2%, then mixed with seven volumes of 90% methanol/100mM TEAB pH 7.1 before being applied to S-trap mini columns. Samples were washed four times with 90% methanol/100mM TEAB pH 7.1 to remove SDS then Trypsin/Lys-C (1:100, Promega, USA) in 100mM TEAB pH 8.5 spun through the S-trap columns. Samples were digested overnight at 37 °C then collected from the S-traps by washing with 100 mM TEAB pH 8.5 followed by 0.2% formic acid followed by 0.2% formic acid/50% MeCN. Peptide washes were pooled, dried and then resuspended in Buffer A* (0.1% TFA, 2% MeCN) before being cleaned up with home-made high-capacity StageTips composed of 1 mg Empore™ C18 material (3M) and 5 mg of OLIGO R3 reverse phase resin (Thermo Fisher Scientific, USA) as previously described^71,72^. Columns were wet with Buffer B (0.1% formic acid, 50% MeCN) and conditioned with Buffer A* prior to use. Resuspended samples were loaded onto conditioned columns, washed with 10 bed volumes of Buffer A* and bound peptides eluted with Buffer B before being dried then stored at -20°C.

### Affinity enrichment of Pse modified peptides

Affinity enrichment was undertaken utilising the approach of Udeshi *et al.*^73,74^. Briefly Protein G Agarose beads (Abcam) were washed with milli-Q water to remove ethanol and then equilibrated by washing with Immunoprecipitation buffer (IAP; 10 mM Na_3_PO_4_, 50 mM NaCl, 50 mM MOPS, pH 7.2) three times. Protein G Agarose beads were resuspended in 1mL of IAP buffer and either 250 μg Pse mAb mixture or 250 μg IgG1 isotype control added to beads and then tumbled overnight at 4 °C. Coupled protein G mAb beads were washed three times with ice-cold IAP buffer before being resuspend in IAP buffer and evenly split across either three or five microcentrifuge tubes depending on the number of biological replicates. For each enrichment 500 μg of reverse phase cleaned up digests resuspended in IAP buffer were used with the pH confirmed to be pH 7.2 prior to being added to mAb-coupled Protein G beads. Each biological replicate was subjected to enrichment with Pse mAb or IgG1 isotype controls by tumbling at room temperature for 4 h. Protein G beads were washed five times with ice-cold IAP buffer before bound peptides were eluted twice with 0.2% TFA by incubating at 37 °C with shaking at 400 rpm for 10 min. Enriched samples were then cleaned-up with home-made StageTips ^71,72^ composed of Empore™ C18 material and OLIGO R3 reverse phase resin to remove potential Protein G Agarose beads as outlined above and then stored at -20 °C.

### LC-MS analysis

Enriched glycopeptide samples were re-suspended in Buffer A* and separated using a two-column chromatography set up on a Dionex Ultimate 3000 UPLC composed of a PepMap100 C18 20 mm × 75 μm trap and a PepMap C18 500 mm × 75 μm analytical column (Thermo Fisher Scientific) coupled to an Orbitrap Fusion™ Lumos™ Tribrid™ Mass Spectrometer (Thermo Fisher Scientific). 145-minute analytical runs were undertaken with samples concentrated onto the trap column at 5 mL/min for 6 min with Buffer A (0.1% formic acid, 2% DMSO) and then infused into the Orbitrap Lumos™ Mass Spectrometer at 300 nL/minute via the analytical columns. The buffer composition was then altered from 2% Buffer B (0.1% formic acid, 2% DMSO, 78% MeCN) to 28% B over 120 minutes, then from 28% B to 40% B over 9 minutes, then from 40% B to 80% B over 3 minutes, the composition was held at 80% B for 2 minutes, and then dropped to 2% B over 2 minutes and held at 2% B for another 3 minutes. The Lumos™ Mass Spectrometer was operated in a data-dependent mode switching between the collection of a single Orbitrap MS scan (350-2000 *m/z*, maximal injection time of 50 ms, an Automated Gain Control (AGC) of maximum of 4*10^5^ ions and a resolution of 120k) every 3 seconds followed by Orbitrap MS/MS HCD scans (NCE 30%, maximal injection time of 80 ms, an a AGC of 250% ions and a resolution of 30k). HCD scans containing the Pse oxonium ions (317.1338; 299.1233; 281.1128 *m/z*) triggered two additional product-dependent MS/MS scans^62^; a Orbitrap EThcD scan (NCE 15%, maximal injection time of 350 ms, a AGC of 500% and a resolution of 30k with the extended mass range setting used to improve the detection of high mass glycopeptide fragment ions^75^); and a stepped collision energy HCD scan (using NCE 28;32 and 36%, maximal injection time of 250 ms, a AGC of 500% and a resolution of 30k).

### Glycoproteomic Data Analysis

Pse enrichments and the analysis of *H. pylori* P12 flagellin preparation were searched with FragPipe version 19^76–80^. To quantify Pse containing glycopeptides modified “glyco-O-Pair” workflows were used allowing carbamidomethyl as a fixed modification of cysteine in addition to oxidation of methionine and N-terminal acetylation as variable modifications. For *H. pylori* P12 Pse, *H. pylori* P12 26695 and *C. jejuni* NCTC 11168, Pse modifications were defined as mass offsets of 316.12704 Da, 632.2541 Da, 948.38116 Da on serine or threonine residues with Pse diagnostic fragments set as 317.1338; 299.1233; 281.1128 m/z. For *A. baumannii* BAL062 and A74 the mass offsets were set to 1686.6244 Da, 1739.5363 Da and 843.3122 Da on serine residues with the diagnostic fragment ions set as 317.1338; 299.1233; 281.1128, 204.0867, 419.0508, 479.1868 m/z. For *A. baumannii* A74 open searches were undertaken using a modified “open” workflow where the delta mass allowed was set to -10 to 2500 Da as previously described^81^ to identify Pse modified O-linked glycans. Searches were performed against either the *C. jejuni* NCTC 11168 proteome (Uniprot Accession: UP000000799, 1623 proteins); the *H. pylori* P12 proteome (Uniprot Accession: UP000008198, 1572 proteins); the *H. pylori* 26695 proteome (Uniprot Accession: UP000000429, 1554 proteins); the *A. baumannii* BAL062 proteome (NCBI assembly GCA_900088705.1, 3849 proteins) and the *A. baumannii* A74 proteome (NCBI assembly GCA_033541955.1, 3527 proteins) depending on the biological samples all with a 1% FDR. To enhance the detection of glycopeptides across samples match between runs was enabled. The resulting “combined_modified_peptide.tsv” files were processed using Perseus (version 1.6.0.7)^82^ with missing values imputed based on the total observed protein intensities with a range of 0.3 σ and a downshift of 1.8 σ. Statistical analysis was undertaken in Perseus using two-tailed unpaired T-tests. Spectra of assigned glycopeptides of interest were annotated with the aid of the Interactive Peptide Spectral Annotator tool^83^. All mass spectrometry data (RAW files and FragPipe outputs) have been deposited into the PRIDE ProteomeXchange repository^84,85^ with the data identifier PXD046836 (Username: Password: JkoHF9oP) associated with *H. pylori* P12 enrichments; PXD048493 (Username: Password: 8KOCxBaC) associated with *H. pylori* 26695 enrichments; the data identifier PXD046858 (Username: Password: mmG9f4SP) associated with *C. jejuni* NCTC 11168 enrichments; the data identifier PXD046859 (Username: Password: UsOXnSlC) associated with *A. baumannii* BAL062 enrichments and the data identifier PXD053904 (Username: Password: j689tJpTS3U0) associated with *A. baumannii* A74 enrichments.

### In vitro A. baumannii gentamycin infection assays

THP-1 cells grown in RPMI + 10% FBS were differentiated for three days at 37 °C with 5% CO_2_ using 100 nM phorbol-12-myristate-13-acetate in 24-well plates for infection assays. Overnight cultures of *A. baumannii* A73, B10, D24 were standardised using an OD 600nm of 1.0 (∼3.5 x 10^9^ cells/mL) and washed three times with 1 mL of prewarmed RPMI + 10% FBS. Bacterial cells were resuspended into 1 mL of prewarmed of fresh RPMI + 10% FBS and diluted to create 3 x aliquots of 2 x 10^8^ cells/mL. To opsonise bacteria, cells were treated with either 20 µg/mL ^86^ of α-Pse, isotype control or no antibody, and incubated at 37 °C with shaking at 600 rpm for 30 min. An MOI of 20 was used to infect 24-well plates with 1x10^6^ THP-1 macrophage cells per well using (2 x 10^7^ cells/mL) of antibody or non-antibody treated bacterial suspension in biological quadruplicate. Plates were centrifuged at 300 x *g* for 5 min to synchronize the infection. All infected cells were incubated at 37 °C with 5% CO_2_ for 1 hr. After incubation the wells were washed three times with 1 mL of RPMI + 10% FBS, then RPMI + 10% FBS containing Gentamycin 50 μg/mL added to all wells, and incubated at 37 °C with 5% CO_2_ for 30 min. Following 30 min the media was replaced with 1 mL of RPMI + 10% FBS containing Gentamycin 10 μg/mL, followed by a further 4 hours of incubation at 37 °C with 5% CO_2_. Following incubation, wells were washed with 1 mL of RPMI + 10% FBS, and then macrophages cells lysed with 1 mL of 1% Triton in PBS for 30 min. Replicates were plated on LB agar and incubated at 37 °C overnight. CFU/ml were calculated from numerated plates and expressed as CFU/well with 4 biological replicates undertaken.

### Immunofluorescence microscopy of A. baumannii in vitro infections

THP-1 cells grown in RPMI + 10% FBS were differentiated for three days at 37 °C with 5% CO_2_ using 100 nM phorbol-12-myristate-13-acetate in 24-well plates for infection assays. Overnight *A. baumannii* A74 cultures were normalised to 1.0 OD_600nm_ (5 x 10^8^ cells/mL) then washed three times with prewarmed RPMI + 10% FBS before being applied to THP-1 macrophage cells (1x10^6^ cell per well). Infections were undertaken using an MOI of 20 (2 x 10^8^ cfu/mL) with plates centrifuged at 300 x *g* for 5 min to synchronize infections. Opsonisation was undertaken as above with either 20 µg/mL of α-Pse, isotype control or no antibody at 37 °C for 30 min with shaking at 600 rpm before being added to seeded THP-1 macrophages and plates centrifuged at 300 x *g* for 5 min to synchronize infections. All infected cells were incubated at 37 °C with 5% CO_2_ for 1 hr. After incubation wells were washed three times with 1 mL of RPMI + 10% FBS, then fixed with 4% prechilled paraformaldehyde for 15 min at room temperature. Fixed cells were then washed 3 times with pre-chilled PBS, followed by permeabilization with PBS with 1% BSA and 0.2% Triton X-100 for 1 hr at room temperature. Fixed cells were then blocked with PBS with 1%BSA and 0.1% Tween-20 for 1 hr at room temperature and α-Pse primary antibody (1:500 α-Pse in TBST 1% BSA) for 1 hr. Fixed cells were then washed three times with pre-chilled PBS, followed by incubation with secondary antibody, Goat α-mouse Alexa Fluor 488 (Invitrogen, 1:2000 in TBST with 1% BSA) for 1 hr in the dark. Cells were then washed three times with pre-chilled PBS, and then stained for 15 min with Hoechst 33342 (0.1 µg/mL, Thermo) and SiR-Actin (0.2 µg/mL, Spirochrome) in the dark for 20 min. Stained cells were then washed five times with pre-chilled PBS, then mounted onto glass slides with ProLong™ Gold Antifade Mountant (Invitrogen), before being set overnight at room temperature in the dark before being sealed with nail polish. Images were acquired using a Zeiss LSM780 confocal microscope with oil immersion and a 63x objective with images visualized using FIJI ^87^. Four biological replicates were assessed with a minimum of 100 infected cells counted per replicates for quantification of macrophage opsonophagocytosis.

### Animal infection

Eight-week-old female and male C57BL/6 mice were acquired from The Peter Doherty Institute for Infection and Immunity (PDI) and Australian BioResources (ABR) and housed in the pathogen-free facility at The Peter Doherty Institute for Infection and Immunity under a 12-hour light/dark cycle maintained at ambient temperature (18–23°C) and humidity (40–60%). Mice were provided standard mouse chow (Barastoc irradiated mouse cubes) and water *ad libitum.* All animal handling and procedures were conducted in a biosafety class 2 cabinet, in compliance with the University of Melbourne guidelines and approved by The University of Melbourne Animal Ethics Committee (application ID: 29017). Following acclimatisation, cages were randomly allocated to treatment or control groups. The *A. baumannii* inoculum for mice was prepared by inoculating Luria-Bertani broth with colonies from freshly sub-cultured bacteria on nutrient agar plates. The culture was grown at 37 °C with shaking to exponential phase (OD_600_= 0.8) and then washed with PBS, resuspended and diluted to the appropriate inoculation concentration in PBS with 5% (w/v) porcine mucin (Sigma-Aldrich). Mice were infected with 10^5^ CFU of *A. baumannii* strain A74 in 0.2 mL via intraperitoneal injection. One-hour post-infection, the animals were divided into four groups: two groups received 100 μg of Pse-specific mAb intravenously, one group received 100 μg of an isotype control mAb intravenously, and one group received PBS as untreated control. By 12 hours post-infection, all animal receiving PBS and those receiving the isotype control mAb reached the study’s ethical endpoints. These animals, along with one group of Pse mAb-treated mice, were humanely killed via CO_2_ asphyxiation. Blood was collected by cardiac puncture into EDTA tubes, and serially diluted in PBS before plating on nutrient agar for *A. baumannii* enumeration. The remaining Pse mAb-treated group was monitored for 7 days post-infection. At this time point, these animals were humanely killed, and their blood was collected and processed as described above to determine *A. baumannii* counts.

## Data Availability

Structure coordinates have been deposited in the Protein Data Bank (https://www.rcsb.org/) under accession code 9BEN. All mass spectrometry data (RAW files, FragPipe outputs, Rmarkdown scripts, and input tables) have been deposited into the PRIDE ProteomeXchange repository with the data identifiers: PXD056875 (Username, Password: 8cZfCmVbZFQK) for proteomic studies of *H. pylori* flagellin; PXD046836 (Username: Password: JkoHF9oP) for *H. pylori* P12 enrichments; PXD048493 (Username: Password: 8KOCxBaC) for *H. pylori* 26695 enrichments; PXD046858 (Username: Password: mmG9f4SP) for *C. jejuni* NCTC 11168 enrichments; PXD046859 (Username: Password: UsOXnSlC) for *A. baumannii* BAL062 enrichments; and PXD053904 (Username: Password: j689tJpTS3U0) for *A. baumannii* A74 enrichments.

## Acknowledgements

E.D.G.-B. acknowledges support from The Walter and Eliza Hall Institute of Medical Research; a Victorian State Government Operational Infrastructure support grant; the Brian M. Davis Charitable Foundation Centenary Fellowship; and NHMRC Ideas Grants GNT2027601 and GNT2000517. N.E.S is supported by an Australian Research Council (ARC) Future Fellowship (FT200100270), an ARC Discovery Project Grant (DP210100362) and a National Health and Medical Research Council Ideas grant (GNT2018980). R.J.P acknowledges funding from an NHMRC Investigator Grant (APP1174941). D.H.D. acknowledges funding from the National Institutes of Health award R15 GM109397. We would like to thank Johanna J. Kenyon (Griffith University) and Ruth M. Hall (University of Sydney) for providing clinical *Acinetobacter* strains and guidance on K types possessing different pseudaminic acid configurations/confirmations. We thank the Melbourne Mass Spectrometry and Proteomics Facility of The Bio21 Molecular Science and Biotechnology Institute for access to MS instrumentation.

## Author contributions

A.H.T., L.C., C.L., R. W. and X.L. planned and performed all chemical syntheses; A.De. and K.A.S. cultured *H. pylori* P12 samples; MK-L and LZ cultured *H. pylori* 26695 samples; M.C. and M.L. performed proteomic analyses of *H. pylori* P12 flagellin; N.M.S. raised/characterised mAbs, collected and analysed all SPR/BLI data, and performed all protein expression and structural biology; A.P.A., K.D.M. and D.H.D. made *H. pylori* Pse-knockout strains; A.Da. and S.J.C. cultured *C. jejuni* strains and purified flagellin from these samples; K.I.K. and N.E.S. cultured *A. baumannii* strains and performed *in vitro* infection assays; K.I.K., N.M.S. and N.E.S. performed glycopeptide enrichments and glycoproteomic analyses; L.L., G.P.C. and B.P.H. performed *in vivo* infection assays; N.E.S, E.D.G.-B. and R.J.P. conceived the project and co-wrote the manuscript.

## References

1 Varki, A., Schnaar, R. L. & Schauer, R. Sialic acids and other nonulosonic acids. In: Essentials of Glycobiology, Cold Spring Harbor (NY): Cold Spring Harbor Laboratory Press (2017).

2 McDonald, N. D. & Boyd, E. F. Structural and biosynthetic diversity of nonulosonic acids (NulOs) that decorate surface structures in bacteria. Trends Microbiol. 29, 142–157 (2021).

3 Logan, S. M., Kelly, J. F., Thibault, P., Ewing, C. P. & Guerry, P. Structural heterogeneity of carbohydrate modifications affects serospecificity of Campylobacter flagellins. Mol. Microbiol. 46, 587–597 (2002).

4 Lewis, A. L. et al. Innovations in host and microbial sialic acid biosynthesis revealed by phylogenomic prediction of nonulosonic acid structure. Proc. Natl. Acad. Sci. U. S. A. 106, 13552–13557 (2009).

5 de Jong, H., Wösten, M. M. & Wennekes, T. Sweet impersonators: Molecular mimicry of host glycans by bacteria. Glycobiology 32, 11–22 (2022).

6 Stephenson, H. N. et al. Pseudaminic acid on Campylobacter jejuni flagella modulates dendritic cell IL-10 expression via Siglec-10 receptor: a novel flagellin-host interaction. J. Infect. Dis. 210, 1487–1498 (2014).

7 Lee, I.-M., Wu, H.-Y., Angata, T. & Wu, S.-H. Bacterial pseudaminic acid binding to Siglec-10 induces a macrophage interleukin-10 response and suppresses phagocytosis. Chem. Commun. 60, 2930–2933 (2024).

8 Knirel, Y. A., Shashkov, A. S., Tsvetkov, Y. E., Jansson, P. E. & Zahringer, U. 5,7-Diamino-3,5,7,9-tetradeoxynon-2-ulosonic acids in bacterial glycopolymers: Chemistry and biochemistry. Adv. Carbohydr. Chem. Biochem., 58, 371–417 (2003).

9 Kiss, E. et al. The rkp-3 gene region of Sinorhizobium meliloti Rm41 contains strain-specific genes that determine K antigen structure. Mol. Plant Microbe Int. 14, 1395–1403 (2001).

10 Senchenkova, S. y. N., et al. Structure of a new pseudaminic acid-containing capsular polysaccharide of Acinetobacter baumannii LUH5550 having the KL42 capsule biosynthesis locus. Carbohydr. Res. 407, 154–157 (2015).

11 Knirel, Y. A. et al. Sialic acids of a new type from the lipopolysaccharides of *Pseudomonas aeruginosa* and *Shigella boydii*. Carbohydr. Res. 133, C5–C8 (1984).

12 Knirel, Y. A. et al. Somatic antigens of *Pseudomonas aeruginosa* - the structure of O-specific polysaccharide chains of the lipopolysaccharides from *Pseudomonas aeruginosa* O5 (Lanyi) and immunotype-6 (Fisher). Eur. J. Biochem. 163, 639–652 (1987).

13 Schirm, M. et al. Structural, genetic and functional characterization of the flagellin glycosylation process in *Helicobacter pylori*. Mol. Microbiol. 48, 1579–1592 (2003).

14 Thibault, P. et al. Identification of the carbohydrate moieties and glycosylation motifs in *Campylobacter jejuni* flagellin. J. Biol. Chem. 276, 34862–34870 (2001).

15 Posch, G. et al. Characterization and scope of S-layer protein O-glycosylation in *Tannerella forsythia*. J. Biol. Chem. 286, 38714–38724 (2011).

16 Horzempa, J., Dean, C. R., Goldberg, J. B. & Castric, P. Pseudomonas aeroginosa 1244 Pilin Glycosylation: Glycan Substrate Recognition. J. Bacteriol. 188, 4244–4252 (2006).

17 McNally, D. J. et al. Functional characterization of the flagellar glycosylation locus in *Campylobacter jejuni* 81-176 using a focused metabolomics approach. J. Biol. Chem. 281, 18489–18498 (2006).

18 Champasa, K., Longwell, S. A., Eldridge, A. M., Stemmler, E. A. & Dube, D. H. Targeted identification of glycosylated proteins in the gastric pathogen *Helicobacter pylori* (Hp). Mol. Cell. Proteomics 12, 2568–2586 (2013).

19 Hopf, P. S. et al. Protein glycosylation in *Helicobacter pylori*: beyond the flagellins? PLoS One 6, e25722 (2011).

20 Mahdavi, J. et al. A novel O-linked glycan modulates *Campylobacter jejuni* major outer membrane protein-mediated adhesion to human histo-blood group antigens and chicken colonization. Open Biol. 4, 130202 (2014).

21 Andolina, G. et al. Metabolic labeling of pseudaminic acid-containing glycans on bacterial surfaces. ACS Chem. Biol. 13, 3030–3037 (2018).

22 Wei, R. et al. Synthetic pseudaminic-acid-based antibacterial vaccine confers effective protection against *Acinetobacter baumannii* infection. ACS Cent. Sci. 7, 1535–1542 (2021).

23 Lee, I.-M. et al. Pseudaminic acid on exopolysaccharide of *Acinetobacter baumannii* plays a critical role in phage-assisted preparation of glycoconjugate vaccine with high antigenicity. J. Am. Chem. Soc. 140, 8639–8643 (2018).

24 Lee, I.-M., Tu, I.-F., Yang, F.-L. & Wu, S.-H. Bacteriophage tail-spike proteins enable detection of pseudaminic-acid-coated pathogenic bacteria and guide the development of antiglycan antibodies with cross-species antibacterial activity. J. Am. Chem. Soc. 142, 19446–19450 (2020).

25 Schirm, M. et al. Structural, genetic and functional characterization of the flagellin glycosylation process in *Helicobacter pylori*. Mol. Microbiol. 48, 1579–1592 (2003).

26 Wei, R., Liu, H., Tang, A. H., Payne, R. J. & Li, X. A Solution to Chemical Pseudaminylation via a Bimodal Glycosyl Donor for Highly Stereocontrolled α- and β-Glycosylation. Org. Lett. 21, 3584–3588 (2019).

27 Zhang, Y. et al. Enhanced Epimerization of Glycosylated Amino Acids During Solid-Phase Peptide Synthesis. J. Am. Chem. Soc. 134, 6316–6325 (2012).

28 Schoenhofen, I. C., McNally, D. J., Brisson, J. R. & Logan, S. M. Elucidation of the CMP-pseudaminic acid pathway in *Helicobacter pylori*: synthesis from UDP-N-acetylglucosamine by a single enzymatic reaction. Glycobiology 16, 8C–14C (2006).

29 Yang, K. Y. et al. Glycosyltransferase Jhp0106 (PseE) contributes to flagellin maturation in *Helicobacter pylori*. Helicobacter 26, e12787 (2021).

30 Castric, P., Cassels, F. J. & Carlson, R. W. Structural characterization of the *Pseudomonas aeruginosa* 1244 pilin glycan. J. Biol. Chem. 276, 26479–26485 (2001).

31 Champasa, K., Longwell, S. A., Eldridge, A. M., Stemmler, E. A. & Dube, D. H. Targeted identification of glycosylated proteins in the gastric pathogen Helicobacter pylori (Hp). Mol. Cell. Proteomics, 12, 2568–2586 (2013).

32 Ulasi, G. N., Creese, A. J., Hui, S. X., Penn, C. W. & Cooper, H. J. Comprehensive mapping of O-glycosylation in flagellin from *Campylobacter jejuni* 11168: A multienzyme differential ion mobility mass spectrometry approach. Proteomics 15, 2733–2745 (2015).

33 Zampronio, C. G., Blackwell, G., Penn, C. W. & Cooper, H. J. Novel glycosylation sites localized in *Campylobacter jejuni* flagellin FlaA by liquid chromatography electron capture dissociation tandem mass spectrometry. J. Proteome Res. 10, 1238–1245 (2011).

34 Thibault, P. et al. Identification of the carbohydrate moieties and glycosylation motifs in *Campylobacter jejuni* flagellin. J. Biol. Chem. 276, 34862–34870 (2001).

35 Mahdavi, J. et al. A novel O-linked glycan modulates *Campylobacter jejuni* major outer membrane protein-mediated adhesion to human histo-blood group antigens and chicken colonization. Open Biol. 4, 130202 (2014).

36 Kenyon, J. J. & Hall, R. M. Variation in the complex carbohydrate biosynthesis loci of *Acinetobacter baumannii* genomes. PloS One 8, e62160 (2013).

37 Lees-Miller, R. G. et al. A common pathway for O-linked protein-glycosylation and synthesis of capsule in *Acinetobacter baumannii*. Mol. Microbiol. 89, 816–830 (2013).

38 Iwashkiw, J. A. et al. Identification of a general O-linked protein glycosylation system in *Acinetobacter baumannii* and its role in virulence and biofilm formation. PLoS Pathog. 8, e1002758 (2012).

39 Nigro, S. J., Post, V. & Hall, R. M. Aminoglycoside resistance in multiply antibiotic-resistant *Acinetobacter baumannii* belonging to global clone 2 from Australian hospitals. J. Antimicrob. Chemother. 66, 1504–1509 (2011).

40 Kenyon, J. J., Marzaioli, A. M., Hall, R. M. & De Castro, C. Structure of the K2 capsule associated with the KL2 gene cluster of *Acinetobacter baumannii*. Glycobiology 24, 554–563 (2014).

41 Tkalec, K. I. et al. Glycan-tailored glycoproteomic analysis reveals serine is the sole residue subjected to O-linked glycosylation in *Acinetobacter baumannii*. J. Proteome Res. 23, 2474–2494 (2024).

42 Shashkov, A. S., et al. Characterisation of the carbapenem-resistant *Acinetobacter baumannii* clinical reference isolate BAL062 (CC2:KL58:OCL1): resistance properties and capsular polysaccharide structure. BioRxiv, 2024.2005.2009.593323, DOI: 10.1101/2024.05.09.593323 (2024).

43 Baker, S. et al. Exploiting human immune repertoire transgenic mice for protective monoclonal antibodies against antimicrobial resistant *Acinetobacter baumannii*. Nat. Commun. 15, 7979 (2024).

44 Yang, X. et al. Discovery of a monoclonal antibody that targets cell-surface pseudaminic acid of *Acinetobacter baumannii* with direct bactericidal effect. ACS Cent. Sci. 10, 439–446 (2024).

45 Nielsen, T. B. et al. Monoclonal antibody therapy against *Acinetobacter baumannii*. Infect. Immun. 89, e0016221 (2021).

46 Kenyon, J. J., Marzaioli, A. M., Hall, R. M. & De Castro, C. Structure of the K6 capsular polysaccharide from *Acinetobacter baumannii* isolate RBH4. Carbohydr. Res. 409, 30–35 (2015).

47 Kenyon, J. J. et al. The K46 and K5 capsular polysaccharides produced by *Acinetobacter baumannii* NIPH 329 and SDF have related structures and the side-chain non-ulosonic acids are 4-O-acetylated by phage-encoded O-acetyltransferases. PloS One 14, e0218461 (2019).

48 Nigro, S. J. & Hall, R. M. Loss and gain of aminoglycoside resistance in global clone 2 *Acinetobacter baumannii* in Australia via modification of genomic resistance islands and acquisition of plasmids. J. Antimicrob. Chemother. 71, 2432–2440 (2016).

49 Nielsen, T. B. et al. Monoclonal antibody requires immunomodulation for efficacy against *Acinetobacter baumannii* infection. J. Infect. Dis. 224, 2133–2147 (2021).

50 Russo, T. A. et al. The K1 capsular polysaccharide from *Acinetobacter baumannii* is a potential therapeutic target via passive immunization. Infect. Immun. 81, 915–922 (2013).

51 Schmitt, W. & Haas, R. Genetic analysis of the *Helicobacter pylori* vacuolating cytotoxin: structural similarities with the IgA protease type of exported protein. Mol. Microbiol. 12, 307–319 (1994).

52 Tomb, J. F. et al. The complete genome sequence of the gastric pathogen *Helicobacter pylori*. Nature 388, 539–547 (1997).

53 Parkhill, J. et al. The genome sequence of the food-borne pathogen *Campylobacter jejuni* reveals hypervariable sequences. Nature 403, 665–668 (2000).

54 Korlath, J. A., Osterholm, M. T., Judy, L. A., Forfang, J. C. & Robinson, R. A. A point-source outbreak of campylobacteriosis associated with consumption of raw milk. J. Infect. Dis. 152, 592–596 (1985).

55 Bouvet, P. J. M. & Grimont, P. A. D. Taxonomy of the Genus *Acinetobacter* with the Recognition of *Acinetobacter baumannii* sp. nov., *Acinetobacter haemolyticus* sp. nov., *Acinetobacter johnsonii* sp. nov., and *Acinetobacter junii* sp. nov. and emended descriptions of *Acinetobacter calcoaceticus* and *Acinetobacter lwoffii*. Int. J. Systematic Evol. Microbiol. 36, 228–240 (1986).

56 Nemec, A., Janda, L., Melter, O. & Dijkshoorn, L. Genotypic and phenotypic similarity of multiresistant *Acinetobacter baumannii* isolates in the Czech Republic. J. Med. Microbiol. 48, 287–296 (1999).

57 Nhu, N. T. K. et al. Emergence of carbapenem-resistant *Acinetobacter baumannii* as the major cause of ventilator-associated pneumonia in intensive care unit patients at an infectious disease hospital in southern Vietnam. J. Med. Microbiol. 63, 1386–1394 (2014).

58 Logan, S. M. & Trust, T. J. Molecular identification of surface protein antigens of *Campylobacter jejuni*. Infect. Immun. 42, 675–682 (1983).

59 Suerbaum, S., Geis, G., Josenhans, C. & Opferkuch, W. Biochemical studies of *Helicobacter mustelae* fatty acid composition and flagella. Infect. Immun. 60, 1695–1698 (1992).

60 Harney, D. J. et al. Proteomics analysis of adipose depots after intermittent fasting reveals visceral fat preservation mechanisms. Cell Rep. 34, 108804 (2021).

61 Harney, D. J. et al. Proteomic analysis of human plasma during intermittent fasting. J. Proteome Res. 18, 2228–2240 (2019).

62 Saba, J., Dutta, S., Hemenway, E. & Viner, R. Increasing the productivity of glycopeptides analysis by using higher-energy collision dissociation-accurate mass-product-dependent electron transfer dissociation. Int. J. Proteomics 2012, 560391 (2012).

63 Schmidt, T. et al. Universal spectrum explorer: A standalone (web-)application for cross-resource spectrum comparison. J. Proteome Res. 20, 3388–3394 (2021).

64 Kabsch, W. XDS. Acta Crystallogr. D Biol. Crystallogr. 66, 125–132 (2010).

65 Storoni, L. C., McCoy, A. J. & Read, R. J. Likelihood-enhanced fast rotation functions. Acta Crystallogr. D Biol. Crystallogr. 60, 432–438 (2004).

66 Emsley, P. & Cowtan, K. Coot: model-building tools for molecular graphics. Acta Crystallogr. D Biol. Crystallogr. 60, 2126–2132 (2004).

67 Adams, P. D. et al. PHENIX: a comprehensive Python-based system for macromolecular structure solution. Acta Crystallogr. D Biol. Crystallogr. 66, 213–221 (2010).

68 Goddard, T. D. et al. UCSF ChimeraX: Meeting modern challenges in visualization and analysis. Protein Sci. 27, 14–25 (2018).

69 Gibson, D. G. et al. Enzymatic assembly of DNA molecules up to several hundred kilobases. Nat. Methods 6, 343–345 (2009).

70 Salama, N. R., Shepherd, B. & Falkow, S. Global transposon mutagenesis and essential gene analysis of *Helicobacter pylori*. J. Bacteriol. 186, 7926–7935 (2004).

71 Rappsilber, J., Mann, M. & Ishihama, Y. Protocol for micro-purification, enrichment, pre-fractionation and storage of peptides for proteomics using StageTips. Nat. Protoc. 2, 1896–1906 (2007).

72 Rappsilber, J., Ishihama, Y. & Mann, M. Stop and go extraction tips for matrix-assisted laser desorption/ionization, nanoelectrospray, and LC/MS sample pretreatment in proteomics. Anal. Chem. 75, 663–670 (2003).

73 Udeshi, N. D., Mertins, P., Svinkina, T. & Carr, S. A. Large-scale identification of ubiquitination sites by mass spectrometry. Nat. Protoc. 8, 1950–1960 (2013).

74 Udeshi, N. D. et al. Refined preparation and use of anti-diglycine remnant (K-epsilon-GG) antibody enables routine quantification of 10,000s of ubiquitination sites in single proteomics experiments. Mol. Cell. Proteomics, 12, 825–831 (2013).

75 Caval, T., Zhu, J. & Heck, A. J. R. Simply extending the mass range in electron transfer higher energy collisional dissociation increases confidence in N-glycopeptide identification. Anal. Chem. 91, 10401–10406 (2019).

76 Teo, G. C., Polasky, D. A., Yu, F. & Nesvizhskii, A. I. Fast deisotoping algorithm and its implementation in the MSFragger search engine. J. Proteome Res. 20, 498–505 (2021).

77 da Veiga Leprevost, F., et al. Philosopher: a versatile toolkit for shotgun proteomics data analysis. Nat. Methods 17, 869–870 (2020).

78 Geiszler, D. J. et al. PTM-Shepherd: Analysis and summarization of post-translational and chemical modifications from open search results. Mol. Cell. Proteomics 20, 100018 (2021).

79 Kong, A. T., Leprevost, F. V., Avtonomov, D. M., Mellacheruvu, D. & Nesvizhskii, A. I. MSFragger: ultrafast and comprehensive peptide identification in mass spectrometry-based proteomics. Nat. Methods 14, 513–520 (2017).

80 Yu, F. et al. Identification of modified peptides using localization-aware open search. Nat. Commun. 11, 4065 (2020).

81 Lewis, J. M., Coulon, P. M. L., McDaniels, T. A. & Scott, N. E. The application of open searching-based approaches for the identification of *Acinetobacter baumannii* O-linked glycopeptides. J. Vis. Exp. (2021).

82 Cox, J. & Mann, M. MaxQuant enables high peptide identification rates, individualized p.p.b.-range mass accuracies and proteome-wide protein quantification. Nat. Biotechnol. 26, 1367–1372 (2008).

83 Brademan, D. R., Riley, N. M., Kwiecien, N. W. & Coon, J. J. Interactive peptide spectral annotator: a versatile web-based tool for proteomic applications. Mol. Cell. Proteomics, 18, S193–S201 (2019).

84 Perez-Riverol, Y. et al. The PRIDE database and related tools and resources in 2019: improving support for quantification data. Nucleic Acids Res. 47, D442–D450 (2019).

85 Vizcaino, J. A. et al. 2016 update of the PRIDE database and its related tools. Nucleic Acids Res. 44, D447–456 (2016).

86 Talyansky, Y. et al. Capsule carbohydrate structure determines virulence in *Acinetobacter baumannii*. PLoS Pathog. 17, e1009291 (2021).

87 Schindelin, J., et al. Fiji: an open-source platform for biological-image analysis. Nat. Methods 9, 676–682 (2012).

