## Supplementary document 1 for "Uncovering bacterial pseudaminylation with pan-specific antibody tools"

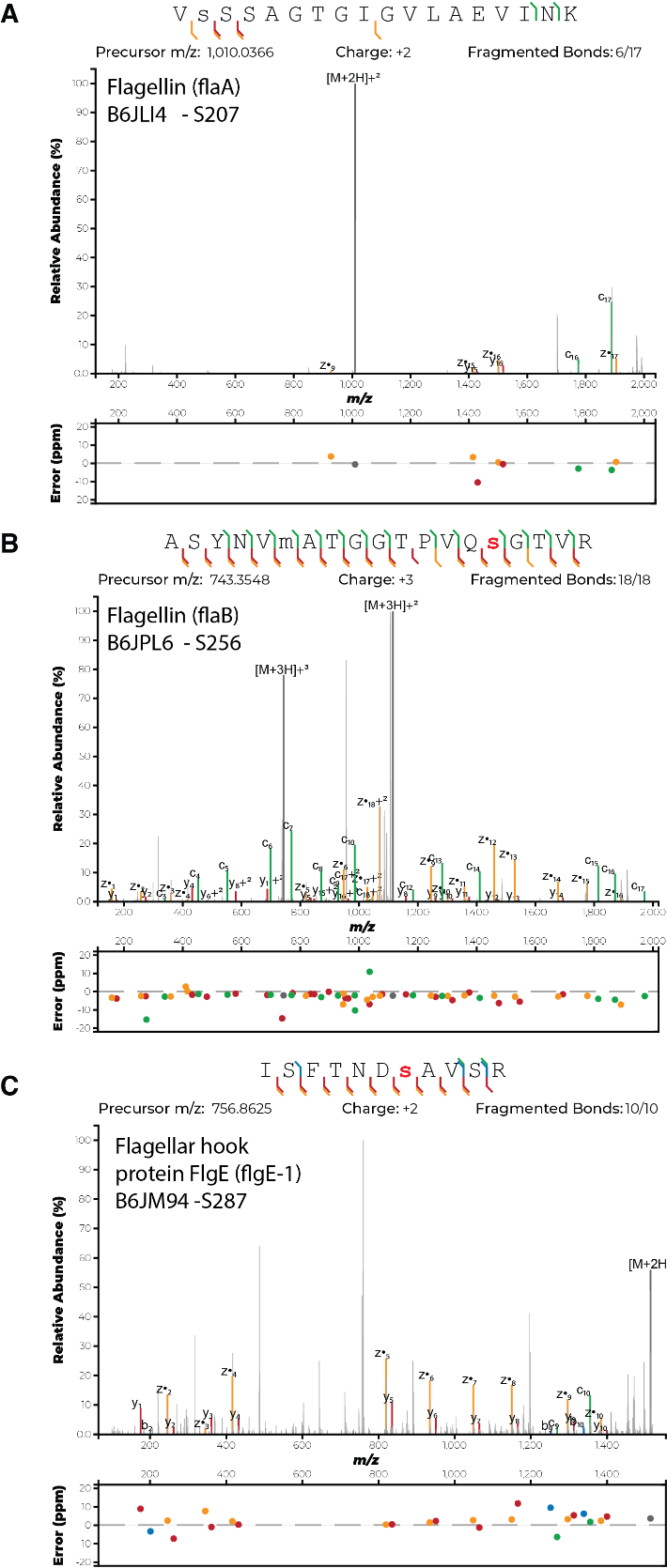

**Extended Data Figure 1. ExD enabled localisation of native *H. pylori* glycopeptides FlaA (206-211), FlaB (242-260) and FlgE(281-291).** Localised pseudaminylated events within tryptic digests of flagellin preparation of *H. pylori* P12 for A) FlaA ^206^VSSSAGTGIGVLAEVINK^211^, B) FlaB ^242^ASYNVMATGGTPVQSGTVR^260^ and C) FlgE ^281^ISFTNDSAVSR^291^. For the FlgE glycopeptide ^281^ISFTNDSAVSR^291^ PRM EAD analysis was undertaken to allow localisation of glycosylation events while EThcD was undertaken on ^206^VSSSAGTGIGVLAEVINK^211^ and ^242^ASYNVMATGGTPVQSGTVR^260^.

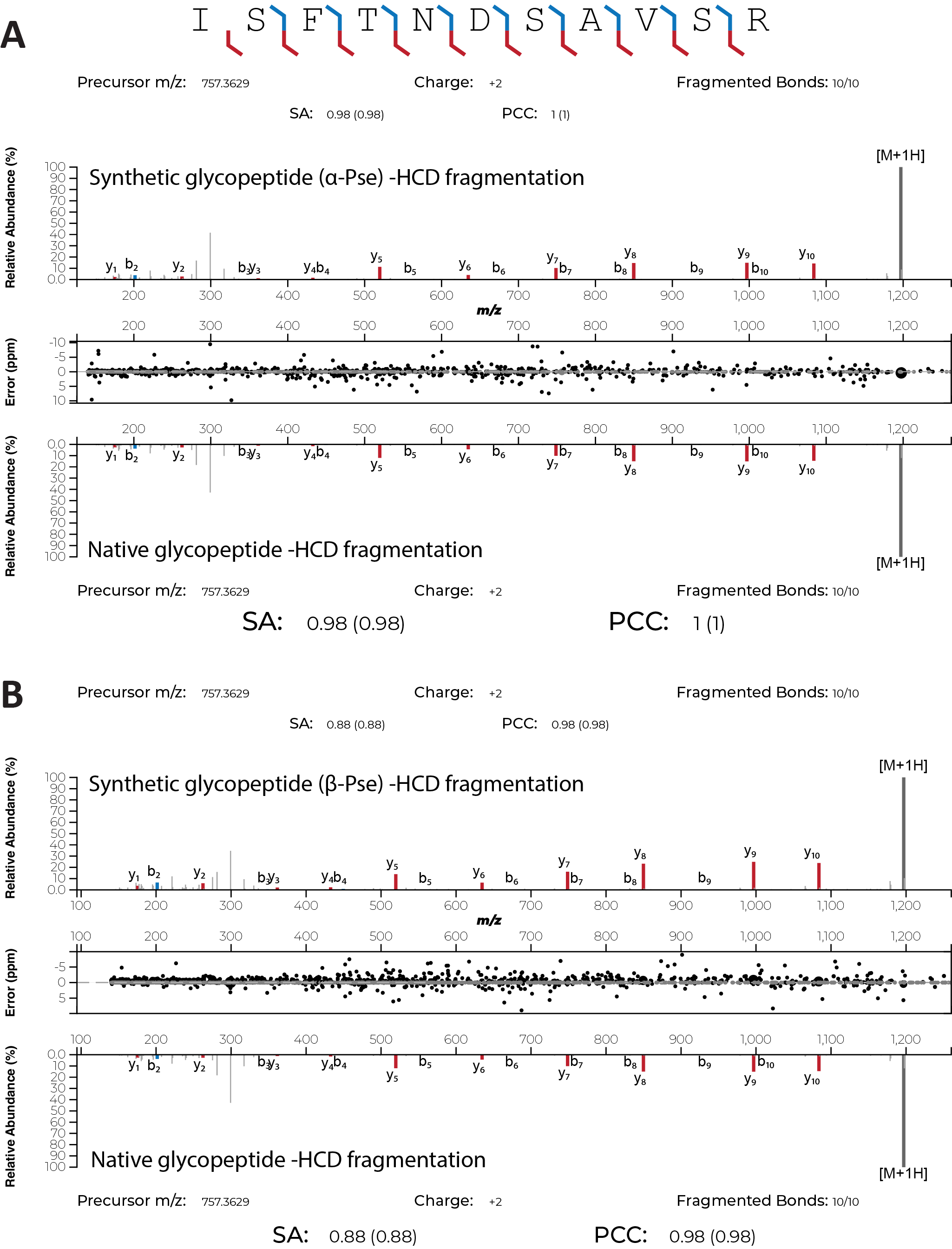

**Extended Data Figure 2. Butterfly plots of synthetic verses native glycopeptide ^281^ISFTNDSAVSR^291^ of FlgE.** Butterfly plots comparing HCD fragmentation of synthetic glycopeptides bearing an α- or β-configured Pse residue with the identical authentic glycopeptides derived from tryptic digests of flagellin preparation of *H. pylori* P12. Spectral angle (SA) analysis comparing matched b- and y-ion intensities to native ^281^ISFTNDSAVSR^291^ of FlgE supports highest similarity to α-pseudaminylation compared to β-pseudaminylation.

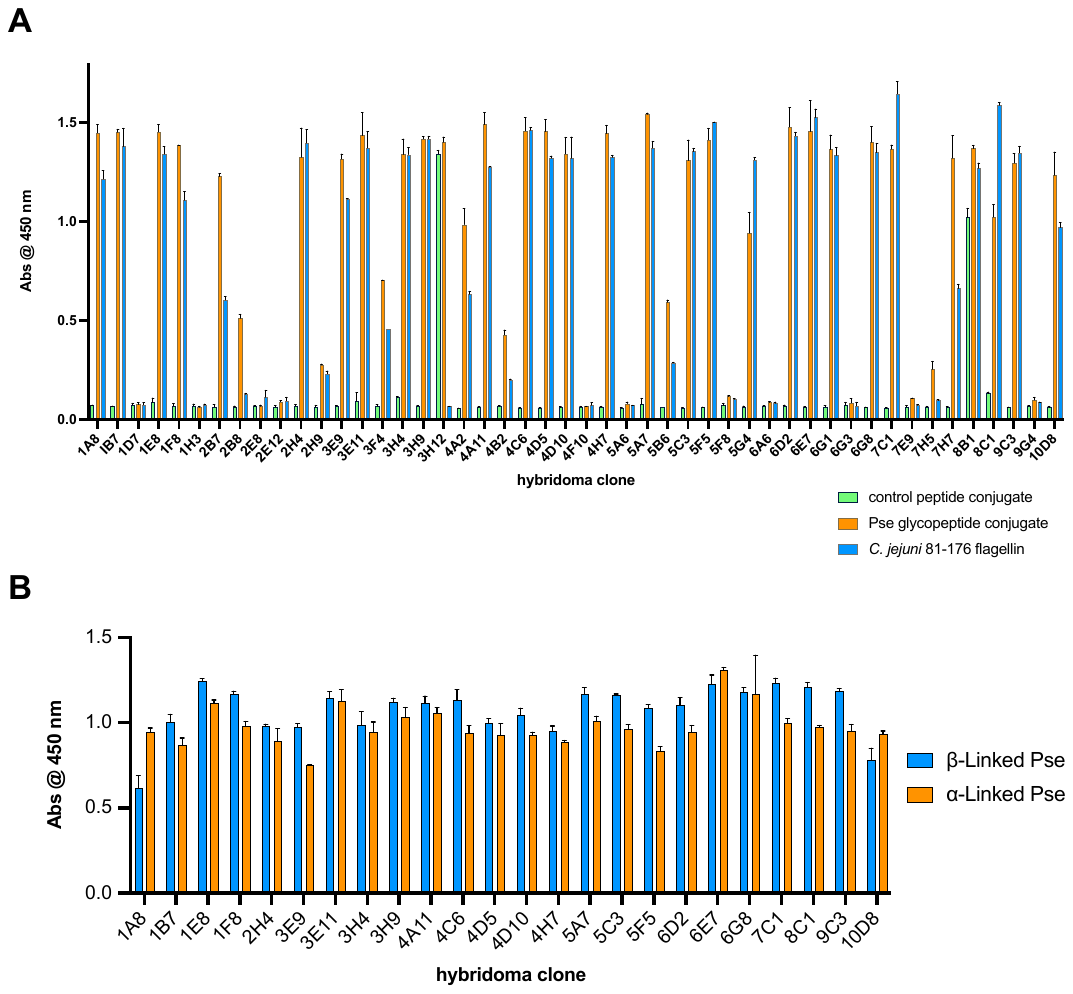

**Extended Data Figure 3. ELISA Screening of potential Pse specific hybridoma supernatant.** (A) ELISA screen of hybridoma supernatant (1:1000) against: (i) control peptides **8**, **9** and **10** conjugated to BSA (green), (ii) Pse peptides **α-8**, **α-9** and **α-10** conjugated to BSA (orange), and flagellin purified from *C. jejuni* 81-176, which is also pseudaminylated (blue). (B) ELISA screen of hybridoma supernatants (1:2000) that are Pse-specific against: (i) Pse peptides **α-8**, **α-9** and **α-10** conjugated to BSA (orange) and (ii) Pse peptides **β-8**, **β-9** and **β-10** conjugated to BSA (blue).

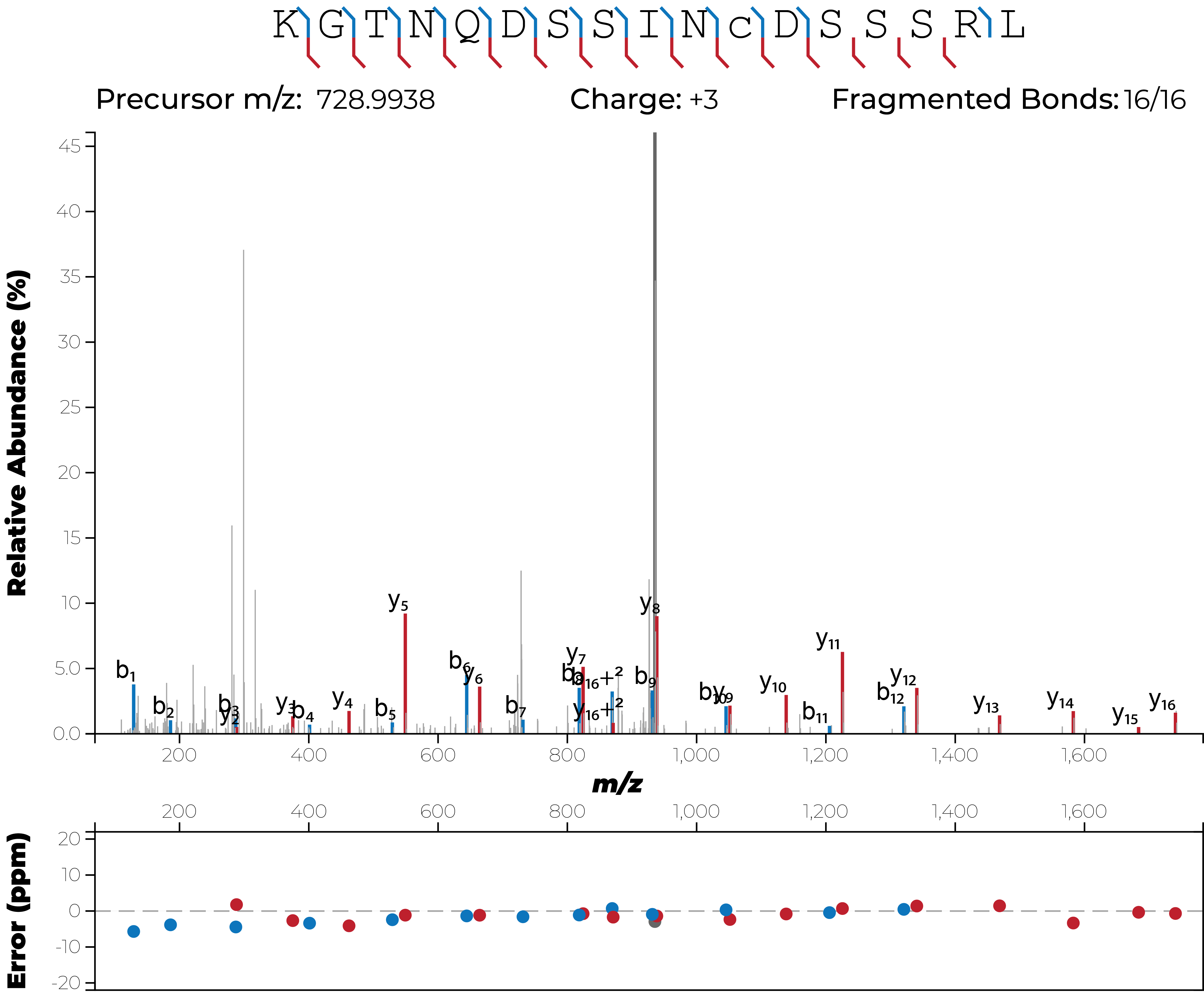

**Extended Data Figure 4.** HCD spectra of the *H. pylori* HPP12_0570 glycopeptide ^65^KGTNQDSSINCDSSSRL^82^.

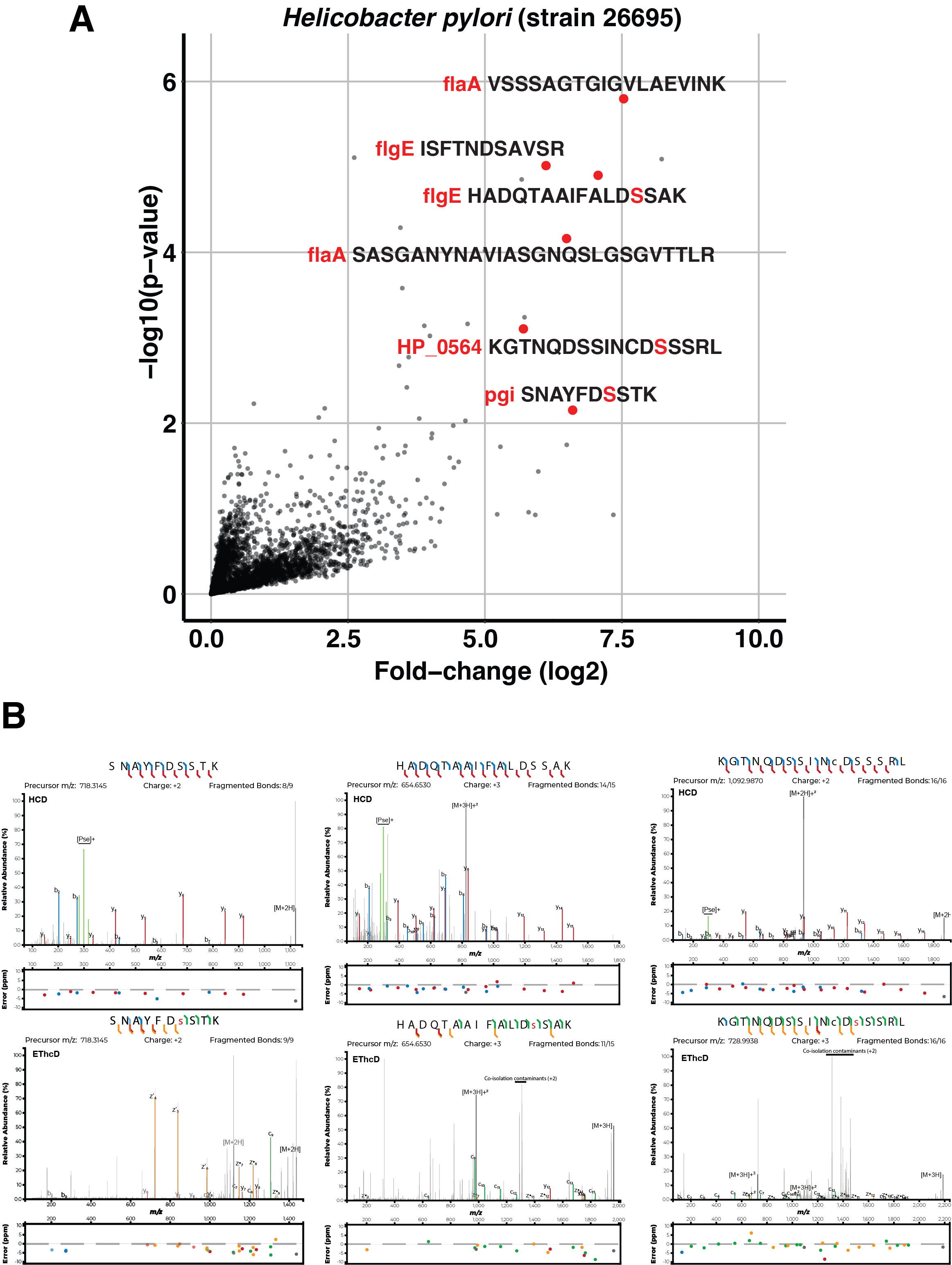

**Extended Data Figure 5. Pse-enrichment of *H. pylori* 26695.** (A) Volcano plot illustrating the enrichment of Pse5Ac7Ac-modified glycopeptides from *H. pylori* 26695. (B) EThcD / HCD spectra of the novel glycopeptides derived from Pgi, FlgE and HP_0564.

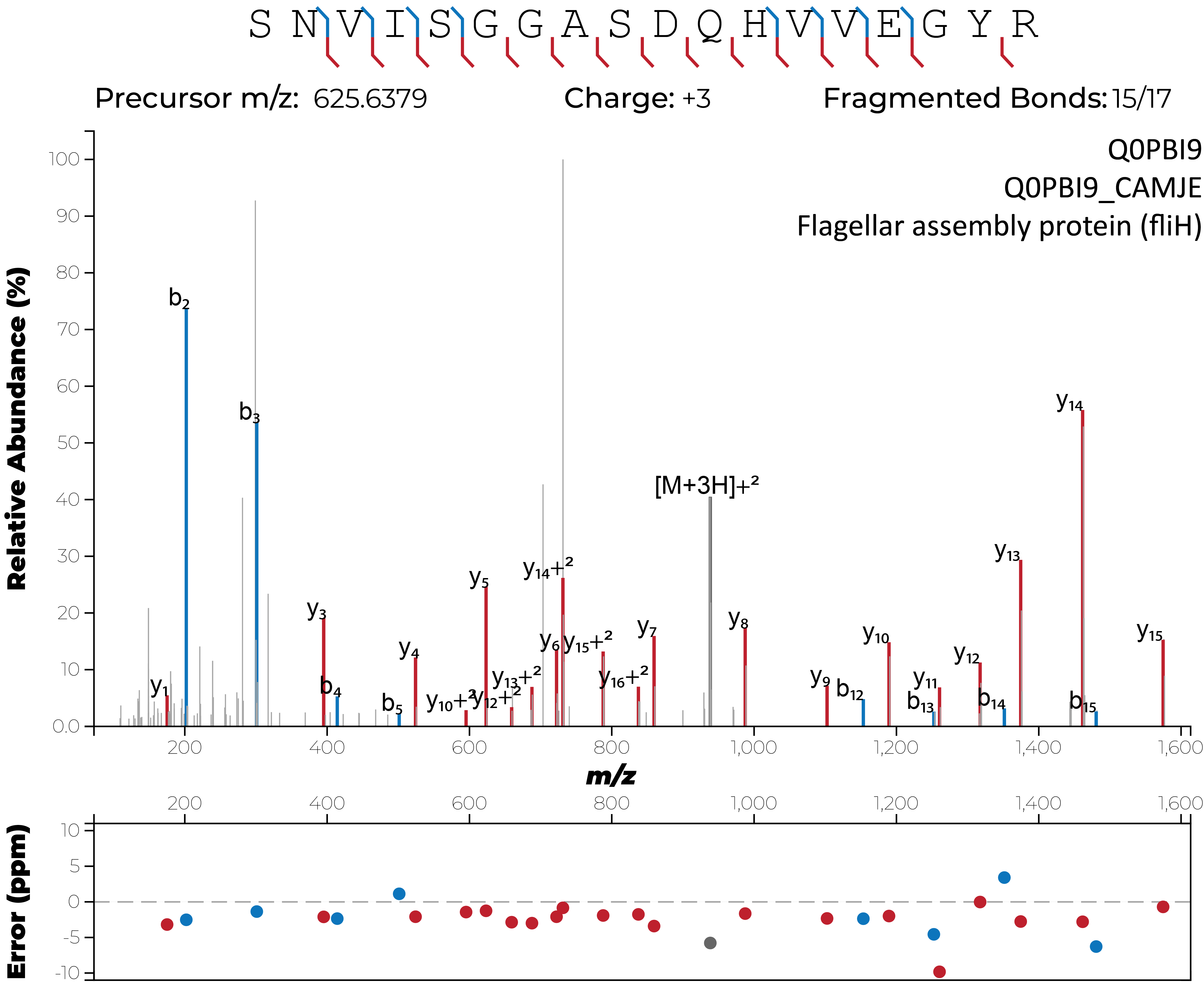

**Extended Data Figure 6.** HCD spectra of the *C. jejuni* FliH glycopeptide ^5^SNVISGGASDQHVVEGYR^22^.

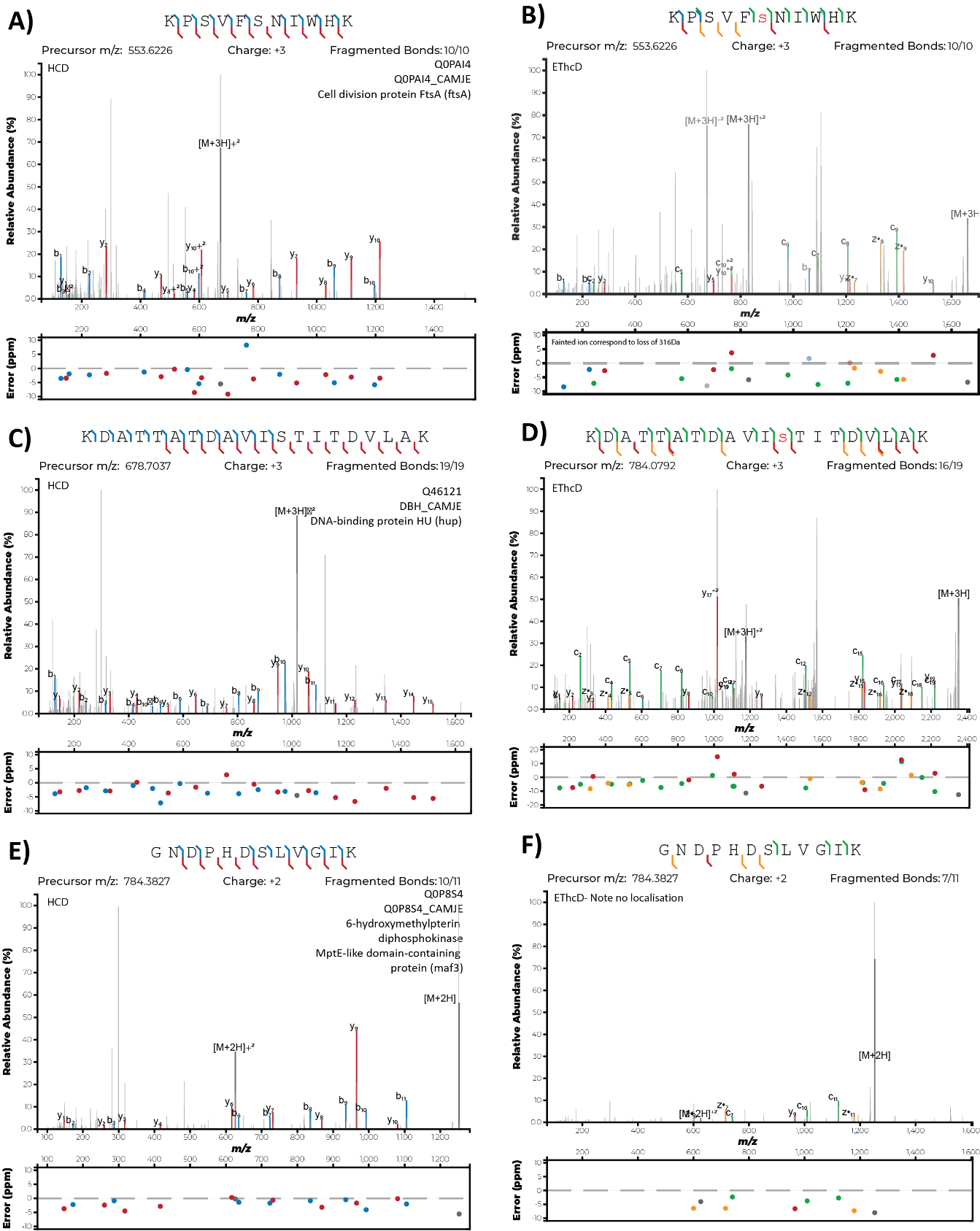

**Extended Data Figure 7. Pseudaminylation of *C. jejuni* glycoproteins.** MS/MS EThcD and HCD spectra of **A / B)** ^447^KPSVFSNIWHK^457^ from FtsA (Q0PAI4_CAMJE) with pseudaminylation localised to Ser452; **C / D)** ^19^KDATTATDAVISTITDVLAK^38^ from Hup (Q46121_CAMJE) with pseudaminylation localised to Ser30 and **E / F)** ^191^GNDPHDSLVGIK^202^ from Maf3 (Q0P8S4_CAMJE) with pseudaminylation inferred to Ser197 by the absence of other hydroxyl containing amino acids.

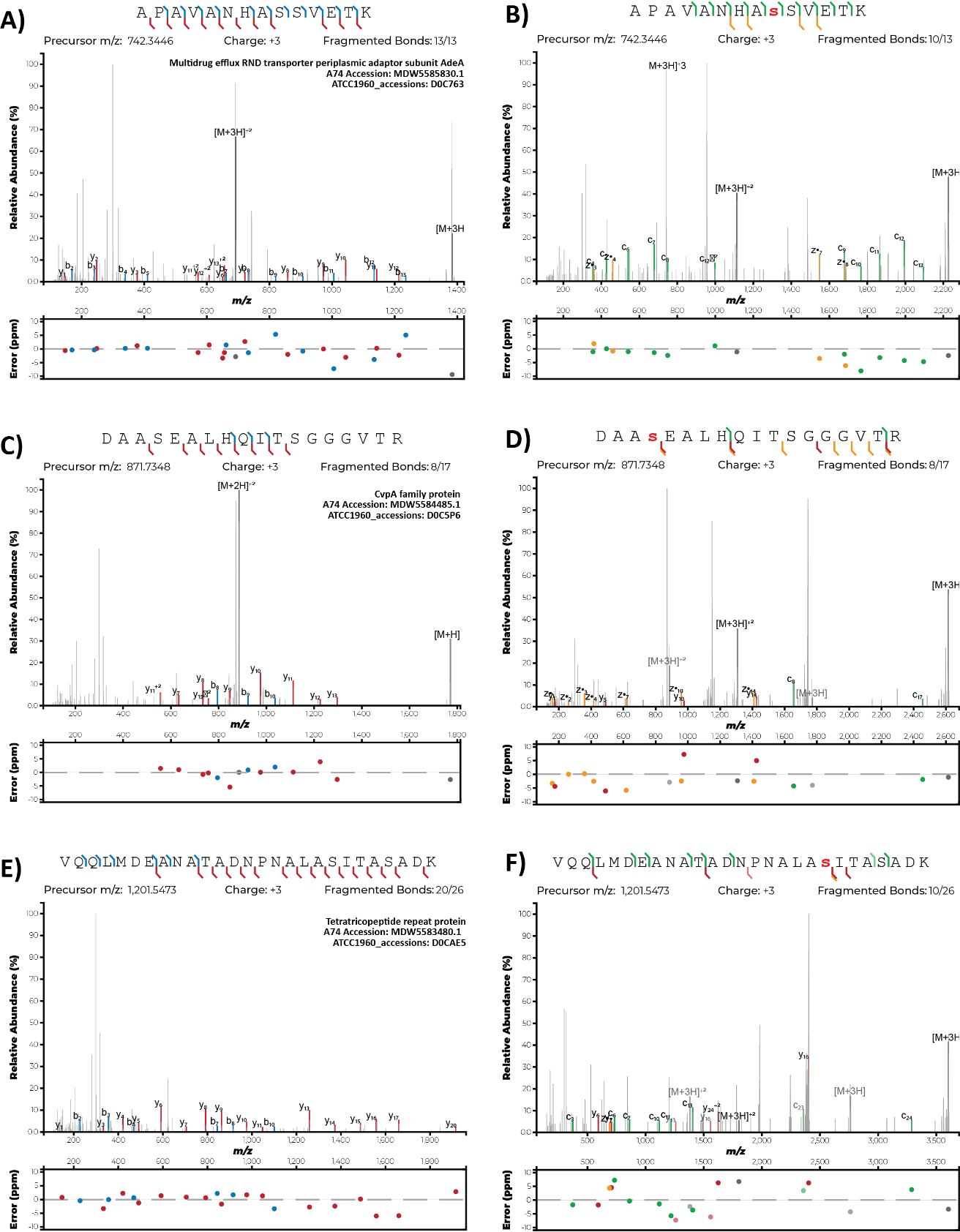

**Extended Data Figure 8. Novel Pseudaminic acid-containing glycopeptides identified in *A. baumannii* A74.** MS/MS EThcD and HCD spectra of unique the glycopeptide identified within *A. baumannii* A74 corresponding to A/B) ^374^APAVANHASSVETK^387^ of the multidrug efflux RND transporter periplasmic adaptor subunit AdeA (MDW5585830.1); **C / D)** ^153^DAASEALHQITSGGGVTR^170^ of the CvpA family protein (MDW5584485.1) and **E / F)** ^55^VQQLMDEANATADNPNALASITASADK^81^ of the tetratricopeptide repeat protein (MDW5583480.1).

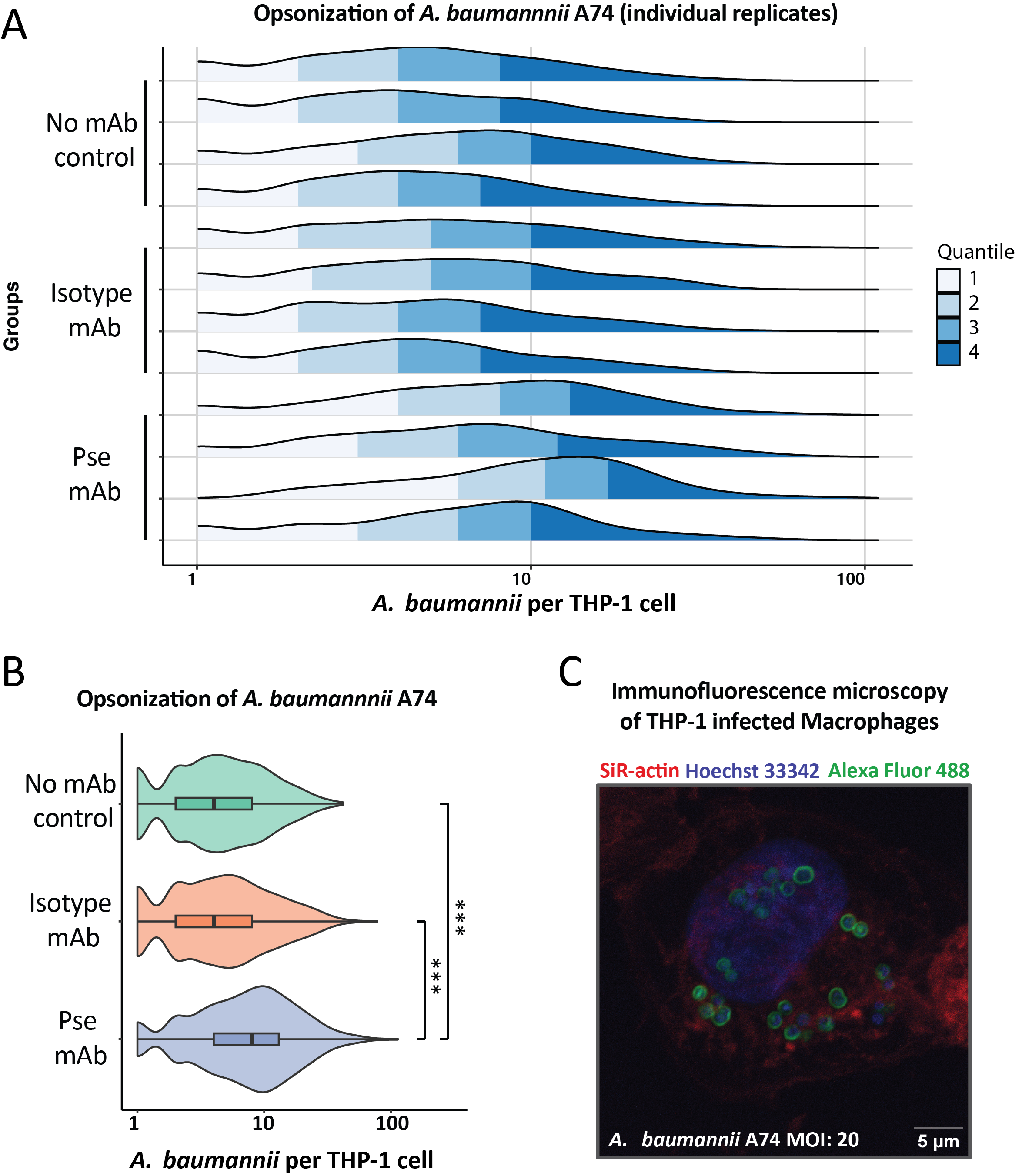

**Extended Data Figure 9. Immunofluorescence based opsonization assays of *A. baumannii* A74. A)** Density plots of biological replicates assessing the intracellular number of *A. baumannii* A74 in THP-1 macrophages treated with in the absence of added antibody, or in the presence of Pse 1E8 or a mouse IgG1 isotype control. **B)** Violin plots depicting the abundance of intracellular bacteria in THP-1 macrophages treated with *A. baumannii* A74 in the absence of added antibody, or in the presence of Pse 1E8 or a mouse IgG1 isotype control. **C)** A representative immunofluorescence microscopy micrograph of THP-1 cells one hour after infection with *A. baumannii* A74. DNA has been stained with Hoechst 33342 (blue), actin with SiR-Actin (red), and Pse mAb 1E8 (green). Scale bar is 5 μm.

**Supplementary Tables**

**Supplementary Table 1. Pseudaminic acid containing glycopeptides from *H. pylori* P12 identified from flagella preparations.** Combined modified peptide MSfragger searches from *H. pylori* P12 flagella preparations (n=3). For all identified peptides the ion intensity observed across replicates, modification status of the modified peptide, spectral counts of the modified peptide, position of first amino acid of the peptide within the protein, position of the last amino acid of the peptide within the protein, peptide length, MaxLFQ intensities of the observed modified peptides, first amino acid in the peptide, last amino acid in the peptide, Uniprot accession number, gene name and protein.ID are provided. Glycopeptides were manually assessed to confirm sites of localisation and correctness, with HCD and EThcD spectra of the best scoring validated glycopeptides provided in Supplementary Data Table1.

| **Structure** | **scFv-1E8(b):Pse-Ser complex**  **(PDB ID: 9BEN)** |
| --- | --- |
| **Data collection** |  |
| Space group | P 65 2 2 |
| No. of protein chains in the AU | 1 |
| Cell dimensions |  |
| *a*, *b*, *c* (Å) | 67.928 67.928 225.89 |
| α, β, γ (°) | 90 90 120 |
| Wavelength (Å) | 0.9537 |
| Resolution (Å)* | 46.36 - 2.063 (2.137 - 2.063) |
| *R_merge_** | 0.1311 (1.398) |
| *R*_pim_* | 0.02139 (0.2238) |
| *I /* σ*I** | 25.17 (3.04) |
| CC(1/2)* | 1 (0.89) |
| Completeness (%)* | 99.71 (97.30) |
| Redundancy* | 38.4 (38.7) |
| Wilson B-factor (Å^2^) | 35.75 |
| **Refinement** |  |
| Resolution (Å) | 46.36 - 2.063 |
| No. reflections* | 19894 (1873) |
| *R*_work_ / *R*_free_ | 0.1929/ 0.2286 |
| No. non-hydrogen atoms | 1917 |
| Protein | 1736 |
| Ligands | 79 |
| Water | 102 |
| *B*-factors | 37.47 |
| Protein | 36.69 |
| Ligands | 48.65 |
| Water | 42.05 |
| R.m.s. deviations |  |
| Bond lengths (Å) | 0.006 |
| Bond angle (°) | 0.82 |
| Ramachandran plot (%) |  |
| Favored | 98.24 |
| Allowed | 1.76 |
| Disallowed | 0.00 |

* Values in parentheses indicate values for highest resolution shell

**Supplementary Table 2.** Data collection and refinement statistics for the scFv1E8b:Pse-Ser complex.

**Supplementary Table 3. Pseudaminic acid containing glycopeptide enrichments of *H. pylori P12*.** Combined modified peptide MSfragger searches from *H. pylori* P12 enrichments across biological replicates (n=5). For all identified peptides the ion intensity observed across replicates, modification status of the modified peptide, spectral counts of the modified peptide, position of first amino acid of the peptide within the protein, position of the last amino acid of the peptide within the protein, peptide length, MaxLFQ intensities of the observed modified peptides, Perseus generated statistical outputs comparing peptide levels between groups, first amino acid in the peptide, last amino acid in the peptide, Uniprot accession number, gene name and protein.ID are provided.

**Supplementary Table 4. Pseudaminic acid containing glycopeptide enrichments of *H. pylori* 26695.** Combined modified peptide MSfragger searches from *H. pylori* 26695 enrichments across biological replicates (n=5). For all identified peptides the ion intensity observed across replicates, modification status of the modified peptide, spectral counts of the modified peptide, position of first amino acid of the peptide within the protein, position of the last amino acid of the peptide within the protein, peptide length, MaxLFQ intensities of the observed modified peptides, Perseus generated statistical outputs comparing peptide levels between groups, first amino acid in the peptide, last amino acid in the peptide, Uniprot accession number, gene name and protein.ID are provided.

**Supplementary Table 5. Pseudaminic acid containing glycopeptide enrichments of *C. jejuni* NCTC11168.** Combined modified peptide MSfragger searches from *C. jejuni* NCTC11168 enrichments across biological replicates (n=3). For all identified peptides the ion intensity observed across replicates, modification status of the modified peptide, spectral counts of the modified peptide, position of first amino acid of the peptide within the protein, position of the last amino acid of the peptide within the protein, Peptide length, MaxLFQ intensities of the observed modified peptides, Perseus generated statistical outputs comparing peptide levels between groups, first amino acid in the peptide, last amino acid in the peptide, Uniprot accession number, gene name and protein.ID are provided.

**Supplementary Table 6. Pseudaminic acid containing glycopeptide enrichments of *A. baumannii* A74.** Open searching and combined modified peptide MSfragger search for *A. baumannii* A74 enrichments across biological replicates (n=5). For the open searches all identified spectra, data files, peptide sequence, peptide length, charge, retention times, observed m/z, calibrated m/z, observed mass, calibrated mass, delta mass, scores, peptide position within protein, assigned modifications, protein, protein.ID and replicate number are provided. For the combined modified peptide search, the ion intensity observed across replicates, modification status of the modified peptide, spectral counts of the modified peptide, position of first amino acid of the peptide within the protein, position of the last amino acid of the peptide within the protein, peptide length, MaxLFQ intensities of the observed modified peptides, Perseus generated statistical outputs comparing peptide levels between groups, first amino acid in the peptide, last amino acid in the peptide, NCBI accession number, gene name and protein.ID are provided for all observed peptides.

**Supplementary Table 7. Pseudaminic acid containing glycopeptide enrichments of *A. baumannii* BAL062.** Open searching and combined modified peptide MSfragger search for *A. baumannii* BAL062 enrichments across biological replicates (n=5). For the open searches all identified spectra, data files, peptide sequence, peptide length, charge, retention times, observed m/z, calibrated m/z, observed mass, calibrated mass, delta mass, scores, peptide position within protein, assigned modifications, protein, protein.ID and replicate number are provided. For the combined modified peptide search, the ion intensity observed across replicates, modification status of the modified peptide, spectral counts of the modified peptide, position of first amino acid of the peptide within the protein, position of the last amino acid of the peptide within the protein, peptide length, MaxLFQ intensities of the observed modified peptides, Perseus generated statistical outputs associated comparing peptide levels between groups, first amino acid in the peptide, last amino acid in the peptide, NCBI accession number, gene name and protein.ID are provided for all observed peptides.

**Supplementary Table 8.** Nucleotide and protein sequences for Fab and scFv.

| protein | dsDNA oligo | protein sequence |
| --- | --- | --- |
| Fab 1E8 | gatatacatATGaaaaagaatatcgcatttcttcttgcatctatgttcgttttttctattgctacaaatgcctatgcatccGACATCCAAATGATTCAGAGCCCATCCTCCATGTTCGCTAGCCTGGGTGACCGTGTATCTCTGTCTTGCCGCGCAAGCCAGGGCATCCGTGGCAACCTGGACTGGTATCAGCAGAAACCGGGTGGCCCTATTAAACTGCTGATCCACAGCACCTCTAAGCTGAACTCCGGTGTTCCGTCTCGTTTCTCCGGTTCCGGCTCTGGTAGCGATTACTCTCTGACCATTTCCTCCCTGGAATCTGAAGATTTCGCAGACTATTATTGCCTGCAGCGTAATGCTTTCCCACTGACCTTTGGCGCGGGTACCAAACTGGAACTGAAACGTGCGGACGCAGCACCTACTGTCAGCATCTTCCCGCCGTCTAGCGAACAACTGACCTCTGGTGGTGCGTCTGTTGTTTGCTTTCTGAACAACTTCTACCCGAAGGACATCAACGTCAAGTGGAAAATTGACGGCTCTGAACGCCAGAACGGCGTTCTGAATTCTTGGACCGACCAGGATTCCAAGGATTCCACCTATTCTATGAGCTCCACTCTGACGCTGACCAAAGACGAATACGAGCGTCACAACAGCTATACCTGTGAAGCAACCCATAAAACCTCCACCTCCCCAATTGTGAAGTCCTTCAACCGTAACGAGTGCTAAttaactcgaggctgagcaaagcagactactaataacataaagtctacgccggacgcatcgtggccctagtacgcaagttcacgtaaaaagggtaactagaggttgaggtgattttatgaaaaagaatatcgcatttcttcttgcatctatgttcgttttttctattgctacaaacgcgtacgctgagatctccGAAGTGAAACTGTTTCAGTCTGGCGGTGGTCTGGTTCAGCCAGGTGGTTCTCTGAAACTGTCCTGCGCTGCATCCGGTATTGATTTCTCCCGCAATTGGATGACCTGGGTACGTCGTGCGCCGGGTAAAGGTCTGGAATGGATCGGTGAAATCTACCCGGATTCCCGTACCATTAACTACGCACCGAGCCTGAAAGACAAGTTTATCATCTCTCGCGACAACGCGAAAAAAATGCTGTACCTGCAGATGTCCAAGGTTCGTAGCGAAGATACCGCCCTGTATTATTGCGCACGTCGTGGTGTTACTACCGTAGTAGGTACCAGCTGGTACTTCGATGTGTGGGGCACCGGTACCACTGTAACCGTTAGCTCCGCGAAAACCACTCCGCCGTCTGTGTACCCACTGGCACCTGGTAGCGCTGCTCAGACCAACTCTATGGTTACGCTGGGTTGTCTGGTAAAAGGCTACTTCCCGGAACCGGTAACTGTCACCTGGAACAGCGGCTCCCTGTCTAGCGGTGTTCACACTTTCCCGGCGGTTCTGCAGTCCGACCTGTACACCCTGTCTTCTAGCGTCACCGTTCCTTCCAGCACCTGGCCATCTCAGACCGTTACTTGCAACGTAGCCCACCCAGCCTCCTCCACCAAAGTTGATAAAAAAATCGTTCCGCGTGATTGTGGCGCAAAACCGtctagacaccaccaccaccaccactaactcgagaattca | MKKNIAFLLASMFVFSIATNAYASDIQMIQSPSSMFASLGDRVSLSCRASQGIRGNLDWYQQKPGGPIKLLIHSTSKLNSGVPSRFSGSGSGSDYSLTISSLESEDFADYYCLQRNAFPLTFGAGTKLELKRADAAPTVSIFPPSSEQLTSGGASVVCFLNNFYPKDINVKWKIDGSERQNGVLNSWTDQDSKDSTYSMSSTLTLTKDEYERHNSYTCEATHKTSTSPIVKSFNRNEC  MKKNIAFLLASMFVFSIATNAYAEISEVKLFQSGGGLVQPGGSLKLSCAASGIDFSRNWMTWVRRAPGKGLEWIGEIYPDSRTINYAPSLKDKFIISRDNAKKMLYLQMSKVRSEDTALYYCARRGVTTVVGTSWYFDVWGTGTTVTVSSAKTTPPSVYPLAPGSAAQTNSMVTLGCLVKGYFPEPVTVTWNSGSLSSGVHTFPAVLQSDLYTLSSSVTVPSSTWPSQTVTCNVAHPASSTKVDKKIVPRDCGAKPSRHHHHHH |
| Fab 3E11 | gatatacatATGaaaaagaatatcgcatttcttcttgcatctatgttcgttttttctattgctacaaatgcctatgcatccGATATTCAGCTGATTCAGTCCCCGTCCTCTATTTTTGCATCTCTGGGTGACCGCGTCTCTCTGTCCTGTCGTGCGTCCCAGGGCATTCGTGGCAACCTGGATTGGTATCAACAGAAGCCTGGTGGTACTATCAAACTGCTGATCTACTCTACCAGCAACCTGAAAAGCGGTGTACCATCCCGTTTCTCCGGTCGTGGTAGCGGCTCTGACTATTCTCTGACCATCTCCAGCCTGGAATCCGAAGATTTCGCGGATTACTACTGTCTGCAACGCAACGCTTTCCCGCTGACTTTTGGTGCGGGTACTAAACTGGAGCTGAAACGCGCTGACGCTGCTCCGACTGTTTCCATCTTCCCGCCGTCTAGCGAACAACTGACGTCTGGTGGCGCCTCCGTTGTATGCTTCCTGAACAACTTCTACCCTAAGGACATCAACGTGAAATGGAAAATCGACGGTTCCGAACGTCAGAACGGTGTTCTGAACTCTTGGACCGATCAGGATAGCAAAGATTCCACCTACAGCATGTCCAGCACCCTGACCCTGACCAAAGACGAGTACGAACGTCACAACTCTTACACTTGTGAAGCTACTCACAAAACTAGCACGTCCCCTATTGTGAAATCCTTCAACCGCAATGAATGTTAAttaactcgaggctgagcaaagcagactactaataacataaagtctacgccggacgcatcgtggccctagtacgcaagttcacgtaaaaagggtaactagaggttgaggtgattttatgaaaaagaatatcgcatttcttcttgcatctatgttcgttttttctattgctacaaacgcgtacgctgagatctc**C**GAAGTGAAACTGCTGCAATCTGGTGGTGGTCTGGTTCAGCCGGGCGGTAGCCTGAAAGTCTCTTGTGCTGCGAGCGGTATCGACTTCTCTCGTAACTGGATGACCTGGGTCCGTCGCGCTCCGGGCAAAGGCCTGGAATGGATCGGTGAGATCTATCCGGATAGCTCCATTATCAACTATGCACCAAGCCTGAAAGACAAGTTCATCATCTCCCGTGACAACGCGAAGAACACCCTGTACCTGCAGATGTCCAAAGTTCGTTCTGAGGATACTGCGCTGTACTACTGTGCCCGTCGCACTACCGCATGGTACTTCGATGTTTGGGGTACCGGTACTTCCGTTACTGTTTCTTCTGCGAAAACTACTCCGCCGAGCGTATACCCGCTGGCACCGGGTTCTGCTGCACAAACCAACAGCATGGTTACCCTGGGTTGCCTGGTTAAAGGTTATTTCCCGGAGCCGGTTACCGTTACCTGGAACTCTGGCTCTCTGTCTTCTGGCGTTCATACTTTTCCGGCTGTGCTGCAGAGCGACCTGTATACCCTGAGCTCTTCTGTTACCGTACCGAGCTCTACCTGGCCTTCTCAAACCGTTACGTGTAATGTTGCTCACCCGGCTTCCTCTACGAAAGTGGACAAAAAGATCGTGCCACGCGATTGCGGTGCTAAACCGtctagacaccaccaccaccaccactaactcgagaattca | MKKNIAFLLASMFVFSIATNAYASDIQLIQSPSSIFASLGDRVSLSCRASQGIRGNLDWYQQKPGGTIKLLIYSTSNLKSGVPSRFSGRGSGSDYSLTISSLESEDFADYYCLQRNAFPLTFGAGTKLELKRADAAPTVSIFPPSSEQLTSGGASVVCFLNNFYPKDINVKWKIDGSERQNGVLNSWTDQDSKDSTYSMSSTLTLTKDEYERHNSYTCEATHKTSTSPIVKSFNRNEC  MKKNIAFLLASMFVFSIATNAYAEISEVKLLQSGGGLVQPGGSLKVSCAASGIDFSRNWMTWVRRAPGKGLEWIGEIYPDSSIINYAPSLKDKFIISRDNAKNTLYLQMSKVRSEDTALYYCARRTTAWYFDVWGTGTSVTVSSAKTTPPSVYPLAPGSAAQTNSMVTLGCLVKGYFPEPVTVTWNSGSLSSGVHTFPAVLQSDLYTLSSSVTVPSSTWPSQTVTCNVAHPASSTKVDKKIVPRDCGAKPSRHHHHHH |
| Fab 6D2 | gatatacatATGaaaaagaatatcgcatttcttcttgcatctatgttcgttttttctattgctacaaatgcctatgcatccGACATTCAGATGATCCAGAGCCCGTCCTCTATTTTCGCATCCCTGGGCGACCGTGTTTCCCTGTCCTGCCGTGCCTCTCAGGGTATCCGTGGCAATCTGGATTGGTACCAGCAGAAACCGGGTGGCACGATCAAGCTGCTGATCTATAGCACTTCCAACCTGAATTCCGGCGTGCCGTCTCGCTTCAGCGGTTCCGGTTCTGGTAGCGACTATTCTCTGACCATCAACAACCTGGAGTCCGATGACTTTGCCGATTATTACTGCCTGCAGCGCAACGCGTTTCCGCTGACTTTTGGTGCGGGCACCAAACTGGAACTGAAGCGTGCTGACGCTGCTCCGACCGTTTCTATCTTTCCGCCTAGCTCTGAGCAGCTGACCTCCGGTGGTGCGTCCGTGGTGTGTTTTCTGAACAACTTCTACCCGAAAGACATCAACGTTAAATGGAAAATCGACGGTTCCGAGCGTCAGAATGGTGTACTGAACTCCTGGACCGACCAGGACTCCAAAGATTCCACCTATTCTATGAGCTCCACCCTGACCCTGACCAAAGACGAATACGAACGTCACAACTCCTATACCTGCGAAGCCACTCACAAAACCTCCACGTCTCCGATCGTTAAAAGCTTTAACCGTAACGAATGCTAAttaactcgaggctgagcaaagcagactactaataacataaagtctacgccggacgcatcgtggccctagtacgcaagttcacgtaaaaagggtaactagaggttgaggtgattttatgaaaaagaatatcgcatttcttcttgcatctatgttcgttttttctattgctacaaacgcgtacgctgagatctc**C**GAAGTGAAACTGTTTCAATCCGGCGGCGGCCTGGTTCAGCCTGGTGGTTCTCTGAAATTCAGCTGCGCCGCGTCCGGTATCGATTTCTCTCGTTACTGGATGACCTGGGTTCGTCGTGCTCCGGGTAAGGGCCTGGAATGGATCGGTGAAATCTACCCGGACTCCTCTATTATCAACTACGCCCCGTCCCTGAAAGACAAATTCATTATCTCTCGCGATAACGCGAAATCTACCCTGTACCTGCAAATGACTAAGGTCCGCTCCGAGGATACCGCTCTGTACTACTGTGCGCGTCGTGAGAATGGCCACTGGTACTTCGACGTTTGGGGTACTGGTACTACGGTAACTGTGTCCAGCGCCAAGACCACTCCGCCGAGCGTTTACCCGCTGGCACCTGGTTCCGCTGCCCAGACTAATTCTATGGTGACCCTGGGCTGCCTGGTAAAAGGTTACTTCCCGGAGCCGGTAACTGTTACTTGGAACTCTGGTTCTCTGTCTTCTGGTGTGCATACCTTCCCGGCTGTACTGCAGTCCGACCTGTATACCCTGTCTAGCTCTGTCACTGTTCCAAGCAGCACTTGGCCGTCCCAAACCGTGACCTGTAACGTTGCACACCCAGCAAGCTCCACCAAAGTGGACAAAAAGATCGTTCCGCGTGACTGCGGTGCTAAACCGtctagacaccaccaccaccaccactaactcgagaattca | MKKNIAFLLASMFVFSIATNAYASDIQMIQSPSSIFASLGDRVSLSCRASQGIRGNLDWYQQKPGGTIKLLIYSTSNLNSGVPSRFSGSGSGSDYSLTINNLESDDFADYYCLQRNAFPLTFGAGTKLELKRADAAPTVSIFPPSSEQLTSGGASVVCFLNNFYPKDINVKWKIDGSERQNGVLNSWTDQDSKDSTYSMSSTLTLTKDEYERHNSYTCEATHKTSTSPIVKSFNRNEC  MKKNIAFLLASMFVFSIATNAYAEISEVKLFQSGGGLVQPGGSLKFSCAASGIDFSRYWMTWVRRAPGKGLEWIGEIYPDSSIINYAPSLKDKFIISRDNAKSTLYLQMTKVRSEDTALYYCARRENGHWYFDVWGTGTTVTVSSAKTTPPSVYPLAPGSAAQTNSMVTLGCLVKGYFPEPVTVTWNSGSLSSGVHTFPAVLQSDLYTLSSSVTVPSSTWPSQTVTCNVAHPASSTKVDKKIVPRDCGAKPSRHHHHHH |
| scFv  1E8b | gatatacatATGAAATATCTGCTGCCTACTGCAGCTGCTGGTCTGCTGCTGCTGGCAGCACAACCTGCTATGGCGGAAGTAAAACTTTTCCAATCAGGCGGCGGCTTAGTACAACCGGGCGGTTCTCTCAAGCTTTCATGCGCTGCTTCCGGGATTGACTTCAGCCGTAACTGGATGACCTGGGTTCGGAGAGCACCAGGTAAGGGCCTGGAGTGGATAGGTGAGATATACCCTGACTCACGGACTATCAATTACGCTCCAAGCCTCAAGGATAAATTCATTATTAGCCGCGATAACGCGAAGAAGATGCTGTATCTGCAAATGTCCAAAGTGCGGTCCGAGGACACGGCTCTTTATTACTGCGCCCGGCGAGGAGTGACAACAGTCGTTGGGACATCTTGGTACTTCGACGTATGGGGCACAGGAACGACTGTTACTGTGAGCAGCGCGGGCGGGAGTTCACGTAGTAGTAGCTCAGGCGGTGGTGGGTCAGGCGGCGGCGGCGATATTCAAATGATCCAGAGTCCGTCAAGCATGTTTGCCTCGTTAGGTGATCGCGTTAGCCTTTCTTGCCGTGCATCCCAGGGCATCCGCGGCAACCTGGATTGGTACCAACAAAAGCCCGGTGGCCCTATCAAGCTGCTTATACATTCAACGTCGAAATTAAACTCCGGAGTACCCTCAAGATTCAGCGGTAGTGGATCCGGCAGTGACTACTCATTGACTATTAGCTCTTTGGAGTCGGAAGACTTCGCCGATTACTACTGTCTCCAACGGAACGCATTTCCCCTTACTTTCGGGGCGGGCACGAAGTTAGAGCTGAAACGCGCAGGCAGACTCGAGCACCAC | MKYLLPTAAAGLLLLAAQPAMAEVKLFQSGGGLVQPGGSLKLSCAASGIDFSRNWMTWVRRAPGKGLEWIGEIYPDSRTINYAPSLKDKFIISRDNAKKMLYLQMSKVRSEDTALYYCARRGVTTVVGTSWYFDVWGTGTTVTVSSAGGSSRSSSSGGGGSGGGGDIQMIQSPSSMFASLGDRVSLSCRASQGIRGNLDWYQQKPGGPIKLLIHSTSKLNSGVPSRFSGSGSGSDYSLTISSLESEDFADYYCLQRNAFPLTFGAGTKLELKRAGRLEHHHHHH |

**Supplementary Table 9. Oligonucleotides for the creation of *H. pylori* mutants.** Primers used in the Gibson Assembly reactions to insert the chloramphenicol acetyl transferase (*cat*) cassette into *pseB* and *pseE* genes in *H. pylori*.

| **Primer** | **Sequence*** |
| --- | --- |
| PseB.1F | TGCTAGACAACCAAACGATT |
| PseB.1R-CAT | CGTAGTAGCTTGGTTTTAATGCATTCTAGGGGA |
| CAT-F-PseB.1 | TAAAACCAA**GCTACTACGGCAGGCTACTA** |
| CAT-R-PseB.2 | TAGGGATTT**TATCAGTGCGACAAACTGGGA** |
| PseB.2F-CAT | GCACTGATAAAATCCCTAGCATGAAAATAACTGA |
| PseB.2R | TAGTAATTTCAATAAATCATCAGGCTCT |
| PseE.1F | TTACCATTCTTTTAAAGCCATTTTGATCG |
| PseE.1R-CAT | CCGTAGTAGCAAAAAATTAGAAGAGTTGGATTTTGA |
| CAT-F-PseE.1 | CTAATTTTTT**GCTACTACGGCAGGCTACTA** |
| CAT-R-PseE.2 | ATGAAAATTT**TATCAGTCCGACAAACTGGGA** |
| PseE.2F-CAT | CGCACTGATAAAATTTTCATTACCTAAAAGCAAGCGA |
| PseE.2R | ATGGATATTTATCAAAAAAACTTACAAGCT |

***** Bold indicates cassette primer overlap with gene fragments; underline indicates gene fragment primer overlap with cassette.
