## Supplementary document 2 for "Uncovering bacterial pseudaminylation with pan-specific antibody tools"

**General methods and materials**

All reagents and solvents were used as received unless otherwise noted. Anhydrous MeOH, THF, MeCN, DMF and CH_2_Cl_2_ were obtained from a PureSolv^TM^ solvent dispensing unit. Unless otherwise stated, solution-phase reactions were carried out under an atmosphere of dry nitrogen or argon and at room temperature (22 °C). Reactions undertaken at -78 °C utilised a bath of dry ice and acetone. Reactions carried out at 0 °C employed a bath of water and ice. Reactions carried out at above room temperature were performed using a magnetic stirrer with heating in a multi-well heating mantle for direct heating of the reaction flask.

Flash column chromatography was performed using 230–400 mesh Kieselgel 60 silica eluting with gradients as specified. Analytical thin layer chromatography (TLC) was performed on commercially prepared silica plates (Merck Kieselgel 60 0.25 mm F254). Visualization of TLC plates was undertaken with ultraviolet (UV) light at λ = 254 nm and using staining with solutions of vanillin or phosphomolybdic acid, followed by exposure of the stained plates to heat.

^1^H NMR, ^13^C NMR and 2D NMR spectra were recorded at 300 K using a Bruker DRX500, DRX400 or DRX300 spectrometer. Chemical shifts are reported in parts per million (ppm) and are referenced to solvent residual signals: DMSO-*d*_6_ δ 2.50 [^1^H] and δ 39.52 [^13^C], CD_3_CN δ 1.94 [^1^H] and δ 1.32 [^13^C], CDCl_3_ δ 7.26 [^1^H] and δ 77.16 [^13^C], MeOD-*d*_4_ δ 3.31 [^1^H] and δ 49.00 [^13^C], D_2_O δ 4.79 [^1^H]. ^1^H NMR data is reported as chemical shift, multiplicity with associated coupling constant (*J* Hz) and assignment are made where possible.

High resolution ESI(+/-) mass spectra were measured on a Bruker–Daltonics Apex Ultra 7.0T Fourier transform mass spectrometer (FTICR) or on a Waters Micromass Q-Tof Premier mass spectrometer. Low resolution ESI(+/-) mass spectra were obtained on a Shimadzu 2020 ESI mass spectrometer operating in positive/negative ion mode. Infrared (IR) absorption spectra were recorded on a Bruker ALPHA Spectrometer with Attenuated Total Reflection (ATR) capability or on a Shimadzu IRAffinity-1 spectrometer. Optical rotations were recorded at ambient temperature (293K) on a Perkin–Elmer 341 polarimeter at 589 nm (sodium D line) with a cell path length of 1 dm. Melting points were determined with a SRS Optimelt melting point apparatus and are uncorrected.

Preparative RP‑HPLC was performed using a Waters 600 Multisolvent Delivery System and pump with Waters 486 Tuneable absorbance detector operating at 210-300 nm. Separations were performed using a mobile phase of 0.1% formic in water (Solvent A) and 0.1% formic acid in MeCN (Solvent B).

Analytical UPLC-MS was performed on a Shimadzu UPLC-MS system consisting of a LC‑M20A pump and a SPD-M30A diode array detector coupled to a Shimadzu 2020 mass spectrometer (ESI) operating in positive and/or negative mode. Separations on the UPLC-MS system were performed using a Waters Acquity UPLC BEH C_18_ 1.7 µm 130 Å (2.1 x 50 mm) column at a total flow rate of 0.60 mL/min. Separations were performed using a mobile phase of 0.1% formic acid in water (Solvent A) and 0.1% formic acid in MeCN (Solvent B).

Analytical RP‑HPLC was performed on a Waters Alliance e2695 HPLC system equipped with a 2998 PDA detector (λ = 210–400 nm). Separations were performed using a Waters SunFire C_18_ 5 µm 100 Å (2.1 × 150 mm) column at 40 ºC with a flow rate of 0.5 mL min^‑1^. Separations were performed using a mobile phase of 0.1% TFA in water (Solvent C) and 0.1% TFA in MeCN (Solvent D).

**General Procedure 1: α-selective glycosylation (0.1 mmol scale)**

A 10 mL Schlenk tube was dried with a heat gun under vacuum for 10 minutes before flushing with dry argon gas. To this tube was added AW-300 molecular sieves (200 mg, 100 mg/mL solvent) that were pre-activated in a microwave for 3 minutes, followed by addition of PseNHTroc adamantyl donor **3** (83.6 mg, 0.1 mmol, 1.0 equiv.) and an amino acid acceptor (2.0 equiv.). The tube was purged under vacuum and flushed with dry argon gas three times. Afterwards, anhydrous CH_2_Cl_2_ (2 mL, 2 mL/0.1 mmol of donor, final concentration: 50 mM) and anhydrous DMF (39 µL, 0.5 mmol, 5.0 equiv.) were added to dissolve the compounds. The reaction mixture was stirred at rt for 1 h and then cooled down to -78 °C before *N*‑iodosuccinimide (NIS) (54 mg, 0.24 mmol, 2.4 equiv.) was added. To initiate the reaction, triflic acid (TfOH) (0.9 µL, 0.01 mmol, 0.1 equiv.) was added dropwise at -78 °C. The reaction mixture was slowly warmed up to -40 °C and stirred for 12 h. The appearance of reddish-orange colour of the solution indicated successful activation of the glycosyl donor. Upon completion, as indicated by TLC, the reaction mixture was quenched with triethylamine (approx. 300 µL) until the reddish colour had turned into an orange-brown colour. The reaction mixture was warmed to rt and diluted with CH_2_Cl_2_ (3 mL). The iodine liberated from the activation of the donor was reduced by the addition of sat. aq. Na_2_S_2_O_3_ solution (3 mL) and stirred vigorously until the solution had turned colourless. Afterwards, this mixture was filtered through a pad of celite to remove molecular sieves. The organic layer was separated from the filtrate, dried with anhydrous MgSO_4_, concentrated *in vacuo* and purified by flash column chromatography.

**General Procedure 2: β-selective glycosylation (0.1 mmol scale)**

A 10 mL Schlenk tube was dried with a heat gun under vacuum for 10 minutes before flushing it with dry argon gas. To this tube was added AW-300 molecular sieves (200 mg, 100 mg/mL solvent) that were pre-activated in a microwave for 3 minutes, followed by addition of PseN_3_ adamantyl donor **4** (68.7 mg, 0.1 mmol, 1.0 equiv.) and an amino acid acceptor (2.0 equiv.). The tube was purged under vacuum and flushed with dry argon gas three times. Afterwards, anhydrous CH_2_Cl_2_ and MeCN (2 mL, CH_2_Cl_2_:MeCN 2:1 v/v, 2 mL/0.1 mmol of donor, final concentration: 50 mM) was added to dissolve the compounds. The reaction mixture was stirred at rt for 1 h and then cooled down to -78 °C before NIS (54 mg, 0.24 mmol, 2.4 equiv.) was added. To initiate the reaction, TfOH (0.9 µL, 0.01 mmol, 0.1 equiv.) was added dropwise at -78 °C. The reaction mixture was maintained at -78 °C and stirred for 12 h. The appearance of reddish‑orange colour of the solution indicated a successful activation of the glycosyl donor. Upon completion, as indicated by TLC, the reaction mixture was quenched by triethylamine (approx. 300 µL) until the reddish colour had turned into an orange-brown colour. The reaction mixture was warmed to rt and diluted with CH_2_Cl_2_ (3 mL). The iodine liberated from the activation of the donor was reduced by the addition of sat. aq. Na_2_S_2_O_3_ solution (3 mL) and stirred vigorously until the solution had turned colourless. Afterwards, this mixture was filtered through a pad of celite to remove the molecular sieves. The organic layer was separated from the filtrate, dried with anhydrous MgSO_4_, concentrated *in vacuo* and purified by flash column chromatography.

**General Procedure 3: Resin Loading**

2-Chlorotrityl chloride (2-CTC) resin (manufacturer’s resin loading: 1.6 mmol/g, 500-600 mg) was swelled in dry CH_2_Cl_2_ (5 mL) for 30 min. The resin was subsequently shaken with a solution of 10% (v/v) *i*Pr_2_NEt in CH_2_Cl_2_ (5 mL) for 30 min and then washed with CH_2_Cl_2_ (5x5 mL). A solution of Fmoc-protected amino acid (1.0 equiv. based on resin loading) and *i*Pr_2_NEt (2.0 equiv. based on resin loading) in CH_2_Cl_2_ (5 mL) was prepared and the resin was shaken with the solution for 16 h. The resin was subsequently washed with CH_2_Cl_2_ (5x5 mL), DMF (5x5 mL) and CH_2_Cl_2_ (5x5 mL) before treating with a capping cocktail of CH_2_Cl_2_/MeOH/ *i*Pr_2_NEt (5 mL, 17:2:1 v/v/v) for 1 h. The resin was subsequently washed with DMF (5x5 mL), CH_2_Cl_2_ (5x5 mL) and DMF (5x5 mL). The efficiency of amino acid loading was determined by UV spectrophotometric analysis of the filtrate after Fmoc deprotection (20% (v/v) piperidine in DMF, 2x5 min) at λ = 301 nm (ε = 7800 M^-1^ cm^-1^).

**General Procedure 4: Fmoc-strategy Iterative Peptide Coupling with DIC/Oxyma**

Fmoc-deprotection was achieved by treating the resin with 20% (v/v) piperidine in DMF (2x5 mL) for 5 min. Afterwards, the resin was washed with DMF (5x5 mL), CH_2_Cl_2_ (5x5 mL) and DMF (5x5 mL). The resin was then treated with a solution of Fmoc-protected amino acid (4.0 equiv. relative to resin loading), *N,N'*-diisopropylcarbodiimide (DIC) (4.0 equiv. relative to resin loading) and ethyl cyano(hydroxyimino)acetate (Oxyma Pure) (8.0 equiv. relative to resin loading) in DMF (final concentration >0.1 M) for 1 h. The resin was subsequently washed with DMF (5x5 mL), CH_2_Cl_2_ (5x5 mL) and DMF (5x5 mL) before capping any uncoupled sequences by treating the resin with a solution of 10% (v/v) acetic anhydride in pyridine (2x5 mL). The resin was again washed with DMF (5x5 mL), CH_2_Cl_2_ (5x5 mL) and DMF (5x5 mL). The above process was repeated iteratively until the desired peptide sequence was assembled on resin.

**General Procedure 5: Glycosylamino acid coupling and Alloc deprotection**

A solution of pseudaminylated building block (**α‑1**/**β‑1**) (1.0 equiv. relative to resin loading), 1‑[*bis*(dimethylamino)methylene]-1*H*-1,2,3-triazolo[4,5-b]pyridinium 3-oxide hexafluorophosphate (HATU) (1.0 equiv. relative to resin loading), 1-hydroxy-7-azabenzotriazole (HOAt) (1.0 equiv. relative to resin loading) and 2,4,6-trimethylpyridine (TMP) (1.0 equiv. relative to resin loading) in DMF (final concentration >0.1 M) was pre‑activated for 10 min before the resin was treated with the solution for 1 h. The resin was subsequently washed with DMF (5x5 mL), CH_2_Cl_2_ (5x5 mL) and DMF (5x5 mL) before capping any uncoupled sequences by treating the resin with a solution of 10% (v/v) acetic anhydride in pyridine (2x5 mL). The resin was again washed with DMF (5x5 mL), CH_2_Cl_2_ (5x5 mL) and DMF (5x5 mL). Next, Alloc deprotection was performed by treating the resin with a solution of Pd(PPh_3_)_4_ (0.2 equiv. relative to resin loading) and PhSiH_3_ (20 equiv. relative to resin loading) in CH_2_Cl_2_ (5 mL) for 30 min. Upon completion, the resin was washed with DMF (5x5 mL), CH_2_Cl_2_ (5x5 mL) and DMF (5x5 mL), and then immediately coupled with the next amino acid residue to prevent intramolecular cyclisation.

**General Procedure 6: Simultaneous resin cleavage and global side chain deprotection**

To the resin‑bound peptide was added a solution of TFA/*i*Pr_3_SiH/H_2_O (90:5:5 v/v/v) and the resin shaken at rt for 3 h. The cleavage solution was collected and concentrated under a stream of nitrogen until the volume of the solution was less than 0.5 mL. Subsequently, cold Et_2_O (35 mL) was added to this solution to precipitate the peptide. The resulted suspension was centrifuged at 0 °C and the supernatant was discarded.

**General Procedure 7: Glycopeptide saponification**

The crude peptide was treated with an aqueous solution of LiOH (50 mM) until it was completely dissolved. The reaction mixture was monitored by UPLC‑MS at regular intervals. Extra LiOH (1.0 equiv.) was added to reactions that stalled (as judged by UPLC-MS analysis). Upon completion, the solution was adjusted to pH 4-5 with dropwise addition of formic acid. Finally, the crude mixture was purified by preparative RP-HPLC and then lyophilised.

***tert*-Butyl *N*-(*tert*-butoxycarbonyl)-*O*-(4,8-di-*O*-acetyl-7-*N*-benzyloxycarbonyl-1-isopropyl-5-*N*-(2,2,2-trichloroethoxycarbonyl)-α-pseudaminosyl)-L-serinate (α‑7)**

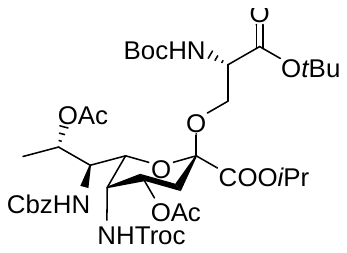

PseNHTroc adamantyl thioglycoside donor **3** (585 mg, 0.70 mmol) was glycosylated with commercially available Boc-Ser(OH)-O*t*Bu acceptor **6** (366 mg, 1.40 mmol, 2.0 equiv.) according to general procedure 1. The crude product was purified by flash column chromatography (eluent gradient: 0%-25% EtOAc in hexane) to afford the *title* compound **α‑7** as a yellow foam (589 mg, 91%).

**[α]^20^_D_** = -21 (c = 0.7, CH_2_Cl_2_); **IR** (thin film) v*_max_* = 2956, 2924, 2870, 2854, 1726, 1538, 1504, 1458, 1392, 1311, 1234, 1154, 1105, 1056, 1043, 969, 847, 818, 776, 772 cm^-1^; **^1^H NMR** (400 MHz, CDCl_3_) δ 7.39 – 7.33 (m, 5H, Ar-H), 5.49 (d, *J* = 9.8 Hz, 1H, NH), 5.35 (d, *J* = 9.0 Hz, 1H, NH), 5.18 (d, *J* = 5.9 Hz, 1H, H-8), 5.14 – 5.05 (m, 3H, H-4, *i*Pr-CH, PhC*H_2_*''), 5.01 (d, *J* = 8.0 Hz, 1H, CCl_3_CH_2_''), 4.98 (d, *J* = 8.1 Hz, 1H, PhC*H_2_*'), 4.89 (d, *J* = 10.6 Hz, 1H, NH), 4.45 (d, *J* = 12.1 Hz, 1H, CCl_3_CH_2_'), 4.34 (s, 2H, H5, Ser α-H), 4.24 (t, *J* = 10.1 Hz, 1H, H-7), 4.05 (d, *J* = 8.4 Hz, 1H, SerCH_2_''), 3.98 – 3.87 (m, 1H, H-6), 3.48 (d, *J* = 9.3 Hz, 1H, SerCH_2_'), 2.23 (dd, *J* = 13.4, 4.9 Hz, 1H, H-3eq), 2.06 (s, 3H, CH_3_CO), 1.96 (s, 3H, CH_3_CO), 1.85 (t, *J* = 12.8 Hz, 1H, H-3ax), 1.50 (s, 9H, 3 x CH_3_), 1.46 – 1.45 (m, 14H, 3 x CH_3_), 1.37 (d, *J* = 6.6 Hz, 3H, H-9), 1.32 (d, *J* = 4.3 Hz, 3H, *i*Pr-CH_3_''), 1.31 (d, *J* = 4.3 Hz, 3H, *i*PrCH_3_'); **^13^C NMR** (101 MHz, CDCl_3_) δ 170.5, 170.1, 169.2, 166.4, 155.8, 155.5, 155.0, 136.3, 128.7, 128.4, 128.3, 99.0, 95.9, 83.0, 80.1, 74.7, 71.7, 71.0, 70.7, 67.4, 67.1, 65.5, 54.1, 53.3, 48.3, 32.1, 28.5, 28.3, 21.8, 21.8, 20.9, 16.3; α only (*J*_C1-H3ax_ = 0 Hz); **HRMS** (ESI+) *m/z* Calc. for C_39_H_56_Cl_3_N_3_O_16_ [M + Na]^+^: 950.2618, found 950.2632.

***tert*-Butyl *N*-(*tert*-butoxycarbonyl)-*O*-(5-acetamido-4,8-di-*O*-acetyl-7-*N*-benzyloxycarbonyl-1-isopropyl-α-pseudaminosyl)-L-serinate (α‑S1)**

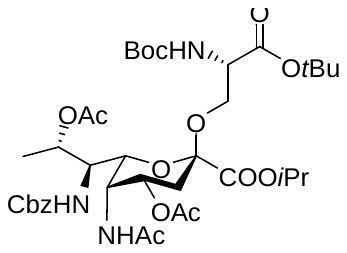

Compound **α‑7** (186 mg, 201 µmol) and zinc powder (131 mg, 2.01 mmol, 10 equiv.) were dissolved in acetic acid (4 mL) and acetic anhydride (4 mL). The reaction mixture was stirred at 50 °C for 2 h. Upon completion, the reaction mixture was diluted with EtOAc (60 mL) and filtered through a pad of celite. The collected filtrate was washed with sat. aq. NaHCO_3_ solution (4x50 mL) and brine (1x50 mL). The organic layer was collected, dried with anhydrous MgSO_4_ and concentrated *in vacuo*. The crude product was purified by flash column chromatography (eluent gradient: 0%-75% EtOAc in hexane) to afford the *title* compound **α‑S1** as a yellow foam (146 mg, 92%).

**[α]^20^_D_** = -47 (c = 0.9, CH_2_Cl_2_); **IR** (thin film) v*_max_* = 2978, 2927, 1722, 1535, 1509, 1456, 1368, 1309, 1234, 1153, 1086, 1051, 907, 859, 845, 800, 775, 738, 698, 485 cm^-1^; **^1^H NMR** (400 MHz, CDCl_3_) δ 7.40 – 7.27 (m, 5H, Ar-H), 6.26 (d, *J* = 9.4 Hz, 1H, NH), 5.35 (d, *J* = 9.0 Hz, 1H, NH), 5.23 – 5.05 (m, 4H, H-4, H-8, PhC*H_2_*'', *i*Pr-CH), 4.95 (d, *J* = 12.1 Hz, 1H, PhC*H_2_*'), 4.82 (d, *J* = 10.8 Hz, 1H, NH), 4.60 (d, *J* = 9.4 Hz, 1H, H-5), 4.41 – 4.24 (m, 2H, H-7, Ser α-H), 3.98 (d, *J* = 9.2 Hz, 1H, SerCH_2_''), 3.89 (d, *J* = 10.2 Hz, 1H, H-6), 3.45 (d, *J* = 8.9 Hz, 1H, SerCH_2_'), 2.19 (dd, *J* = 13.2, 5.0 Hz, 1H, H-3eq), 2.03 (s, 6H, 2 x CH_3_CO), 1.97 (s, 3H, CH_3_CO), 1.80 (t, *J* = 12.8 Hz, 1H, H-3ax), 1.50 (s, 9H, 3 x CH_3_), 1.45 (s, 9H, 3 x CH_3_), 1.33 (d, *J* = 4.3 Hz, 3H, H-9), 1.33 – 1.29 (m, 6H, 2 x *i*Pr-CH_3_); **^13^C NMR** (101 MHz, CDCl_3_) δ 171.0, 170.6, 170.2, 169.1, 166.8, 156.0, 155.5, 136.3, 128.6, 128.4, 128.3, 99.1, 83.0, 80.1, 71.2, 71.0, 70.8, 67.5, 67.0, 65.6, 54.1, 53.0, 45.8, 32.3, 28.5, 28.3, 23.4, 21.7, 21.3, 21.0, 15.3; **HRMS** (ESI+) *m/z* Calc. for C_38_H_57_N_3_O_15_ [M + Na]^+^: 818.3682, found 818.3679.

***tert*-Butyl *N*-(*tert*-butoxycarbonyl)-*O*-(5,7-diacetamido-4,8-di-*O*-acetyl-1-isopropyl-α-pseudaminosyl)-L-serinate (α‑S2)**

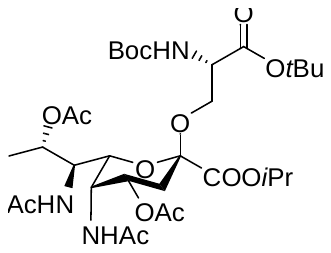

Compound **α‑S1** (142 mg, 179 µmol) was dissolved in MeOH (5 mL) before Pd/C (95.2 mg, 89.5 µmol, 0.5 equiv., 10 wt% palladium on activated carbon) was added. The reaction mixture was stirred under 1 atm H_2_ at rt for 2 h. Upon completion, the reaction mixture was filtered through a pad of celite, and the collected filtrate was concentrated *in vacuo*. The Cbz‑deprotected intermediate was dissolved in CH_2_Cl_2_ (10 mL) before triethylamine (125 µL, 895 µmol, 5.0 equiv.) and acetic anhydride (84.6 µL, 895 µmol, 5.0 equiv.) were added. The reaction mixture was stirred at rt for 1 h. Upon completion, the reaction mixture was diluted with CH_2_Cl_2_ (40 mL), washed with 1 M HCl (2x50 mL), sat. aq. NaHCO_3_ solution (1x50 mL) and brine (1x50 mL). The organic layer was collected, dried with anhydrous MgSO_4_ and concentrated *in vacuo*. The crude product was purified by flash column chromatography (eluent: 45% EtOAc and 5% MeOH in hexane) to afford the *title* compound **α‑S2** as a yellow oil (98.1 mg, 78% over 2 steps).

**[α]^25^_D_** = -59 (c = 1.3, CH_2_Cl_2_); **IR** (thin film) v*_max_* = 2978, 2924, 2853, 1723, 1664, 1542, 1508, 1457, 1368, 1257, 1235, 1152, 1088, 1031, 971, 907, 845, 800, 736, 702, 602, 492 cm^-1^; **^1^H NMR** (400 MHz, CDCl_3_) δ 6.21 (d, *J* = 9.4 Hz, 1H, NH), 5.66 (d, *J* = 10.5 Hz, 1H, NH), 5.36 (d, *J* = 9.0 Hz, 1H, NH), 5.13 – 5.06 (m, 3H, H-4, H-8, *i*Pr-CH), 4.60 – 4.45 (m, 2H, H-5, H-7), 4.33 (d, *J* = 9.0 Hz, 1H, Ser α-H), 4.06 – 4.00 (m, 1H, SerCH_2_''), 3.98 (d, *J* = 10.2 Hz, 1H, H-6), 3.46 (d, *J* = 9.6 Hz, 1H, SerCH_2_'), 2.17 (dd, *J* = 12.8, 4.8 Hz, 1H, H-3eq), 2.07 (s, 3H, CH_3_CO), 1.98 (s, 3H, CH_3_CO), 1.96 (s, 3H, CH_3_CO), 1.93 (s, 3H, CH_3_CO), 1.80 (t, *J* = 12.7 Hz, 1H, H-3ax), 1.51 (s, 9H, 3 x CH_3_), 1.45 (s, 9H, 3 x CH_3_), 1.35 (d, *J* = 6.7 Hz, 3H, H-9), 1.32 (d, *J* = 5.9 Hz, 3H, *i*Pr-CH_3_''), 1.30 (d, *J* = 6.0 Hz, 3H, *i*Pr-CH_3_'); **^13^C NMR** (101 MHz, CDCl_3_) δ 171.2, 170.8, 170.4, 170.0, 169.2, 166.8, 155.5, 99.1, 82.9, 80.1, 71.7, 70.9, 70.8, 67.0, 65.6, 54.1, 50.7, 45.7, 32.2, 28.5, 28.2, 23.4, 23.3, 21.8, 21.7, 21.4, 21.0, 16.0; **HRMS** (ESI+) *m/z* Calc. for C_32_H_53_N_3_O_14_ [M + Na]^+^: 726.3420, found 726.3424.

***N*-(allyloxycarbonyl)-*O*-(5,7-diacetamido-4,8-di-*O*-acetyl-1-isopropyl-α-pseudaminosyl)-L-serine (α‑1)**

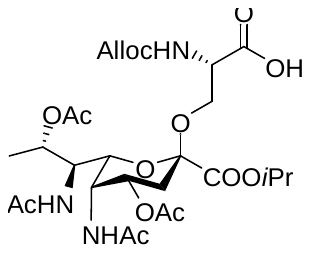

Compound **α‑S2** (308 mg, 438 µmol) was dissolved in a solution of TFA/water (10 mL, 19:1 v/v). The reaction mixture was stirred at rt for 2 h. Upon completion, the reaction mixture was concentrated *in vacuo*. This dried intermediate was redissolved in THF (15 mL) and sat. aq. NaHCO_3_ (15 mL). Allyl chloroformate (46.5 µL, 438 µmol, 1.0 equiv.) was added and the reaction mixture was stirred vigorously at rt for 12 h. Upon completion, the reaction mixture was extracted with EtOAc (60 mL). The organic layer was collected, dried with anhydrous MgSO_4_ and concentrated *in vacuo*. The crude product was purified by flash column chromatography (eluent gradient: 0%-10% MeOH in CH_2_Cl_2_ with 1% acetic acid as additive) to afford the *title* compound **α‑1** as a yellow oil (250 mg, 90% over 2 steps).

**[α]^25^_D_** = -43 (c = 1.1, MeOH); **IR** (thin film) v*_max_* = 3305, 3089, 2985, 2941, 1723, 1664, 1550, 1431, 1376, 1246, 1205, 1183, 1142, 1104, 1051, 721 cm^-1^; **^1^H NMR** (500 MHz, DMSO-*d*_6_) δ 7.78 (d, *J* = 10.6 Hz, 1H, NH), 7.75 (d, *J* = 10.6 Hz, 1H, NH), 7.36 (d, *J* = 8.3 Hz, 1H, NH), 5.89 (ddt, *J* = 16.1, 10.4, 5.3 Hz, 1H, CH_2_C*H*=CH_2_), 5.35 – 5.25 (m, 1H, CH_2_CH=C*H_2_*''), 5.21 – 5.16 (m, 1H, CH_2_CH=C*H_2_*'), 5.07 – 4.99 (m, 2H, H-4, H-8), 4.96 (p, *J* = 6.2 Hz, 1H, *i*Pr-CH), 4.48 (d, *J* = 5.3 Hz, 2H, C*H_2_*CH=CH_2_), 4.45 – 4.38 (m, 1H, H-5), 4.26 – 4.18 (m, 2H, H-7, Ser α-H), 3.90 (dd, *J* = 10.3, 2.1 Hz, 1H, H-6), 3.74 (dd, *J* = 10.0, 5.5 Hz, 1H, SerCH_2_''), 3.60 (dd, *J* = 10.0, 5.2 Hz, 1H, SerCH_2_'), 2.05 – 1.97 (m, 2H, H-3), 1.90 (s, 3H, CH_3_CO), 1.84 (s, 3H, CH_3_CO), 1.77 (s, 3H, CH_3_CO), 1.69 (s, 3H, CH_3_CO), 1.25 (d, *J* = 3.9 Hz, 3H, *i*Pr-CH_3_''), 1.23 (d, *J* = 3.8 Hz, 3H, *i*PrCH_3_'), 1.20 (d, *J* = 6.7 Hz, 2H, H-9); **^13^C NMR** (126 MHz, DMSO-*d*_6_) δ 171.3, 169.9, 169.5, 169.4, 168.7, 165.5, 155.7, 133.3, 117.1, 98.1, 70.3, 70.2, 69.2, 66.7, 64.6, 63.4, 53.7, 49.8, 44.5, 31.0, 22.6, 22.5, 21.3, 21.2, 20.9, 20.7, 15.2; **HRMS** (ESI+) *m/z* Calc. for C_27_H_41_N_3_O_14_ [M + Na]^+^: 654.2481, found 654.2477.

***tert*-Butyl *N*-(tert-butoxycarbonyl)-*O*-(4,8-di-*O*-acetyl-5-azido-7-*N*-benzyloxycarbonyl-1-isopropyl-β-pseudaminosyl)-L-serinate (β‑7)**

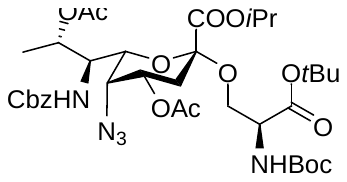

PseN_3_ adamantyl thioglycoside donor **4** (275 mg, 0.40 mmol) was glycosylated with commercially available Boc-Ser(OH)-O*t*Bu acceptor **6** (209 mg, 0.80 mmol, 2.0 equiv.) according to general procedure 2. The crude product (β:α = 10:1) was purified by flash column chromatography (eluent gradient: 0%-25% EtOAc in hexane) to afford the *title* compound **β‑7** as a yellow foam (214 mg, 69%).

**[α]^25^_D_** = -35 (c = 1.2, CH_2_Cl_2_); **IR** (thin film) v*_max_* = 2979, 2935, 2112, 1735, 1531, 1509, 1456, 1391, 1368, 1304, 1229, 1187, 1153, 1094, 1057, 1001, 848, 745, 699 cm^-1^; **^1^H NMR** (400 MHz, CDCl_3_) δ 7.42 – 7.28 (m, 5H, Ar-H), 5.29 (d, *J* = 8.8 Hz, 1H, NH), 5.20 (dd, *J* = 6.8, 3.6 Hz, 1H, H-8), 5.16 – 5.03 (m, 3H, PhC*H_2_*, *i*Pr-CH), 4.98 (d, *J* = 9.3 Hz, 1H, NH), 4.89 (ddd, *J* = 12.7, 4.5, 3.1 Hz, 1H, H-4), 4.40 (td, *J* = 9.6, 3.4 Hz, 1H, H-7), 4.27 (d, *J* = 8.3 Hz, 1H, Ser α-H), 3.91 (dd, *J* = 9.3, 3.3 Hz, 1H, SerCH_2_''), 3.85 (s, 1H, H-5), 3.81 – 3.62 (m, 2H, H-6, SerCH_2_'), 2.36 (dd, *J* = 12.6, 4.5 Hz, 1H, H-3eq), 2.17 (t, *J* = 12.4 Hz, 1H, H-3ax), 2.11 (s, 3H, CH_3_CO), 1.98 (s, 3H, CH_3_CO), 1.45 (s, 18H, 6 x CH_3_), 1.34 – 1.23 (m, 9H, H-9, 2 x *i*Pr-CH_3_); **^13^C NMR** (101 MHz, CDCl_3_) δ 170.2, 170.0, 169.2, 167.0, 156.2, 155.6, 136.4, 128.7, 128.4, 128.3, 99.1, 82.2, 79.9, 72.7, 70.7, 70.0, 69.8, 67.4, 65.1, 58.8, 54.1, 53.9, 32.0, 28.5, 28.1, 21.8, 21.2, 20.8, 14.7; β only (*J*_C1-H3ax_ = 6.4 Hz); **HRMS** (ESI+) *m/z* Calc. for C_36_H_53_N_5_O_14_ [M + Na]^+^: 802.3481, found 802.3480.

***tert*-Butyl *N*-(tert-butoxycarbonyl)-*O*-(5,7-diacetamido-4,8-di-*O*-acetyl-1-isopropyl-β-pseudaminosyl)-L-serinate (β‑S2)**

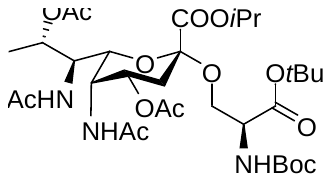

Compound **β‑7** (214 mg, 275 µmol) was dissolved in MeOH (11 mL) before Pd/C (147 mg, 138 µmol, 0.5 equiv., 10 wt% palladium on activated carbon) was added. The reaction mixture was stirred under 1 atm H_2_ at rt for 2 h. Upon completion, the reaction mixture was filtered through a pad of celite, and the collected filtrate was concentrated *in vacuo*. The free amine intermediate was dissolved in CH_2_Cl_2_ (11 mL) before triethylamine (192 µL, 1.37 mmol, 5.0 equiv.) and acetic anhydride (130 µL, 1.37 mmol, 5.0 equiv.) were added. The reaction mixture was stirred at rt for 1 h. Upon completion, the reaction mixture was diluted with CH_2_Cl_2_ (60 mL), washed with 1 M HCl (2x50 mL), sat. aq. NaHCO_3_ solution (1x50 mL) and brine (1x50 mL). The organic layer was collected, dried with anhydrous MgSO_4_ and concentrated *in vacuo*. The crude product was purified by flash column chromatography (eluent: 45% EtOAc and 5% MeOH in hexane) to afford the *title* compound **β‑S2** as a yellow oil (127 mg, 66% over 2 steps).

**[α]^25^_D_** = -52 (c = 0.9, CH_2_Cl_2_); **IR** (thin film) v*_max_* = 2980, 2930, 1736, 1664, 1542, 1455, 1368, 1281, 1239, 1204, 1156, 1102, 1069, 1047, 847 cm^-1^; **^1^H NMR** (400 MHz, CDCl_3_) δ 6.95 (d, *J* = 9.6 Hz, 1H, NH), 5.44 (d, *J* = 10.4 Hz, 1H, NH), 5.24 (d, *J* = 9.2 Hz, 1H, NH), 5.18 (td, *J* = 6.7, 2.7 Hz, 1H, H-8), 5.11 (p, *J* = 6.3 Hz, 1H, *i*Pr-CH), 4.83 (dd, *J* = 10.6, 6.6 Hz, 1H, H-4), 4.68 (dt, *J* = 8.7, 5.2 Hz, 1H, Ser α-H), 4.60 – 4.49 (m, 2H, H-5, H-7), 4.18 (dd, *J* = 10.8, 3.6 Hz, 1H, SerCH_2_''), 3.97 (dd, *J* = 9.7, 1.8 Hz, 1H, H-6), 3.52 – 3.43 (m, 1H, SerCH_2_'), 2.39 (dd, *J* = 13.1, 4.7 Hz, 1H, H-3eq), 2.04 (s, 3H, CH_3_CO), 2.03 (s, 3H, CH_3_CO), 1.98 (s, 3H, CH_3_CO), 1.92 (s, 3H, CH_3_CO), 1.85 (t, *J* = 13.2 Hz, 1H, H-3ax), 1.47 (s, 9H, 3 x CH_3_), 1.44 (s, 9H, 3 x CH_3_), 1.38 (d, *J* = 6.7 Hz, 3H, H-9), 1.32 (s, 3H, *i*Pr-CH_3_''), 1.31 (s, 3H, *i*Pr-CH_3_'); **^13^C NMR** (101 MHz, CDCl_3_) δ 171.5, 170.7, 170.4, 169.8, 169.1, 167.5, 155.7, 99.5, 82.7, 79.9, 72.9, 71.2, 70.6, 67.8, 65.9, 55.3, 53.6, 50.7, 45.5, 32.9, 28.5, 28.1, 23.5, 23.2, 21.9, 21.9, 21.4, 21.1, 15.1; **HRMS** (ESI+) *m/z* Calc. for C_32_H_53_N_3_O_14_ [M + Na]^+^: 726.3420, found 726.3407.

***N*-(allyloxycarbonyl)-*O*-(5,7-diacetamido-4,8-di-*O*-acetyl-1-isopropyl-β-pseudaminosyl)-L-serine (β‑1)**

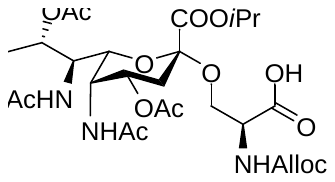

Compound **β‑S2** (127 mg, 181 µmol) was dissolved in a solution of TFA/water (5 mL, 19:1 v/v). The reaction was stirred at rt for 2 h. Upon completion, the reaction mixture was concentrated *in vacuo*. The resulting residue was redissolved in THF (6 mL) and sat. aq. NaHCO_3_ (6 mL). Allyl chloroformate (19.2 µL, 181 µmol, 1.0 equiv.) was added and the reaction mixture was stirred vigorously at rt for 12 h. Upon completion, the reaction mixture was extracted with EtOAc (60 mL). The organic layer was collected, dried with anhydrous MgSO_4_ and concentrated *in vacuo*. The crude product was purified by flash column chromatography (eluent gradient: 0%-10% MeOH in CH_2_Cl_2_ with 1% acetic acid as additive) to afford the *title* compound **β‑7** as a yellow oil (90.0 mg, 71% over 2 steps).

**[α]^25^_D_** = -39 (c = 0.8, MeOH); **IR** (thin film) v*_max_* = 3347, 2986, 2927, 1723, 1659, 1550, 1423, 1373, 1330, 1251, 1204, 1186, 1138, 1102, 1067, 1045, 931 cm^-1^; **^1^H NMR** (400 MHz, DMSO-*d*_6_) δ 7.76 (d, *J* = 10.0 Hz, 1H, NH), 7.62 (d, *J* = 9.8 Hz, 1H, NH), 7.24 (s, 1H, NH), 5.98 – 5.83 (m, 1H, CH_2_C*H*=CH_2_), 5.31 (dd, *J* = 17.3, 1.7 Hz, 1H, CH_2_CH=C*H_2_*''), 5.18 (dd, *J* = 10.4, 1.7 Hz, 1H, CH_2_CH=C*H_2_*'), 5.09 (dd, *J* = 6.7, 3.1 Hz, 1H, H-8), 5.03 (p, *J* = 6.2 Hz, 1H, *i*Pr-CH), 4.67 (dt, *J* = 13.3, 4.2 Hz, 1H, H-4), 4.49 (dt, *J* = 5.4, 1.6 Hz, 2H, C*H_2_*CH=CH_2_), 4.39 – 4.33 (m, 1H, H-5), 4.31 (dd, *J* = 10.1, 3.1 Hz, 1H, H-7), 4.29 – 4.18 (m, 1H, Ser α-H), 4.05 (dd, *J* = 10.2, 3.7 Hz, 1H, SerCH_2_''), 3.73 (dd, *J* = 10.3, 2.1 Hz, 1H, H-6), 3.64 (dd, *J* = 10.1, 7.2 Hz, 1H, SerCH_2_'), 2.20 (dd, *J* = 12.4, 4.4 Hz, 1H, H-3eq), 1.94 (t, *J* = 12.6 Hz, 1H, H-3ax), 1.87 (s, 3H, CH_3_CO), 1.86 (s, 3H, CH_3_CO), 1.79 (s, 3H, CH_3_CO), 1.69 (s, 3H, CH_3_CO), 1.27 (d, *J* = 6.5 Hz, 3H, *i*Pr-CH_3_''), 1.24 (d, *J* = 6.6 Hz, 3H, *i*Pr-CH_3_'), 1.20 (d, *J* = 6.6 Hz, 3H, H-9); **^13^C NMR** (101 MHz, DMSO-*d*_6_) δ 171.1, 169.9, 169.4, 169.3, 168.7, 166.8, 155.8, 133.4, 117.1, 99.1, 72.4, 69.9, 69.2, 67.2, 64.5, 63.9, 54.4, 49.3, 44.2, 32.2, 22.6, 22.4, 21.3, 21.3, 21.0, 20.6, 13.6; **HRMS** (ESI+) *m/z* Calc. for C_27_H_41_N_3_O_14_ [M + Na]^+^: 654.2481, found 654.2475.

***tert*-Butyl *N*-(tert-butoxycarbonyl)-*O*-(4,8-di-*O*-acetyl-7-*N*-benzyloxycarbonyl-1-isopropyl-5-*N*-(2,2,2-trichloroethoxycarbonyl)-α-pseudaminosyl) L-threoninate (α‑S3)**

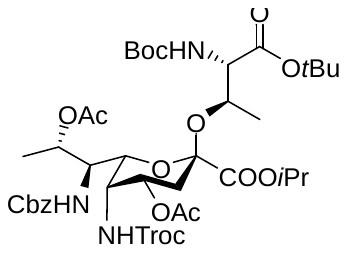

PseNHTroc adamantyl thioglycoside donor **3** (418 mg, 0.50 mmol) was glycosylated with commercially available Boc-Thr(OH)-O*t*Bu acceptor (366 mg, 1.00 mmol, 2.0 equiv.) according to general procedure 1. The crude product was purified by flash column chromatography (eluent gradient: 0%-25% EtOAc in hexane) to afford the *title* compound **α‑S3** as a yellow foam (360 mg, 76%).

**[α]^25^_D_** = -34 (c = 1.4, CH_2_Cl_2_); **IR** (thin film) v*_max_* = 2980, 2929, 1720, 1505, 1455, 1368, 1312, 1258, 1230, 1151, 1089, 1027, 907, 873, 800, 736, 699, 602, 571, 486 cm^-1^; **^1^H NMR** (400 MHz, CDCl_3_) δ 7.39 – 7.26 (m, 5H, Ar-H), 5.67 (d, *J* = 9.6 Hz, 1H, NH), 5.57 (d, *J* = 10.5 Hz, 1H, NH), 5.40 – 5.16 (m, 2H, H-8, NH), 5.15 – 4.89 (m, 5H, H-4, CCl_3_CH_2_'', PhC*H_2_*, *i*Pr-CH), 4.52 (d, *J* = 12.2 Hz, 1H, CCl_3_CH_2_'), 4.40 – 4.31 (m, 2H, H-5, Thr β-H), 4.27 (ddd, *J* = 10.9, 8.3, 3.6 Hz, 1H, H-7), 4.19 (dd, *J* = 8.9, 2.8 Hz, 1H, Thr α-H), 4.10 (d, *J* = 8.2 Hz, 1H, H-6), 2.21 (dd, *J* = 13.4, 5.0 Hz, 1H, H-3eq), 2.04 (s, 3H, CH_3_CO), 1.95 (s, 3H, CH_3_CO), 1.73 (t, *J* = 12.8 Hz, 1H, H-3ax), 1.49 (s, 9H, 3 x CH_3_), 1.41 (s, 9H, 3 x CH_3_), 1.37 (d, *J* = 6.6 Hz, 3H, H-9), 1.31 (d, *J* = 6.2 Hz, 3H, *i*Pr-CH_3_''), 1.29 (d, *J* = 6.2 Hz, 3H, *i*Pr-CH_3_'), 1.12 (d, *J* = 6.4 Hz, 3H, Thr β-CH_3_); **^13^C NMR** (101 MHz, CDCl_3_) δ 170.4, 170.0, 169.7, 168.4, 156.0, 155.9, 155.0, 136.5, 128.6, 128.3, 128.2, 97.4, 95.9, 83.1, 80.3, 74.7, 72.6, 71.2, 70.6, 70.3, 67.3, 67.2, 58.9, 53.8, 48.8, 33.1, 28.4, 28.2, 21.7, 21.6, 21.3, 21.0, 16.4, 15.4; α only (*J*_C1-H3ax_ = 0 Hz); **HRMS** (ESI+) *m/z* Calc. for C_40_H_58_Cl_3_N_3_O_16_ [M + Na]^+^: 964.2775, found 964.2765.

***tert*-Butyl *N*-(tert-butoxycarbonyl)-*O*-(5-acetamido-4,8-di-*O*-acetyl-7-*N*-benzyloxycarb-onyl-1-isopropyl-α-pseudaminosyl)-L-threoninate (α‑S4)**

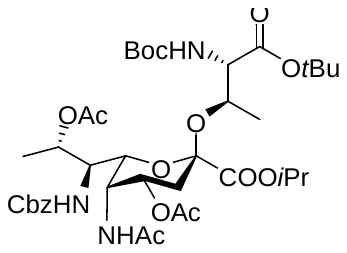

Compound **α‑S3** (143 mg, 151 µmol) and zinc powder (98.8 mg, 1.51 mmol, 10 equiv.) were dissolved in acetic acid (3 mL) and acetic anhydride (3 mL). The reaction mixture was stirred at 50 °C for 2 h. Upon completion, the reaction mixture was diluted with EtOAc (50 mL) and filtered through a pad of celite. The collected filtrate was washed with sat. aq. NaHCO_3_ solution (4x50 mL) and brine (1x50 mL). The organic layer was layer was collected, dried with anhydrous MgSO_4_ and concentrated *in vacuo*. The crude product was purified by flash column chromatography (eluent gradient: 0%-75% EtOAc in hexane) to afford the *title* compound **α‑S4** as a yellow foam (107 mg, 87%).

**[α]^25^_D_** = -61 (c = 1.7, CH_2_Cl_2_); **IR** (thin film) v*_max_* = 2980, 2933, 1720, 1534, 1503, 1455, 1368, 1309, 1230, 1152, 1090, 1048, 908, 842, 774, 737, 699 cm^-1^; **^1^H NMR** (500 MHz, CDCl_3_) δ 7.38 – 7.27 (m, 5H, Ar-H), 6.20 (d, J = 9.3 Hz, 1H, NH), 5.30 – 5.19 (m, 3H, H-8, 2 x NH), 5.13 (d, J = 12.1 Hz, 1H, PhC*H_2_*), 5.10 – 5.03 (m, 2H, H4, *i*Pr-CH), 4.96 (d, J = 12.1 Hz, 1H, PhC*H_2_*'), 4.59 (d, J = 9.5 Hz, 1H, H-5), 4.36 – 4.29 (m, 2H, H-7, Thr β-H), 4.18 (dd, J = 9.2, 2.6 Hz, 1H, Thr α-H), 4.06 – 4.02 (m, 1H, H-6), 2.19 (dd, J = 13.2, 5.0 Hz, 1H, H-3eq), 2.04 (s, 3H, CH_3_CO), 2.02 (s, 3H, CH_3_CO), 1.97 (s, 3H, CH_3_CO), 1.68 (t, J = 12.7 Hz, 1H, H-3ax), 1.49 (s, 9H, 3 x CH_3_), 1.43 (s, 9H, 3 x CH_3_), 1.35 (d, J = 6.6 Hz, 3H, H-9), 1.32 (d, J = 6.3 Hz, 3H, *i*Pr-CH_3_''), 1.29 (d, J = 6.2 Hz, 3H, *i*Pr-CH_3_'), 1.14 (d, J = 6.4 Hz, 3H, Thr β-CH_3_); **^13^C NMR** (126 MHz, CDCl_3_) δ 170.9, 170.7, 170.1, 169.7, 169.0, 156.1, 155.9, 136.4, 128.6, 128.3, 128.2, 97.2, 83.1, 80.2, 72.5, 70.8, 70.4, 67.3, 67.1, 58.9, 53.4, 46.1, 33.3, 28.4, 28.2, 23.4, 21.7, 21.6, 21.3, 21.0, 15.5, 15.4; **HRMS** (ESI+) *m/z* Calc. for C_39_H_59_N_3_O_15_ [M + Na]^+^: 832.3838, found 832.3827.

***tert*-Butyl *N*-(tert-butoxycarbonyl)-*O*-(5,7-diacetamido-4,8-di-*O*-acetyl-1-isopropyl-α-pseudaminosyl)-L-threoninate (α‑S5)**

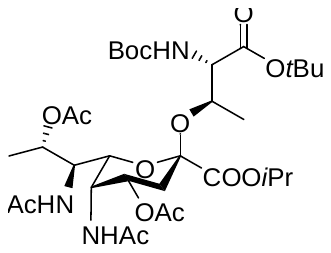

Compound **α‑S4** (162 mg, 203 µmol) was dissolved in MeOH (7 mL) before Pd/C (108 mg, 102 µmol, 0.5 equiv., 10 wt% palladium on activated carbon) was added. The reaction mixture was stirred under 1 atm H_2_ at rt for 2 h. Upon completion, the reaction mixture was filtered through a pad of celite, and the collected filtrate was concentrated *in vacuo*. The Cbz‑deprotected intermediate was dissolved in CH_2_Cl_2_ (10 mL) before triethylamine (142 µL, 1.02 mmol, 5.0 equiv.) and acetic anhydride (96.1 µL, 1.02 mmol, 5.0 equiv.) were added. The reaction mixture was stirred at rt for 1 h. Upon completion, the reaction mixture was diluted with CH_2_Cl_2_ (40 mL), washed with 1 M HCl (2x50 mL), sat. aq. NaHCO_3_ solution (1x50 mL) and brine (1x50 mL). The organic layer was collected, dried with anhydrous MgSO_4_ and concentrated *in vacuo*. The crude product was purified by flash column chromatography (eluent: 45% EtOAc and 5% MeOH in hexane) to afford the *title* compound **α‑S5** as a yellow oil (127 mg, 87% over 2 steps).

**[α]^25^_D_** = -66 (c = 1.1, CH_2_Cl_2_); **IR** (thin film) v*_max_* = 2997, 2931, 1741, 1726, 1669, 1549, 1506, 1457, 1370, 1275, 1237, 1156, 1104, 1050 cm^-1^; **^1^H NMR** (400 MHz, CDCl_3_) δ 6.21 (d, *J* = 9.2 Hz, 1H, NH), 6.02 (d, *J* = 10.4 Hz, 1H, NH), 5.23 (d, *J* = 9.4 Hz, 1H, NH), 5.17 (dd, *J* = 6.7, 2.8 Hz, 1H, H-8), 5.12 – 5.00 (m, 2H, H-4, *i*Pr-CH), 4.58 (td, *J* = 9.8, 2.7 Hz, 1H, H-7), 4.52 (d, *J* = 8.7 Hz, 1H, H-5), 4.36 (dd, *J* = 6.5, 2.7 Hz, 1H, Thr β-H), 4.20 (dd, *J* = 9.4, 2.6 Hz, 1H, Thr α-H), 4.10 (dd, *J* = 9.0, 2.0 Hz, 1H, H-6), 2.17 (dd, *J* = 13.1, 4.8 Hz, 1H, H-3eq), 2.05 (s, 3H, CH_3_CO), 2.00 (s, 3H, CH_3_CO), 1.95 (s, 3H, CH_3_CO), 1.94 (s, 3H, CH_3_CO), 1.67 (t, *J* = 12.8 Hz, 1H, H-3ax), 1.51 (s, 9H, 3 x CH_3_), 1.46 (s, 9H, 3 x CH_3_), 1.37 (d, *J* = 6.7 Hz, 3H, H-9), 1.32 (d, *J* = 6.3 Hz, 3H, *i*Pr-CH_3_''), 1.29 (d, *J* = 6.2 Hz, 3H, *i*Pr-CH_3_'), 1.15 (d, *J* = 6.3 Hz, 3H, Thr β-CH_3_); **^13^C NMR** (101 MHz, CDCl_3_) δ 171.1, 170.8, 170.2, 169.9, 169.8, 168.9, 156.0, 97.4, 83.1, 80.2, 72.4, 71.2, 70.8, 70.4, 67.1, 58.8, 51.1, 46.1, 33.3, 29.8, 28.5, 28.2, 23.3, 21.7, 21.6, 21.4, 21.0, 16.1, 15.5; **HRMS** (ESI+) *m/z* Calc. for C_33_H_55_N_3_O_14_ [M + Na]^+^: 740.3576, found 740.3568.

***N*-(allyloxycarbonyl)-*O*-(5,7-diacetamido-4,8-di-*O*-acetyl-1-isopropyl-α-pseudaminosyl)-L-threonine (α‑S6)**

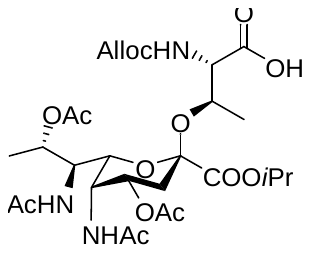

Compound **α‑S5** (66.0 mg, 91.9 µmol) was dissolved in a solution of TFA/water (4 mL, 19:1 v/v). The reaction was stirred at rt for 2 h. Upon completion, the reaction mixture was concentrated *in vacuo*. This dried intermediate was redissolved in THF (5 mL) and sat. aq. NaHCO_3_ (5 mL). Allyl chloroformate (9.8 µL, 91.9 µmol, 1.0 equiv.) was added and the reaction mixture was stirred vigorously at rt for 12 h. Upon completion, the reaction mixture was extracted with EtOAc (40 mL). The organic layer was layer was collected, dried with anhydrous MgSO_4_ and concentrated *in vacuo*. The crude product was purified by flash column chromatography (eluent gradient: 0%-10% MeOH in CH_2_Cl_2_ with 1% acetic acid as additive) to afford the *title* compound **α‑S6** as a yellow oil (51.7 mg, 87% over 2 steps).

**[α]^25^_D_** = -49 (c = 1.3, MeOH); **IR** (thin film) v*_max_* = 3284, 3066, 2984, 2936, 1721, 1665, 1547, 1433, 1374, 1244, 1203, 1178, 1134, 1103, 1043, 976, 957, 836, 800, 775, 750, 720, 668, 604 cm^-1^; **^1^H NMR** (400 MHz, DMSO-*d*_6_) δ 7.91 (d, *J* = 9.5 Hz, 1H, NH), 7.72 (d, *J* = 10.1 Hz, 1H, NH), 6.65 (s, 1H, NH), 5.90 (ddt, *J* = 17.3, 10.6, 5.3 Hz, 1H, CH_2_C*H*=CH_2_), 5.29 (dd, *J* = 17.1, 2.0 Hz, 1H, CH_2_CH=C*H_2_*''), 5.17 (dd, *J* = 10.5, 1.7 Hz, 1H, CH_2_CH=C*H_2_*'), 5.12 (dd, *J* = 6.6, 2.9 Hz, 1H, H-8), 5.03 (td, *J* = 8.7, 3.8 Hz, 1H, H-4), 4.94 (p, *J* = 6.2 Hz, 1H, *i*Pr-CH), 4.48 (dd, *J* = 5.3, 1.5 Hz, 2H, C*H_2_*CH=CH_2_), 4.38 (dt, *J* = 9.9, 3.3 Hz, 1H, H-5), 4.25 – 4.16 (m, 2H, Thr β-H, H-7), 4.09 (d, *J* = 9.4 Hz, 1H, H-6), 3.93 (d, *J* = 8.0 Hz, 1H, Thr α-H), 1.94 (d, *J* = 8.7 Hz, 2H, H-3), 1.90 (s, 3H, CH_3_CO), 1.83 (s, 3H, CH_3_CO), 1.78 (s, 3H, CH_3_CO), 1.73 (s, 3H, CH_3_CO), 1.25 (d, *J* = 3.0 Hz, 3H, *i*Pr-CH_3_''), 1.23 (d, *J* = 3.0 Hz, 3H, *i*Pr-CH_3_'), 1.20 (d, *J* = 6.6 Hz, 3H, H-9), 1.03 (d, *J* = 6.2 Hz, 3H, Thr β-CH_3_); **^13^C NMR** (101 MHz, DMSO-*d*_6_) δ 169.9, 169.5, 169.3, 168.9, 167.1, 158.1, 157.8, 155.8, 133.6, 118.8, 116.9, 115.8, 96.5, 70.9, 70.1, 69.3, 68.8, 67.0, 64.5, 59.6, 50.3, 44.9, 32.0, 22.7, 22.5, 21.4, 21.2, 21.0, 20.7, 16.3, 15.1. **HRMS** (ESI+) *m/z* Calc. for C_28_H_43_N_3_O_14_ [M + Na]^+^: 668.2637, found 668.2633.

***tert*-Butyl *N*-(tert-butoxycarbonyl)-*O*-(4,8-di-*O*-acetyl-5-azido-7-*N*-benzyloxycarbonyl-1-isopropyl-β-pseudaminosyl)-L-threoninate (β‑S3)**

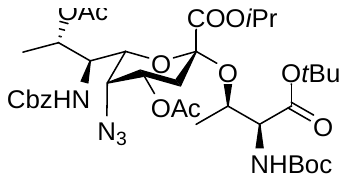

PseN_3_ adamantyl thioglycoside donor **4** (275 mg, 0.40 mmol) was glycosylated with commercially available Boc-Thr(OH)-O*t*Bu acceptor (221 mg, 0.80 mmol, 2.0 equiv.) according to general procedure 2. The crude product was purified by flash column chromatography (eluent gradient: 0%-25% EtOAc in hexane) to afford the *title* compound **β‑S3** as a yellow foam (282 mg, 89%).

**[α]^25^_D_** = -25 (c = 1.0, CH_2_Cl_2_); **IR** (thin film) v*_max_* = 2979, 2925, 2111, 1733, 1532, 1500, 1456, 1368, 1300, 1230, 1182, 1154, 1094, 1057, 1022, 1002, 925, 846, 743, 699 cm^-1^; **^1^H NMR**  (500 MHz, CDCl_3_) δ 7.40 – 7.28 (m, 5H, Ar-H), 5.25 (dd, *J* = 6.7, 4.1 Hz, 1H, H-8), 5.19 (d, *J* = 8.7 Hz, 1H, NH), 5.16 – 5.04 (m, 3H, PhC*H_2_*, *i*Pr-CH), 4.98 (d, *J* = 10.0 Hz, 1H, NH), 4.82 (dt, *J* = 12.7, 3.8 Hz, 1H, H-4), 4.44 (td, *J* = 9.8, 3.9 Hz, 1H, H-7), 4.25 – 4.15 (m, 1H, Thr β-H), 4.02 (dd, *J* = 8.9, 4.1 Hz, 1H, Thr α-H), 3.90 – 3.82 (m, 1H, H-5), 3.64 (d, *J* = 7.9 Hz, 1H, H-6), 2.39 (dd, *J* = 12.5, 4.1 Hz, 1H, H-3eq), 2.15 (t, *J* = 12.7 Hz, 1H, H-3ax), 2.11 (s, 3H, CH_3_CO), 1.98 (s, 3H, CH_3_CO), 1.45 (s, 9H, 3 x CH_3_), 1.44 (s, 9H, 3 x CH_3_), 1.36 (d, *J* = 6.3 Hz, 3H, *i*Pr-CH_3_''), 1.34 (d, *J* = 6.7 Hz, 3H, 15, *i*Pr-CH_3_'), 1.33 – 1.29 (m, 6H, H-9, Thr β-CH_3_); **^13^C NMR** (126 MHz, CDCl_3_) δ 170.2, 170.0, 169.6, 166.9, 156.3, 156.0, 136.4, 128.7, 128.4, 128.4, 100.7, 81.9, 79.8, 73.6, 72.2, 71.2, 69.7, 69.2, 67.3, 59.6, 58.6, 53.9, 33.4, 28.5, 28.1, 21.9, 21.8, 21.2, 20.8, 19.2, 14.3; β only (*J*_C1-H3ax_ = 6.3 Hz); **HRMS** (ESI+) *m/z* Calc. for C_37_H_55_N_5_O_14_ [M + Na]^+^: 816.3638, found 816.3634.

***tert*-Butyl *N*-(*tert*-butoxycarbonyl)-*O*-(5,7-diacetamido-4,8-di-*O*-acetyl-1-isopropyl-α-pseudaminosyl)-L-threoninate (β‑S5)**

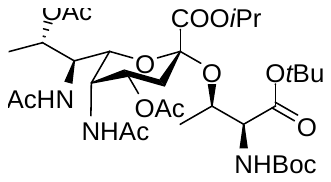

Compound **β‑S3** (131 mg, 165 µmol) was dissolved in MeOH (7 mL) before Pd/C (87.8 mg, 82.5 µmol, 0.5 equiv., 10 wt% palladium on activated carbon) was added. The reaction mixture was stirred under 1 atm H_2_ at rt for 2 h. Upon completion, the reaction mixture was filtered through a pad of celite, and the collected filtrate was concentrated *in vacuo*. The free amine intermediate was dissolved in CH_2_Cl_2_ (7 mL) before triethylamine (115 µL, 823 µmol, 5.0 equiv.) and acetic anhydride (77.8 µL, 823 µmol, 5.0 equiv.) were added. The reaction mixture was stirred at rt for 1 h. Upon completion, the reaction mixture was diluted with CH_2_Cl_2_ (40 mL), washed with 1 M HCl (2x50 mL), sat. aq. NaHCO_3_ solution (1x50 mL) and brine (1x50 mL). The organic layer was collected, dried with anhydrous MgSO_4_ and concentrated *in vacuo*. The crude product was purified by flash column chromatography (eluent: 45% EtOAc and 5% MeOH in hexane) to afford the *title* compound **β‑S5** as a yellow oil (56.3 mg, 48% over 2 steps).

**[α]^25^_D_** = -57 (c = 1.2, CH_2_Cl_2_); **IR** (thin film) v*_max_* = 2979, 2925, 2853, 1723, 1664, 1544, 1508, 1457, 1368, 1256, 1236, 1152, 1102, 1088, 1047, 972, 844, 736, 683 cm^-1^; **^1^H NMR** (500 MHz, CDCl_3_) δ 6.46 (d, *J* = 9.2 Hz, 1H, NH), 5.69 (s, 1H, NH), 5.32 – 5.26 (m, 2H, H-8, NH), 5.13 (p, *J* = 6.3 Hz, 1H, *i*Pr-CH), 4.76 (dd, *J* = 10.7, 6.5 Hz, 1H, H-4), 4.57 (dt, *J* = 11.8, 5.9 Hz, 1H, H-7), 4.48 (d, *J* = 9.5 Hz, 1H, H-5), 4.39 (dd, *J* = 7.7, 3.9 Hz, 1H, Thr α-H), 4.26 (t, *J* = 5.5 Hz, 1H, Thr β-H), 3.88 (d, *J* = 10.0 Hz, 1H, H-6), 2.33 (dd, *J* = 13.1, 3.7 Hz, 1H, H-3eq), 2.06 (s, 3H, CH_3_CO), 2.02 – 1.97 (m, 7H, H-3ax, 2 x CH_3_CO), 1.95 (s, 3H, CH_3_CO), 1.46 (s, 9H, 3 x CH_3_), 1.46 (s, 9H, 3 x CH_3_), 1.36 (d, *J* = 6.3 Hz, 3H, *i*Pr-CH_3_''), 1.34 (d, *J* = 6.0 Hz, 3H, *i*Pr-CH_3_'), 1.32 (d, *J* = 5.6 Hz, 3H, H-9), 1.28 (d, *J* = 6.3 Hz, 3H, Thr β-CH_3_); **^13^C NMR** (126 MHz, CDCl_3_) δ 171.9, 170.6, 170.4, 169.7, 166.8, 155.8, 101.0, 82.2, 80.0, 72.5, 71.0, 69.7, 68.0, 59.1, 50.6, 45.3, 31.8, 28.5, 28.1, 23.3, 21.9, 21.9, 21.8, 21.3, 21.1, 18.0, 14.0. **HRMS** (ESI+) *m/z* Calc. for C_33_H_55_N_3_O_14_ [M + Na]^+^: 740.3576, found 740.3561.

***N*-(allyloxycarbonyl)-*O*-(5,7-diacetamido-4,8-di-*O*-acetyl-1-isopropyl-β-pseudaminosyl)-L-threonine (β‑S6)**

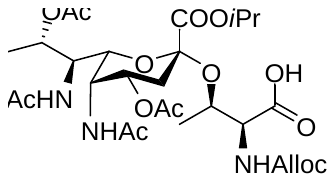

Compound **β‑S5** (46.0 mg, 64.1 µmol) was dissolved in a solution of TFA/water (2 mL, 19:1 v/v). The reaction was stirred at rt for 2 h. Upon completion, the reaction mixture was concentrated *in vacuo*. The resulting residue was redissolved in THF (2.5 mL) and sat. aq. NaHCO_3_ (2.5 mL) added. Allyl chloroformate (6.8 µL, 64.1 µmol, 1.0 equiv.) was added and the reaction mixture was stirred vigorously at rt for 12 h. Upon completion, the reaction mixture was extracted with EtOAc (20 mL). The organic layer was collected, dried with anhydrous MgSO_4_ and concentrated *in vacuo*. The crude product was purified by flash column chromatography (eluent gradient: 0%-10% MeOH in CH_2_Cl_2_ with 1% acetic acid as additive) to afford the *title* compound **β‑S6** as a yellow oil (31.5 mg, 76% over 2 steps).

**[α]^25^_D_** = -46 (c = 1.2, MeOH); **IR** (thin film) v*_max_* = 3337, 3081, 2984, 2939, 1732, 1663, 1544, 1445, 1373, 1238, 1204, 1183, 1149, 1102, 1063, 1045 cm^-1^; **^1^H NMR** (500 MHz, DMSO-*d*_6_) δ 12.84 (s, 1H, COOH), 7.76 (d, *J* = 10.0 Hz, 1H, NH), 7.59 (d, *J* = 9.8 Hz, 1H, NH), 6.57 (s, 1H, NH), 5.96 – 5.84 (m, 1H, CH_2_C*H*=CH_2_), 5.31 (dd, *J* = 17.3, 2.0 Hz, 1H, CH_2_CH=C*H_2_*''), 5.24 – 5.11 (m, 2H, H8, CH_2_CH=C*H_2_*'), 5.03 (p, *J* = 6.2 Hz, 1H, *i*Pr-CH), 4.61 (dt, *J* = 13.3, 4.2 Hz, 1H, H-4), 4.49 (dt, *J* = 5.3, 1.6 Hz, 2H, C*H_2_*CH=CH_2_), 4.39 – 4.27 (m, 3H, H-5, H-7, Thr β-H), 4.03 – 3.97 (m, 1H, Thr α-H), 3.64 (dd, *J* = 10.4, 2.3 Hz, 1H, H-6), 2.16 (dd, *J* = 12.2, 4.3 Hz, 1H, H-3eq), 1.98 (t, *J* = 12.8 Hz, 1H, H-3ax), 1.87 (s, 3H, CH_3_CO), 1.85 (s, 3H, CH_3_CO), 1.79 (s, 3H, CH_3_CO), 1.68 (s, 3H, CH_3_CO), 1.29 (d, *J* = 5.2 Hz, 3H, Thr β-CH_3_), 1.28 (d, *J* = 6.2 Hz, 3H, *i*Pr-CH_3_''), 1.25 (d, *J* = 6.3 Hz, 3H, *i*Pr-CH_3_'), 1.21 (d, *J* = 6.6 Hz, 3H, H-9); **^13^C NMR** (126 MHz, DMSO-*d*_6_) δ 171.1, 170.1, 169.5, 169.3, 168.8, 166.6, 155.9, 133.4, 117.1, 100.3, 72.3, 72.0, 70.1, 69.1, 67.2, 64.7, 58.8, 49.2, 44.0, 32.8, 22.7, 22.5, 21.2, 21.0, 20.7, 19.4, 13.3; **HRMS** (ESI+) *m/z* Calc. for C_28_H_43_N_3_O_14_ [M + Na]^+^: 668.2637, found 668.2614.

**FlaA(206-211): H_2_N-VS(α-Pse5Ac7Ac)SSAG-OH.formate salt (α‑8)**

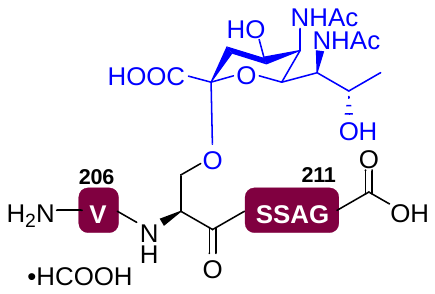

Fmoc-Gly‑OH (39.5 mg, 133 µmol, 0.5 equiv.) was loaded onto 2-CTC resin (200 mg, manufacturer’s resin loading: 1.33 mmol/g) according to general procedure 3 (final resin loading: 0.45 mmol/g). Iterative coupling of the FlaA S208-A210 fragment was performed on 30 µmol of resin-bound Fmoc-protected glycine according to general procedure 4. Coupling of pseudaminylated serine building block **α‑1** was then performed according to general procedure 5. Afterwards, the final Fmoc‑Val‑OH amino acid was coupled following general procedure 4, then Fmoc deprotected by treating the resin with 20% (v/v) piperidine in DMF (2x5 mL) for 5 min, and subsequently washed with DMF (5x5 mL) and CH_2_Cl_2_ (10x5 mL). The resulting peptide was then cleaved off from the resin and side chain deprotected according to general procedure 6 to generate the crude peptide. Finally, this peptide was saponified according to general procedure 7, and then purified by preparative RP-HPLC (column: Waters Sunfire C_18_ 5 μm 130 Å (19 x 150 mm), flow rate: 15 mL/min, gradient: 0%-30% B in A over 45 min). The product was lyophilised to afford the *title* compound **α‑8** as a white fluffy solid (3.6 mg, 21% over 11 steps). **LRMS** (ESI+) *m/z* [M + H]^+^: 823.50, [M + 2H]^2+^: 412.30, [2M + H]^+^: 1645.95.

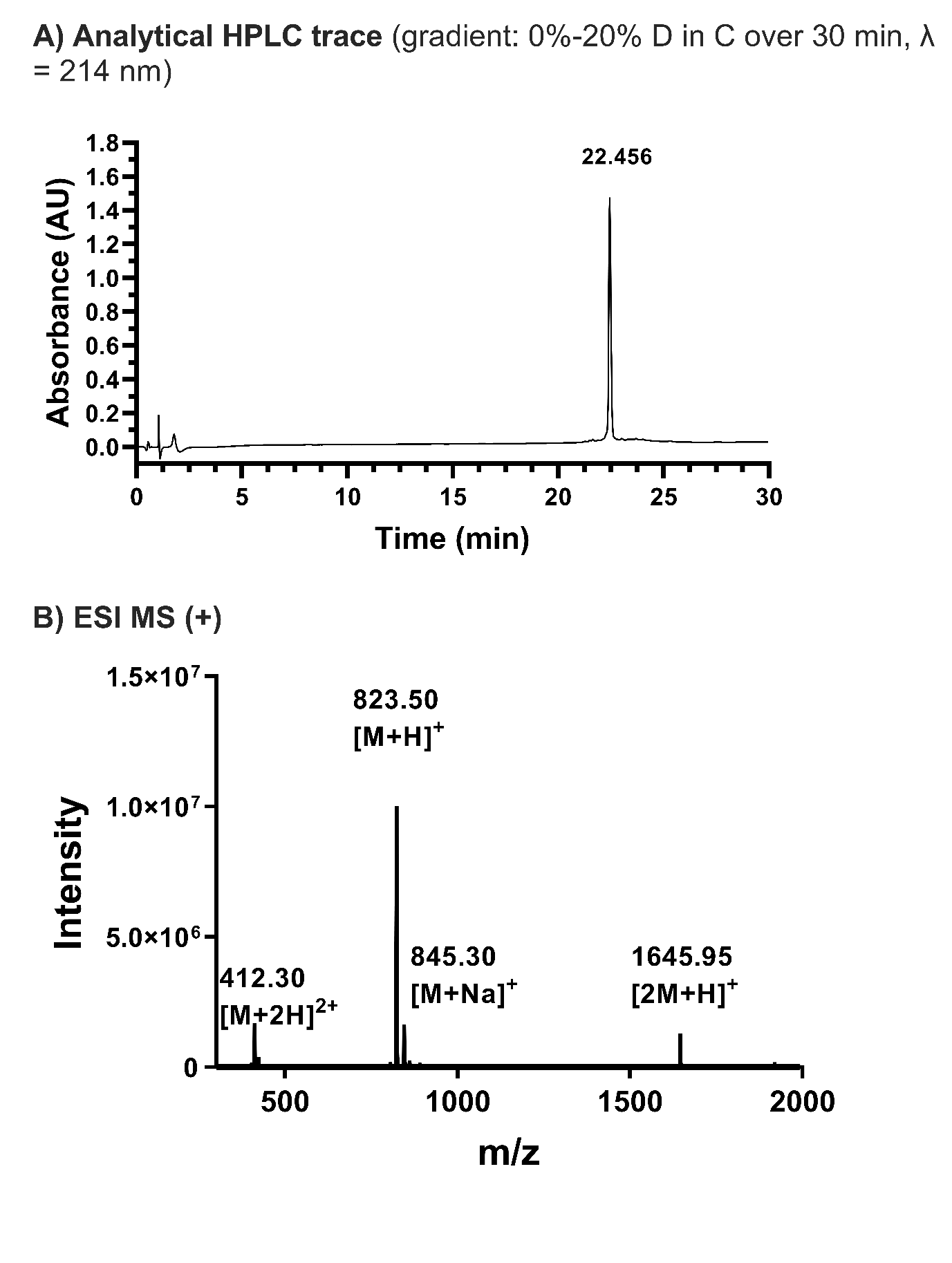

**FlaA(206-211): H_2_N-VS(β-Pse5Ac7Ac)SSAG-OH.formate salt (β‑8)**

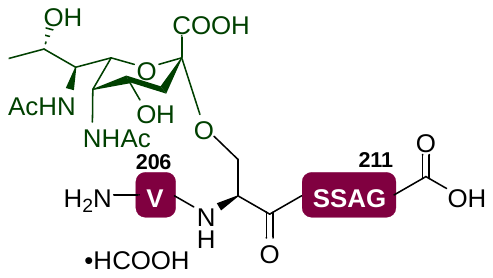

Iterative coupling of the FlaA S208-A210 fragment was performed on 30 µmol of resin-bound Fmoc-protected glycine (*vide supra*) (30 µmol) according to general procedure 4. Coupling of pseudaminylated serine building block **β‑1** was then performed according to general procedure 5. Afterwards, the final Fmoc‑Val‑OH amino acid was coupled following general procedure 4, then Fmoc deprotected by treating the resin with 20% (v/v) piperidine in DMF (2x5 mL) for 5 min, and subsequently washed with DMF (5x5 mL) and CH_2_Cl_2_ (10x5 mL). The resulting peptide was then cleaved off from the resin and side chain deprotected according to general procedure 6 to generate the crude peptide. Finally, this peptide was saponified according to general procedure 7, and then purified by preparative RP-HPLC (column: Waters Sunfire C_18_ 5 μm 130 Å (19 x 150 mm), flow rate: 15 mL/min, gradient: 0%-30% B in A over 45 min). The product was lyophilised to afford the *title* compound **β‑8** as a white fluffy solid (4.0 mg, 23% over 11 steps). **LRMS** (ESI+) *m/z* [M + H]^+^: 823.45, [M + 2H]^2+^: 412.15, [2M + H]^+^: 1646.30.

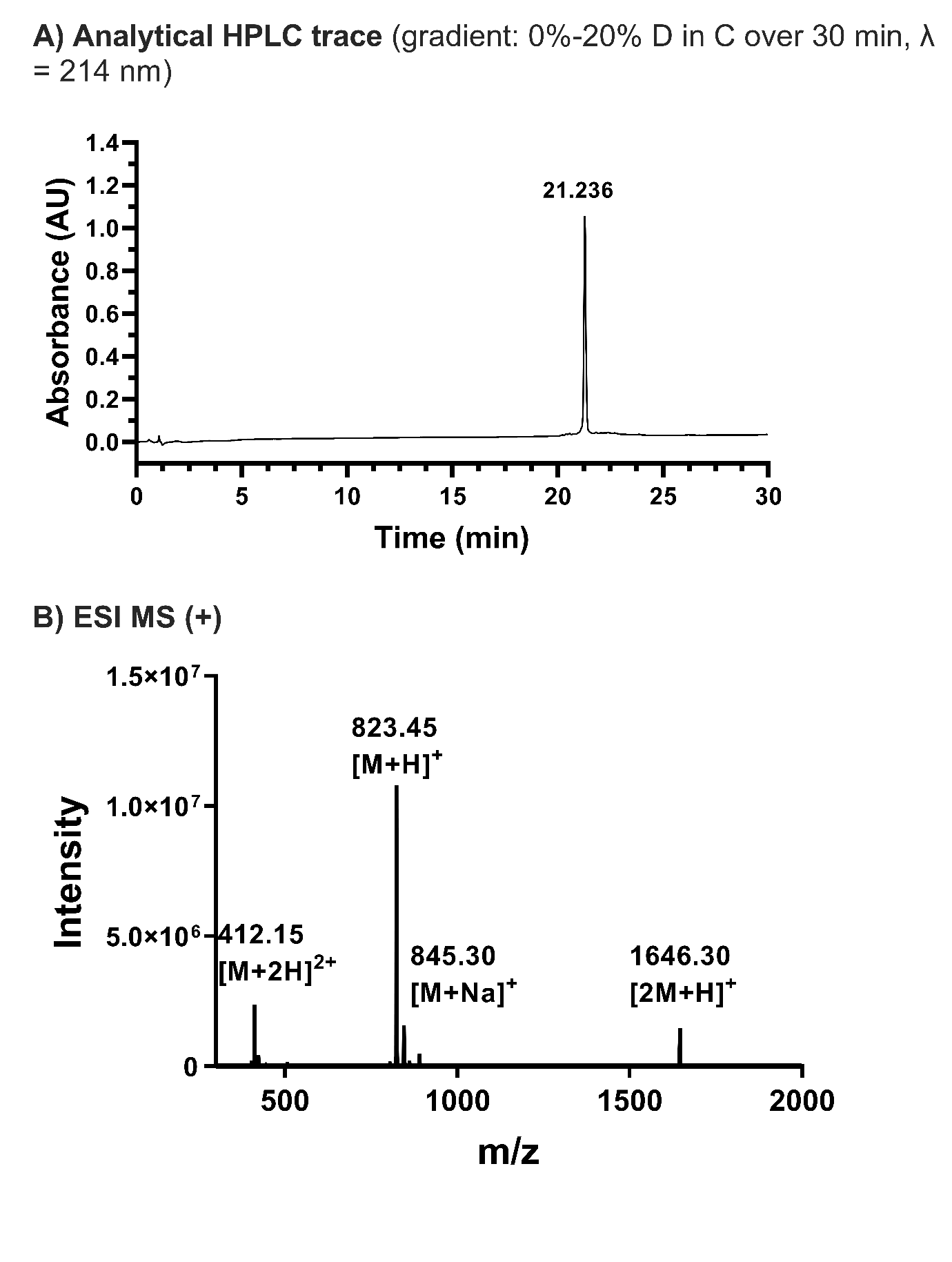

**FlaB(242-260): H_2_N-ASYNVMATGGTPVQS(α-Pse5Ac7Ac)GTVR-OH.diformate salt (α‑9)**

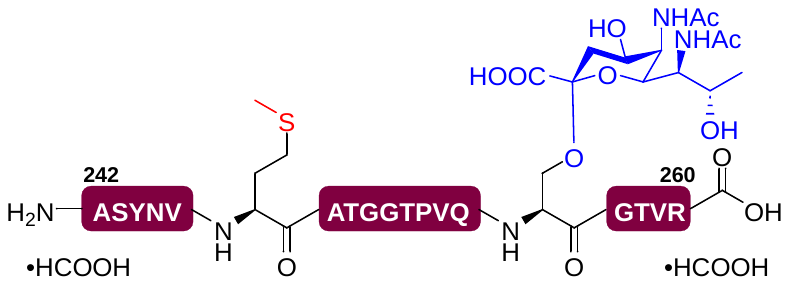

Fmoc-Arg(Pbf)‑OH (424 mg, 654 µmol, 0.5 equiv.) was loaded onto 2-CTC resin (1000 mg, manufacturer’s resin loading: 1.33 mmol/g) according to general procedure 3 (final resin loading: 0.29 mmol/g). Iterative coupling of the G257-V259 fragment was performed on resin‑bound Fmoc‑protected arginine (30 µmol) according to general procedure 4. Coupling of pseudaminylated serine building block **α‑1** was then performed according to general procedure 5. Afterwards, fragment FlaB A242-Q255 was completed following general procedure 4. After the final coupling, the peptide was Fmoc deprotected by treating the resin with 20% (v/v) piperidine in DMF (2x5 mL) for 5 min, then washed with DMF (5x5 mL) and CH_2_Cl_2_ (10x5 mL). The resulting peptide was then cleaved off from the resin and side chain deprotected according to general procedure 6 to generate the crude peptide. Finally, this peptide was saponified according to general procedure 7, and then purified by preparative RP-HPLC (column: Waters Sunfire C_18_ 5 μm 130 Å (19 x 150 mm), flow rate: 15 mL/min, gradient: 0%-40% B in A over 40 min). The product was lyophilised to afford the *title* compound **α‑9** as a white fluffy solid (1.6 mg, 2% over 39 steps). **LRMS** (ESI+) *m/z* [M + 2H]^2+^: 1106.90, [M + 3H]^3+^: 738.25, [2M + 3H]^3+^: 1475.85.

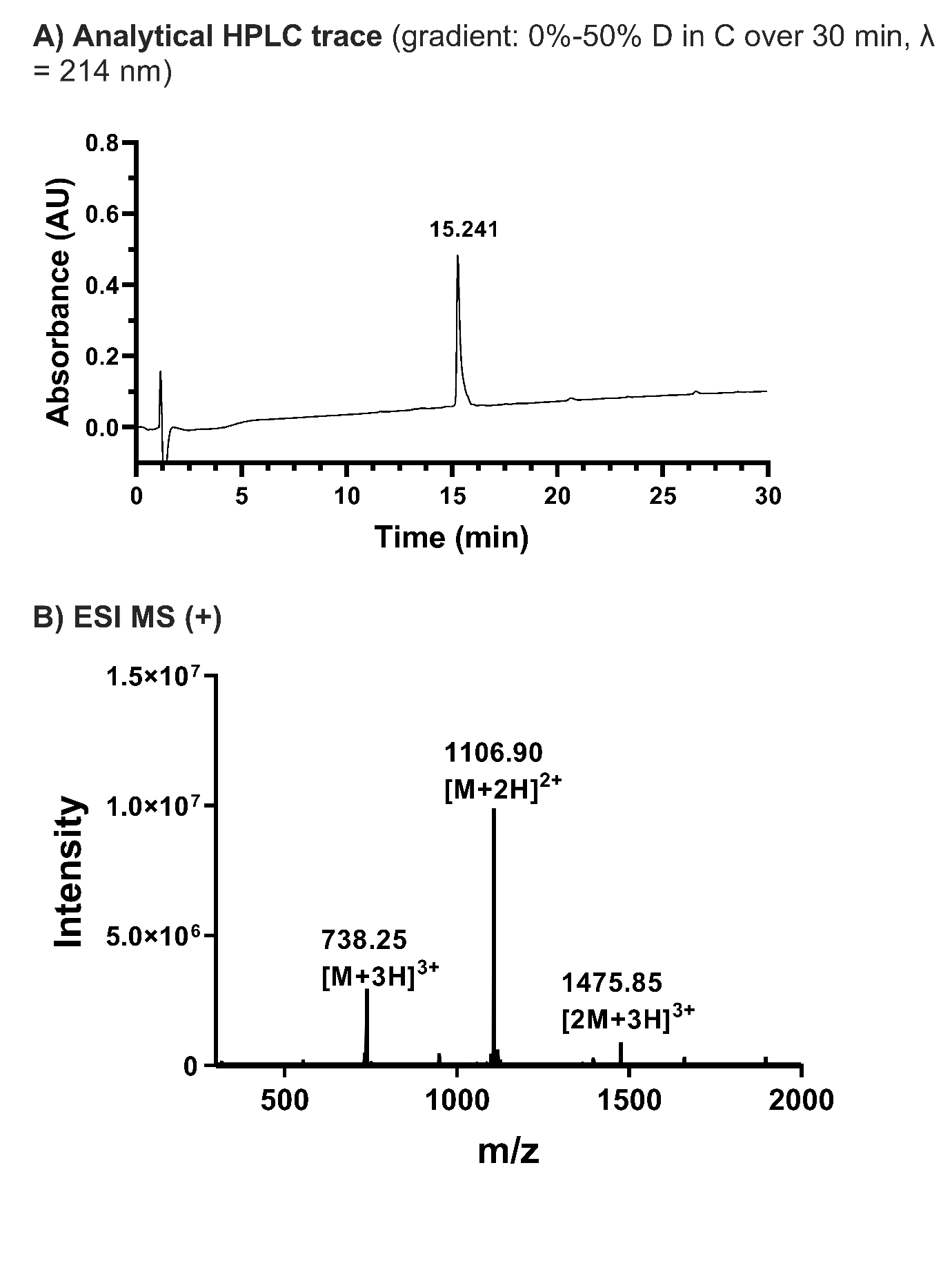

**FlaB(242-260): H_2_N-ASYNVMATGGTPVQS(β-Pse5Ac7Ac)GTVR-OH.diformate salt (β‑9)**

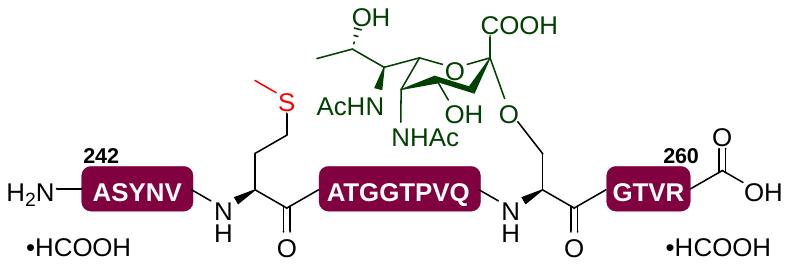

Iterative coupling of the FlaB G257-V259 fragment was performed on resin‑bound Fmoc‑protected arginine (*vide supra*) (30 µmol) according to general procedure 4. Coupling of pseudaminylated serine building block **β‑1** was then performed according to general procedure 5. Afterwards, fragment FlaB A242-Q255 was completed following general procedure 4. After the final coupling, the peptide was Fmoc deprotected by treating the resin with 20% (v/v) piperidine in DMF (2x5 mL) for 5 min, then washed with DMF (5x5 mL) and CH_2_Cl_2_ (10x5 mL). The resulting peptide was then cleaved off from the resin and side chain deprotected according to 6 to generate the crude peptide. Finally, this peptide was saponified according to general procedure 7, and then purified by preparative RP-HPLC (column: Waters Sunfire C_18_ 5 μm 130 Å (19 x 150 mm), flow rate: 15 mL/min, gradient: 0%-40% B in A over 40 min). The product was lyophilised to afford the *title* compound **β‑9** as a white fluffy solid (1.2 mg, 2% over 39 steps). **LRMS** (ESI+) *m/z* [M + 2H]^2+^: 1106.95, [M + 3H]^3+^: 738.35, [2M + 3H]^3+^: 1475.80.

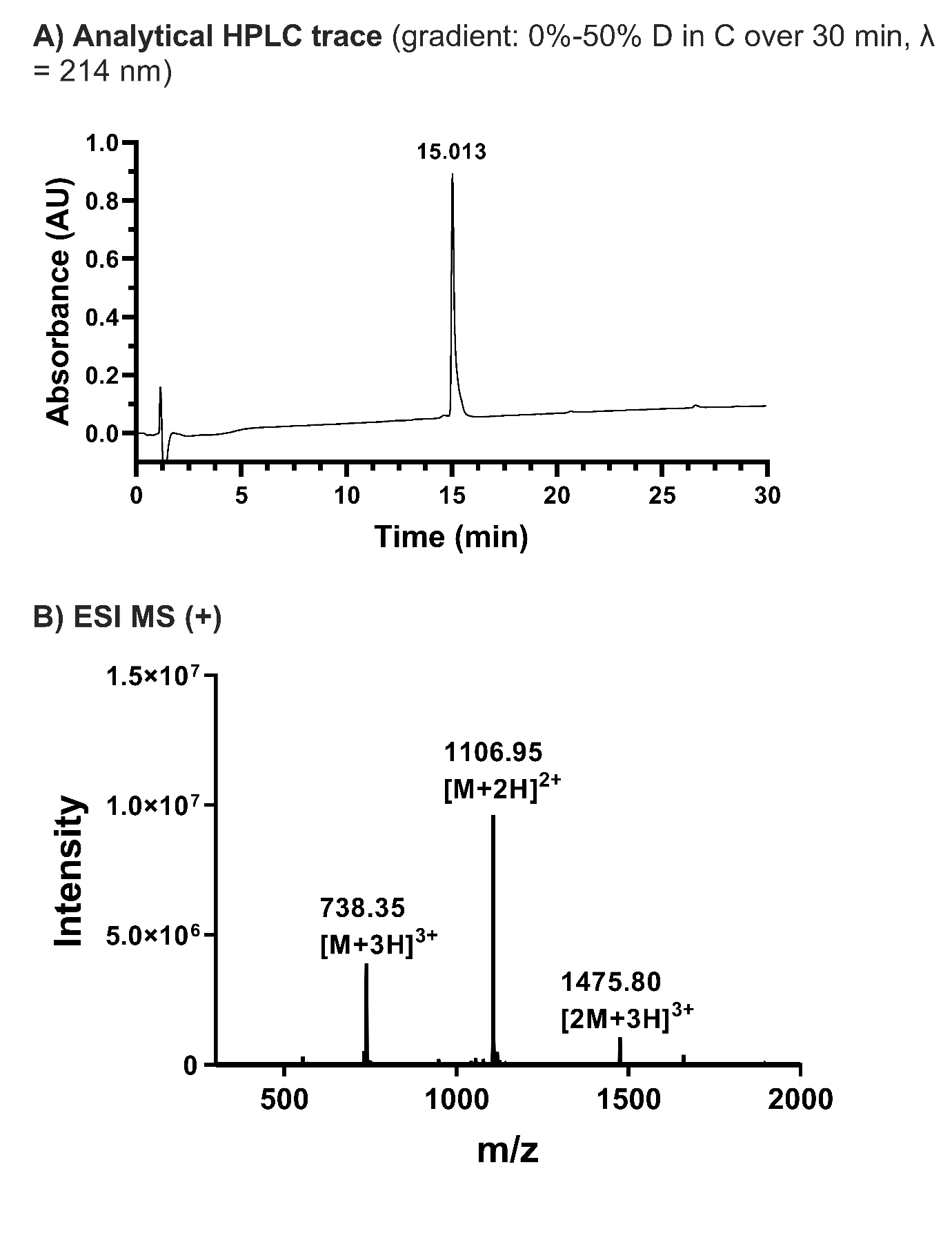

**FlaB(242-260): H_2_N-ASYNVM(O)ATGGTPVQS(α-Pse5Ac7Ac)GTVR-OH.diformate salt (α‑9_ox_)**

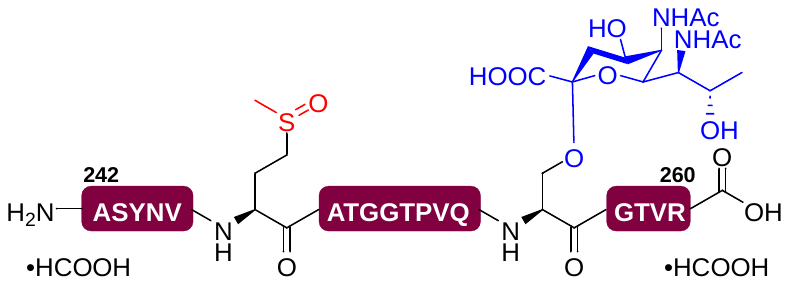

Iterative coupling of the FlaB G257-V259 fragment was performed on resin‑bound Fmoc‑protected arginine (*vide supra*) (30 µmol) according to general procedure 4. Coupling of pseudaminylated serine building block **α‑1** was then performed according to general procedure 5. Afterwards, fragment FlaB A242-Q255 was completed following general procedure 4. Residue 247 was coupled with the non‑canonical Fmoc‑Met(O)‑OH. After the final coupling, the peptide was Fmoc deprotected by treating the resin with 20% (v/v) piperidine in DMF (2x5 mL) for 5 min, then washed with DMF (5x5 mL) and CH_2_Cl_2_ (10x5 mL). The resulting peptide was then cleaved off from the resin and side chain deprotected according to general procedure 6 to generate the crude peptide. Finally, this peptide was saponified according to general procedure 7, and then purified by preparative RP-HPLC (column: Waters Sunfire C_18_ 5 μm 130 Å (19 x 150 mm), flow rate: 15 mL/min, gradient: 0%-40% B in A over 40 min). The product was lyophilised to afford the *title* compound **α‑9_ox_** as a white fluffy solid (0.8 mg, 1% over 39 steps). **LRMS** (ESI+) *m/z* [M + 2H]^2+^: 1114.95, [M + 3H]^3+^: 743.70, [2M + 3H]^3+^: 1486.35.

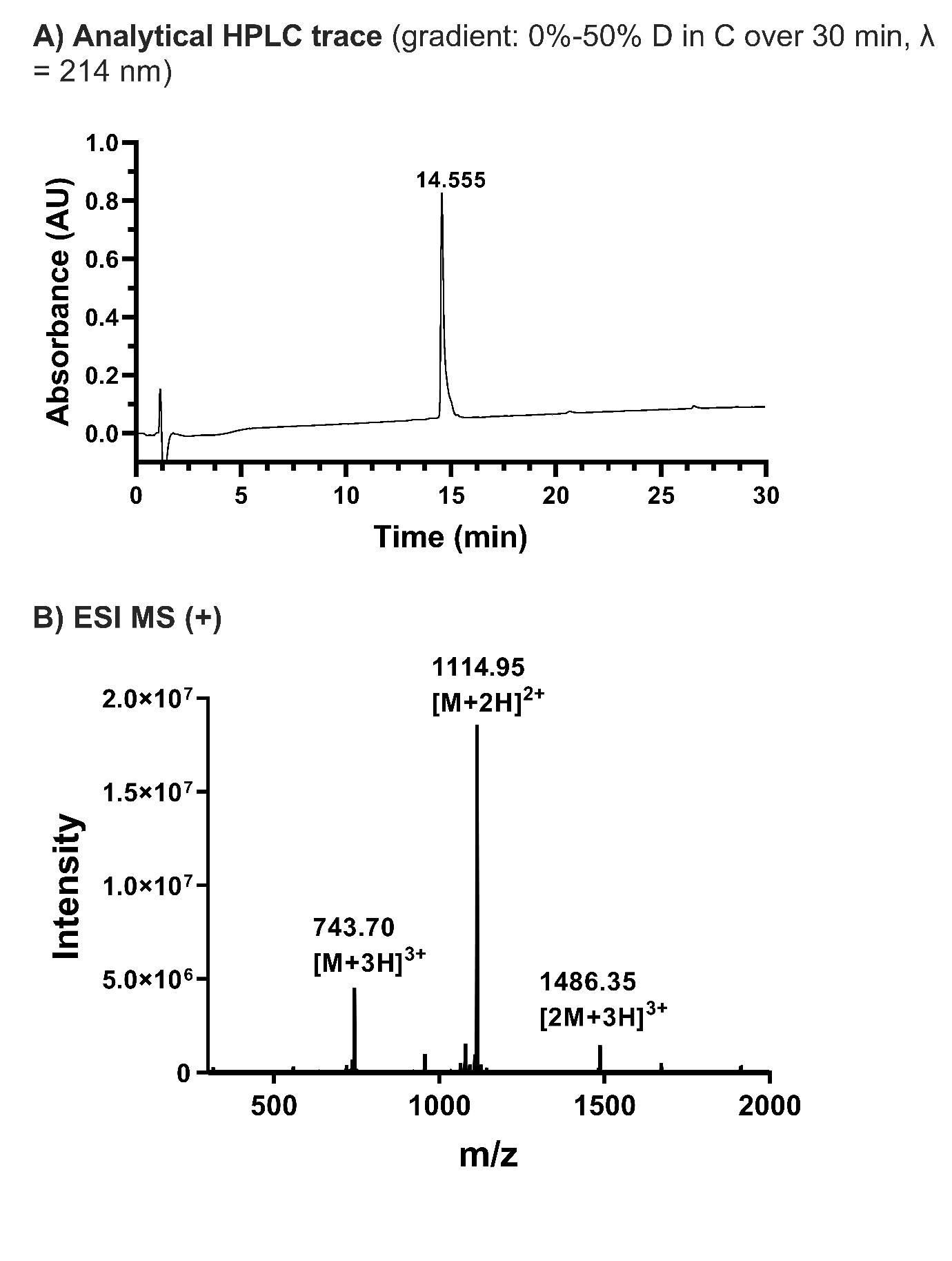

**FlaB(242-260): H_2_N-ASYNVM(O)ATGGTPVQS(β-Pse5Ac7Ac)GTVR-OH.diformate salt (β‑9_ox_)**

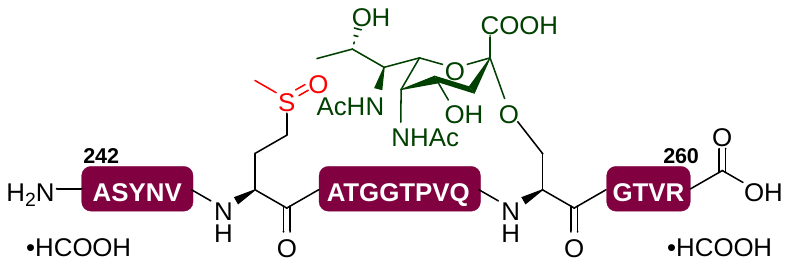

Iterative coupling of the FlaB G257-V259 fragment was performed on resin‑bound Fmoc‑protected arginine (*vide supra*) (30 µmol) according to general procedure 4. Coupling of pseudaminylated serine building block **β‑1** was then performed according to general procedure 5. Afterwards, fragment FlaB A242-Q255 was completed following general procedure 4. Residue 247 was coupled with the non‑canonical Fmoc‑Met(O)‑OH. After the final coupling, the peptide was Fmoc deprotected by treating the resin with 20% (v/v) piperidine in DMF (2x5 mL) for 5 min, then washed with DMF (5x5 mL) and CH_2_Cl_2_ (10x5 mL). The resulting peptide was then cleaved off from the resin and side chain deprotected according to general procedure 6 to generate the crude peptide. Finally, this peptide was saponified according to general procedure 7, and then purified by preparative RP-HPLC (column: Waters Sunfire C_18_ 5 μm 130 Å (19 x 150 mm), flow rate: 15 mL/min, gradient: 0%-40% B in A over 40 min). The product was lyophilised to afford the *title* compound **β‑9_ox_** as a white fluffy solid (0.5 mg, 1% over 39 steps). **LRMS** (ESI+) *m/z* [M + 2H]^2+^: 1114.95, [M + 3H]^3+^: 743.65, [2M + 3H]^3+^: 1486.50.

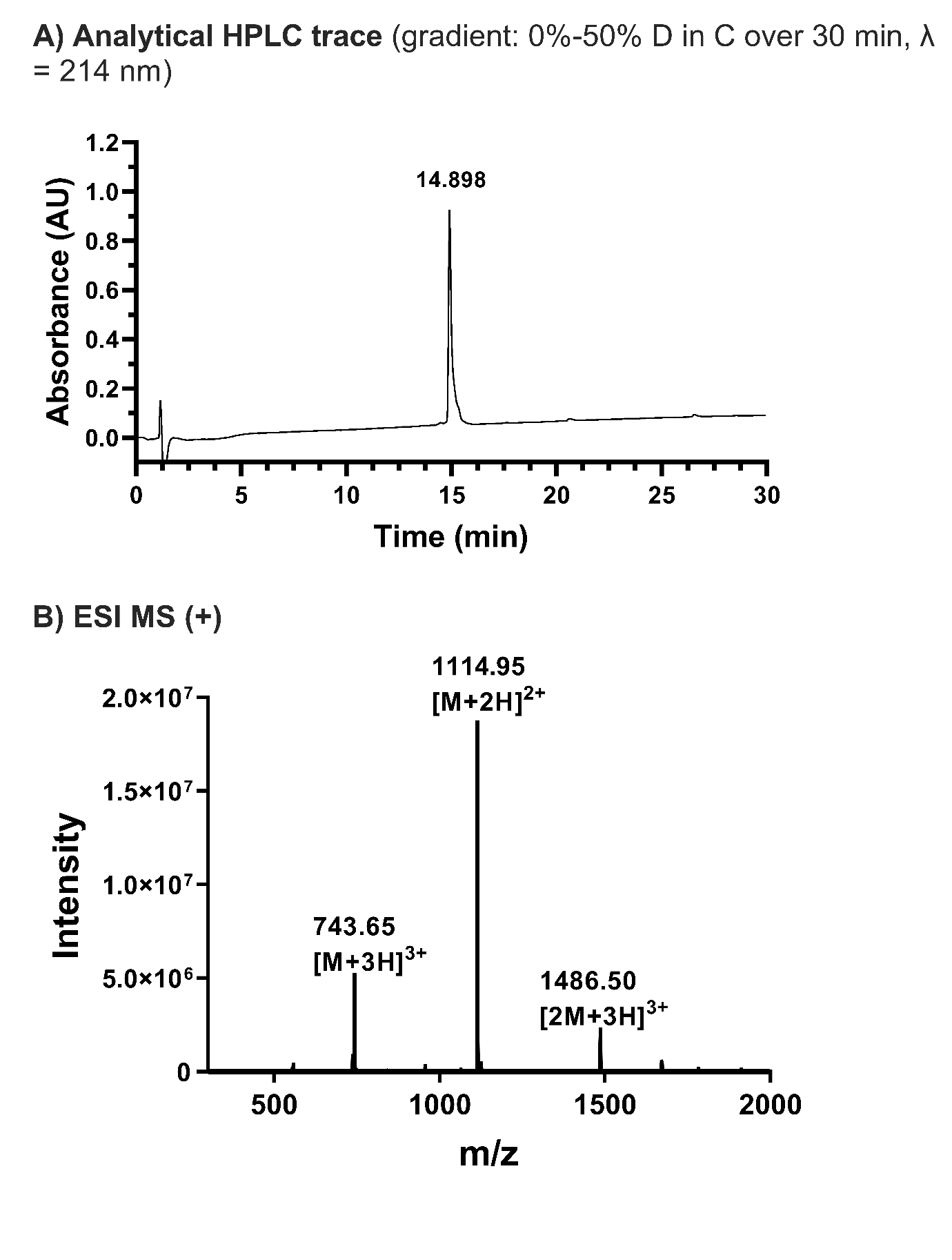

**FlgE(281-291): H_2_N-ISFTNDS(α-Pse5Ac7Ac)AVSR-OH.diformate salt (α‑10)**

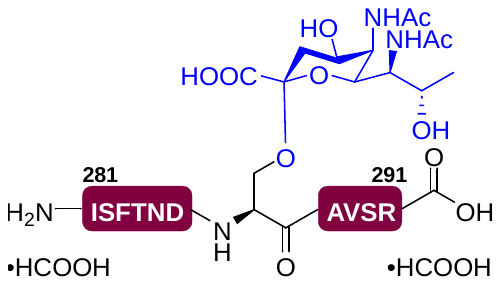

Iterative coupling of the FlgE A288-S290 fragment was performed on resin‑bound Fmoc‑protected arginine (*vide supra*) (20 µmol) according to general procedure 4. Coupling of pseudaminylated serine building block **α‑1** was then performed according to general procedure 5. Afterwards, fragment FlgE I281-D286 was completed following general procedure 4. After the final coupling, the peptide was Fmoc deprotected by treating the resin with 20% (v/v) piperidine in DMF (2x5 mL) for 5 min, then washed with DMF (5x5 mL) and CH_2_Cl_2_ (10x5 mL). The resulting peptide was then cleaved off from the resin and side chain deprotected according to general procedure 6 to generate the crude peptide. Finally, this peptide was saponified according to general procedure 7, and then purified by preparative RP-HPLC (column: Waters Sunfire C_18_ 5 μm 130 Å (19 x 150 mm), flow rate: 15 mL/min, gradient: 0%-40% B in A over 45 min). The product was lyophilised to afford the *title* compound **α‑10** as a white fluffy solid (5.2 mg, 16% over 21 steps). **LRMS** (ESI+) *m/z* [M + H]^+^: 1513.40.

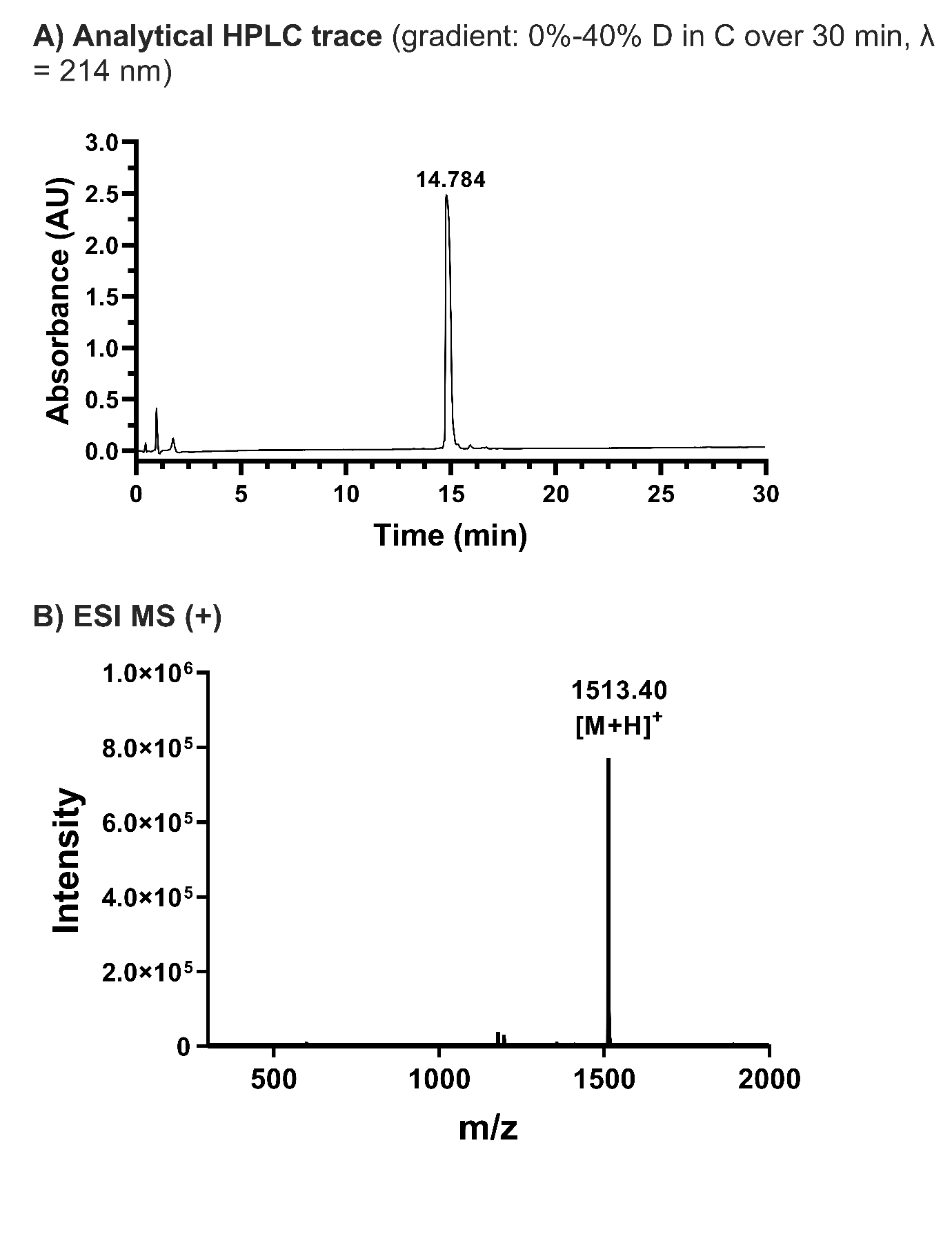

**FlgE(281-291): H_2_N-ISFTNDS(β-Pse5Ac7Ac)AVSR-OH.diformate salt (β‑10)**

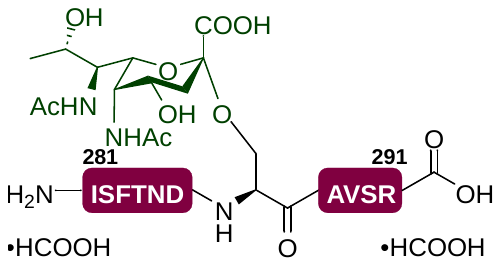

Iterative coupling of the FlgE A288-S290 fragment was performed on resin‑bound Fmoc‑protected arginine (*vide supra*) (20 µmol) according to general procedure 4. Coupling of pseudaminylated serine building block **β‑1** was then performed according to general procedure 5. Afterwards, fragment FlgE I281-D286 was completed following general procedure 4. After the final coupling, the peptide was Fmoc deprotected by treating the resin with 20% (v/v) piperidine in DMF (2x5 mL) for 5 min, then washed with DMF (5x5 mL) and CH_2_Cl_2_ (10x5 mL). The resulting peptide was then cleaved off from the resin and side chain deprotected according to 6 to generate the crude peptide. Finally, this peptide was saponified according to general procedure 7, and then purified by preparative RP-HPLC (column: Waters Sunfire C_18_ 5 μm 130 Å (19 x 150 mm), flow rate: 15 mL/min, gradient: 0%-40% B in A over 45 min). The product was lyophilised to afford the *title* compound **β‑10** as a white fluffy solid (2.1 mg, 6% over 21 steps). **LRMS** (ESI+) *m/z* [M + H]^+^: 1513.00.

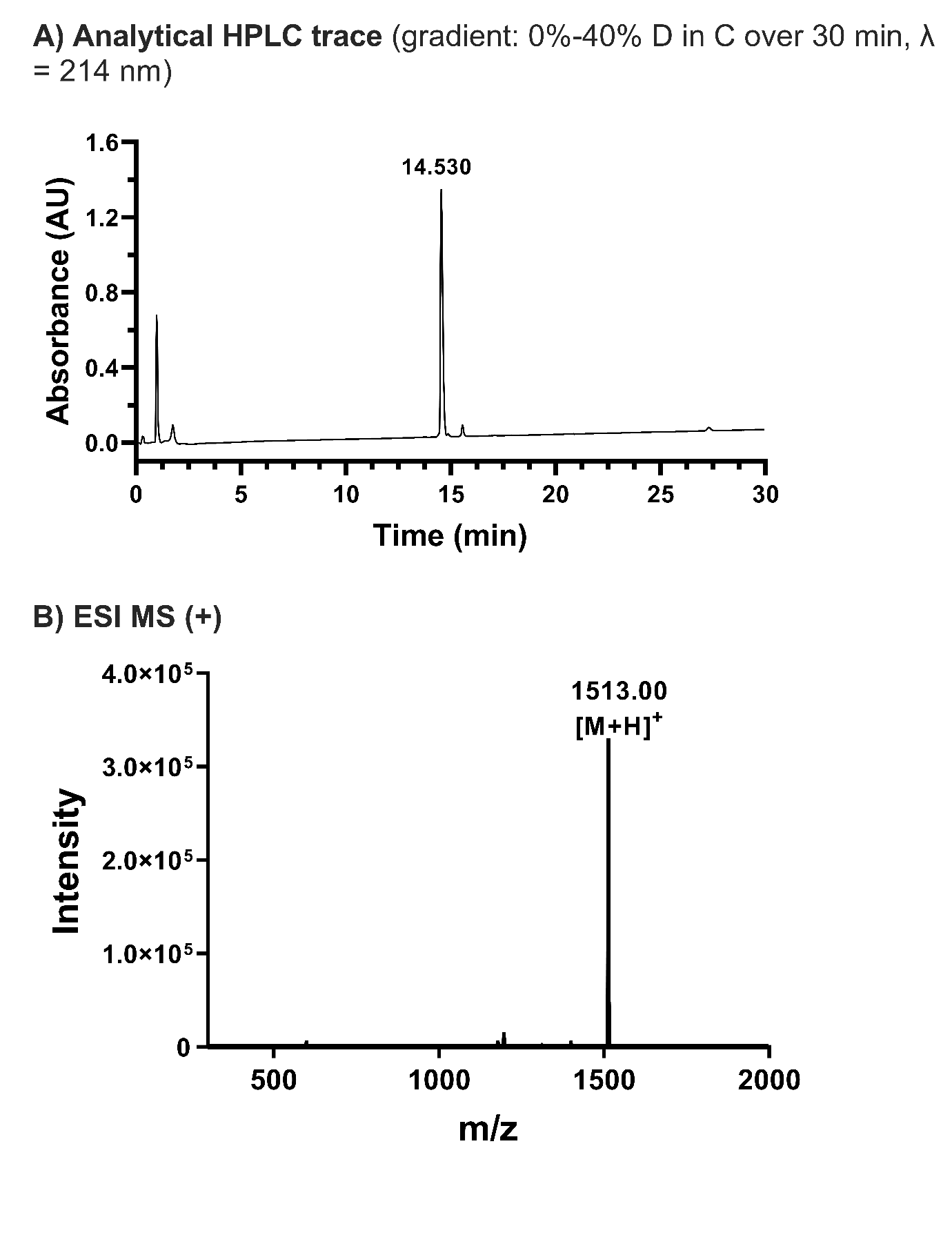

**NMR Spectra of final pseudaminylated amino acid building blocks**

Compound **α‑1 ^1^H NMR** (500 MHz, DMSO-*d*_6_)

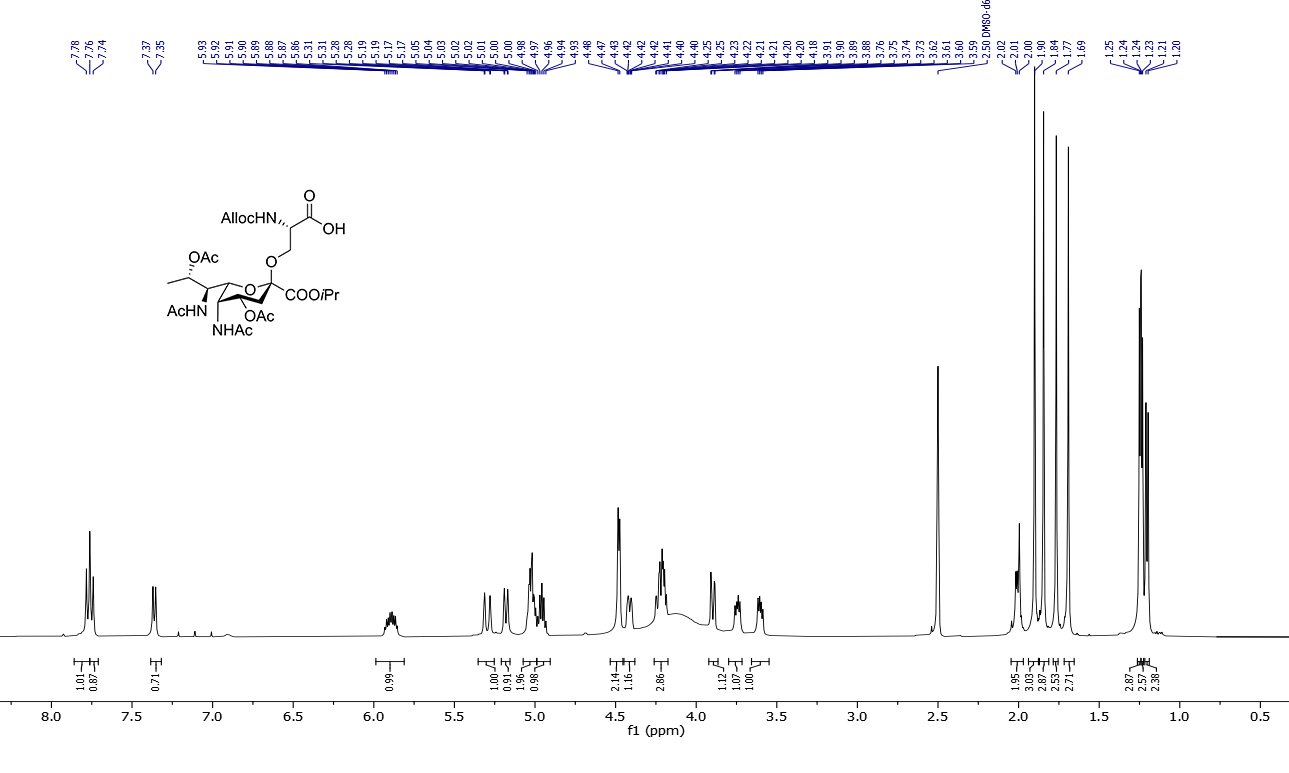

Compound **α‑1 ^13^C NMR** (126 MHz, DMSO-*d*_6_)

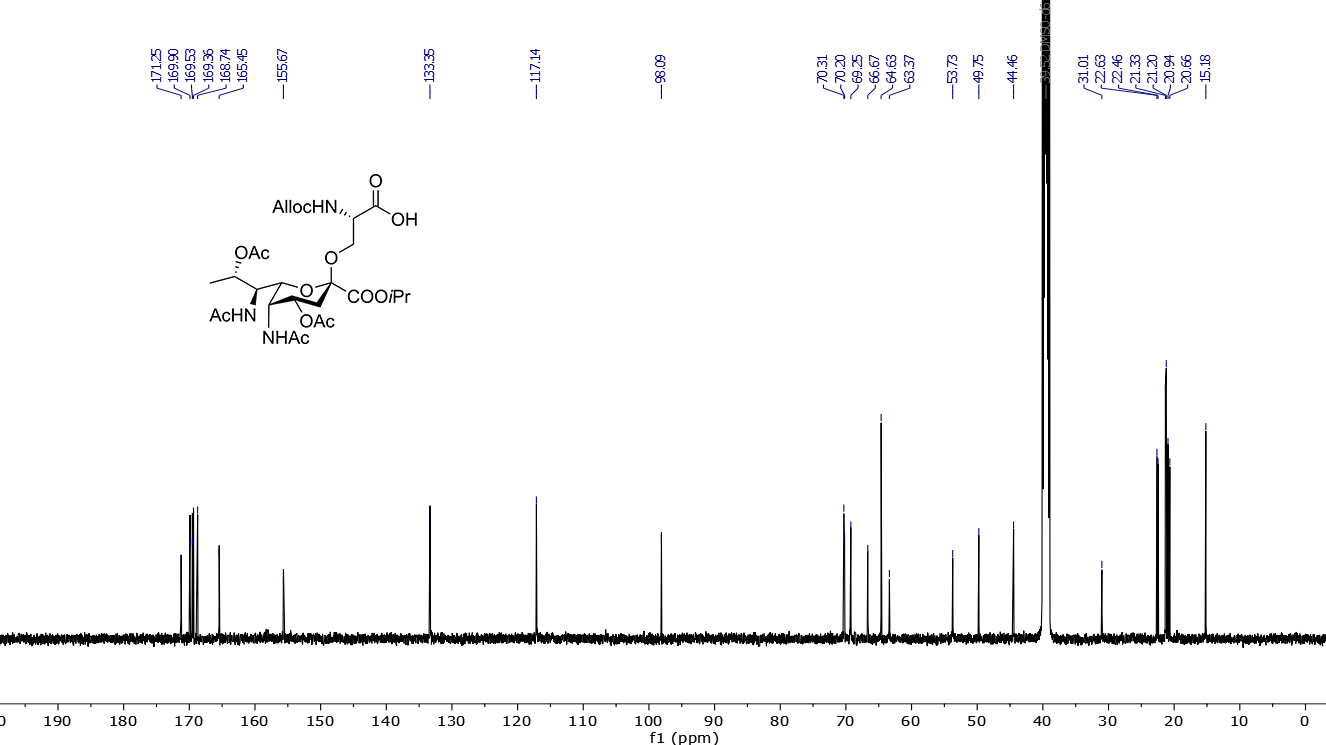

Compound **α‑1 COSY** (500 MHz, DMSO-*d*_6_)

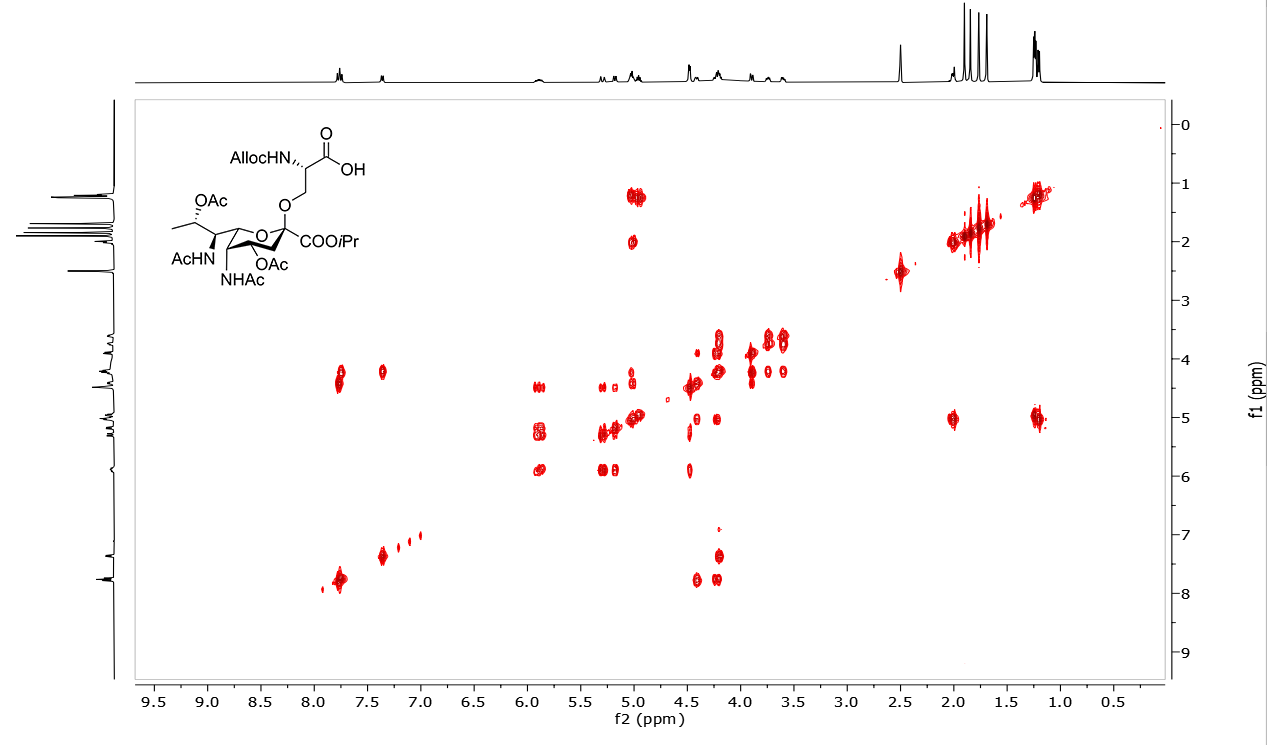

Compound **α‑1 HSQC** (500/126 MHz, DMSO-*d*_6_)

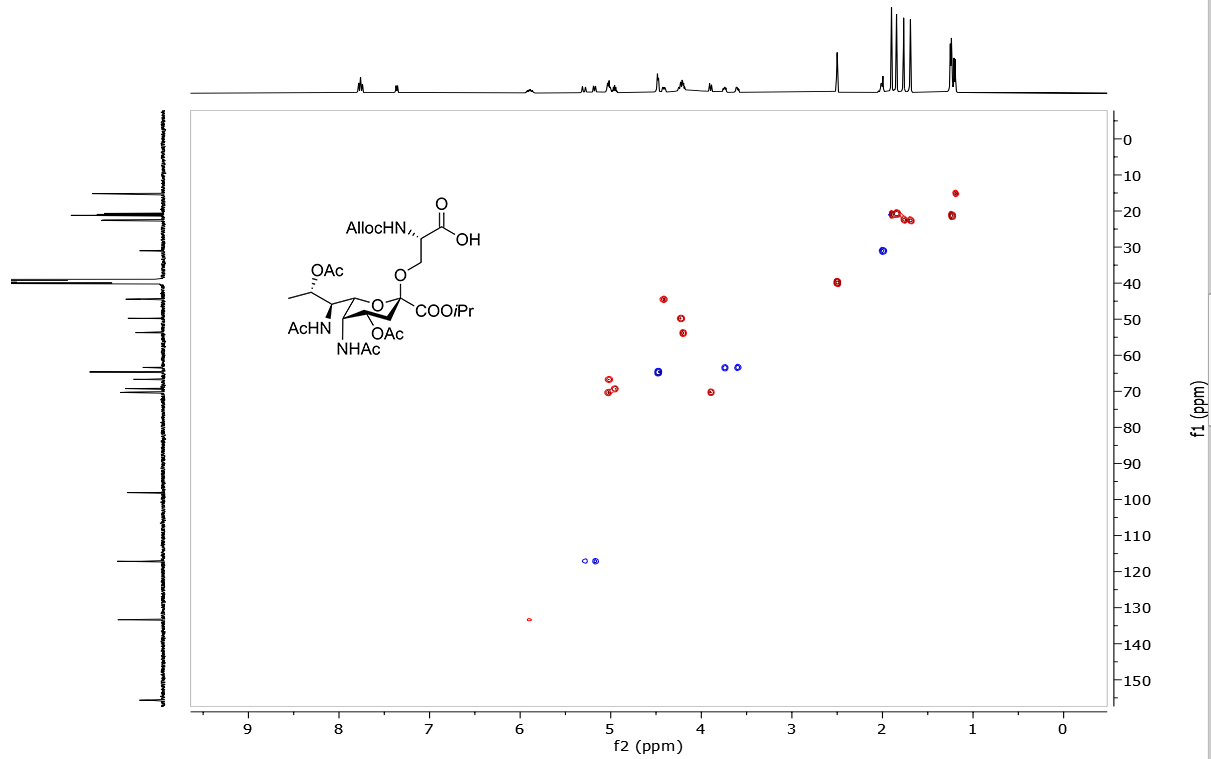

Compound **α‑1 HMBC** (500/126 MHz, DMSO-*d*_6_)

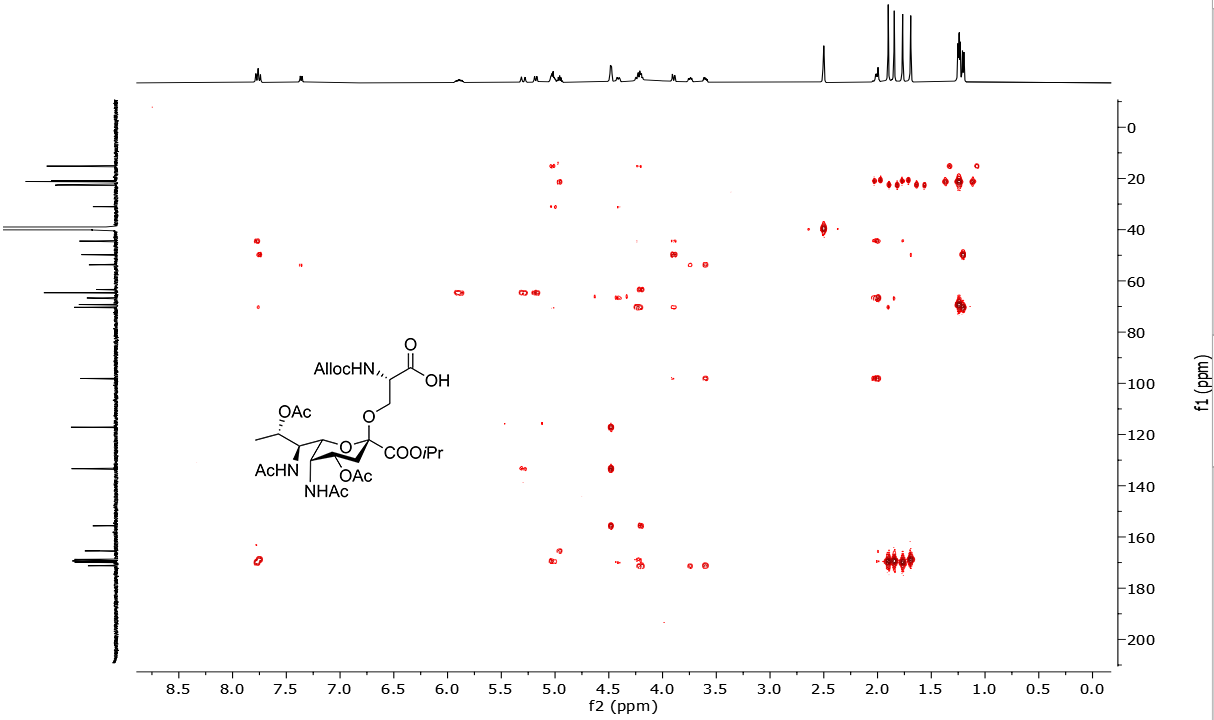

Compound **β‑1 ^1^H NMR** (400 MHz, DMSO-*d*_6_)

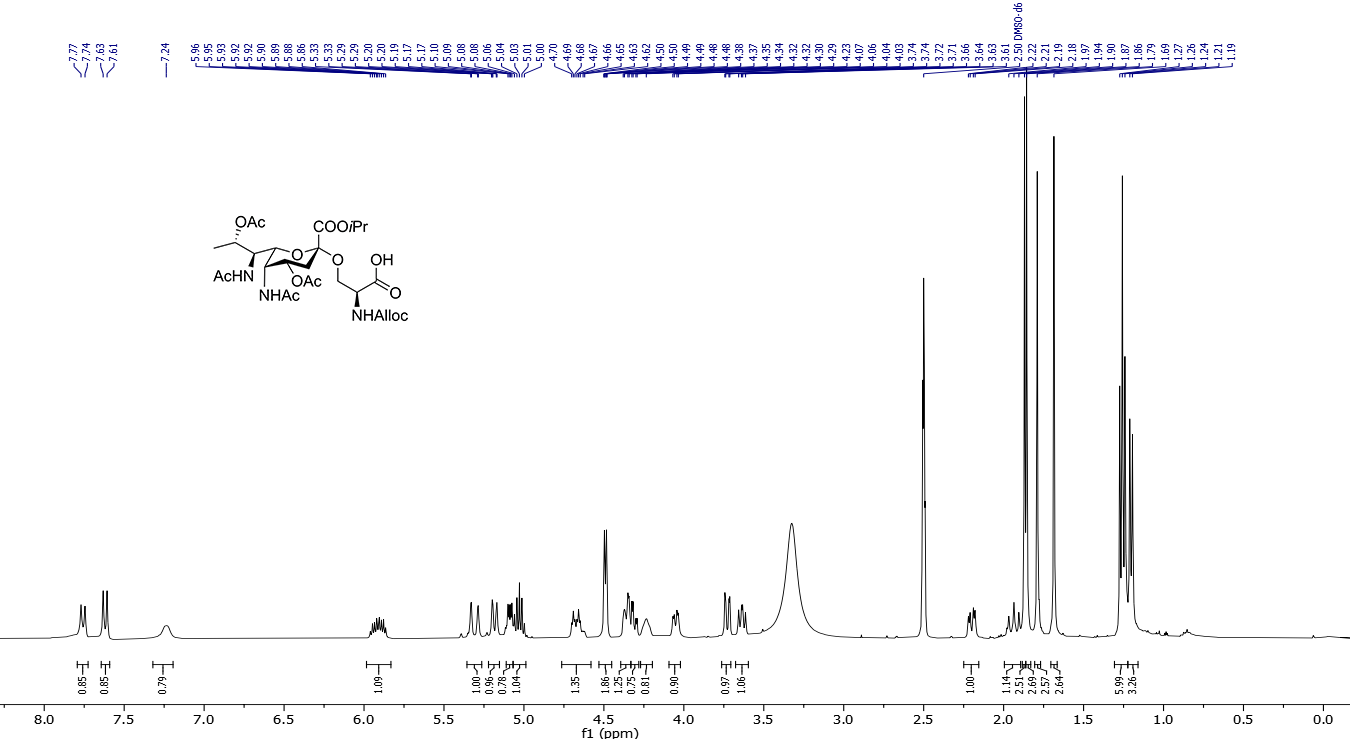

Compound **β‑1 ^13^C NMR** (101 MHz, DMSO-*d*_6_)

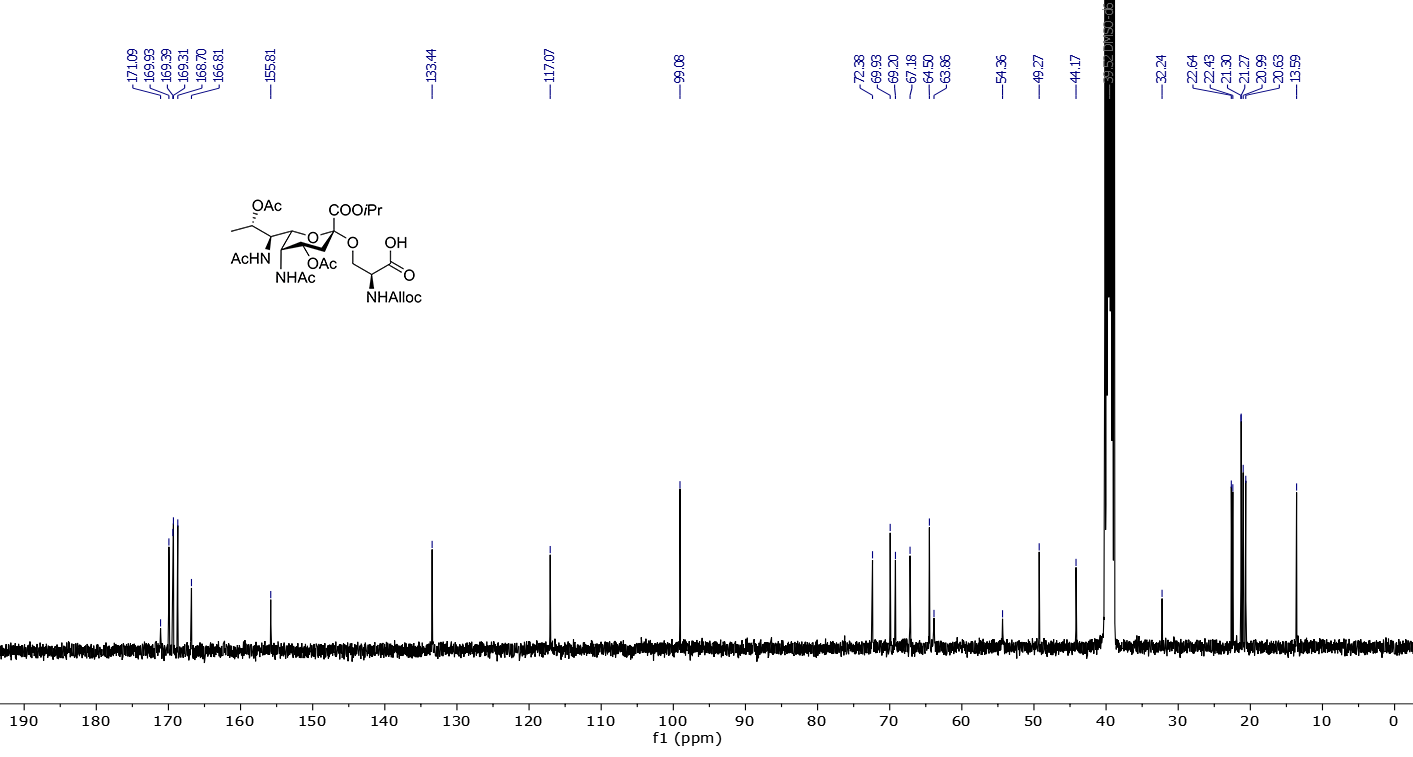

Compound **β‑1 COSY** (400 MHz, DMSO-*d*_6_)

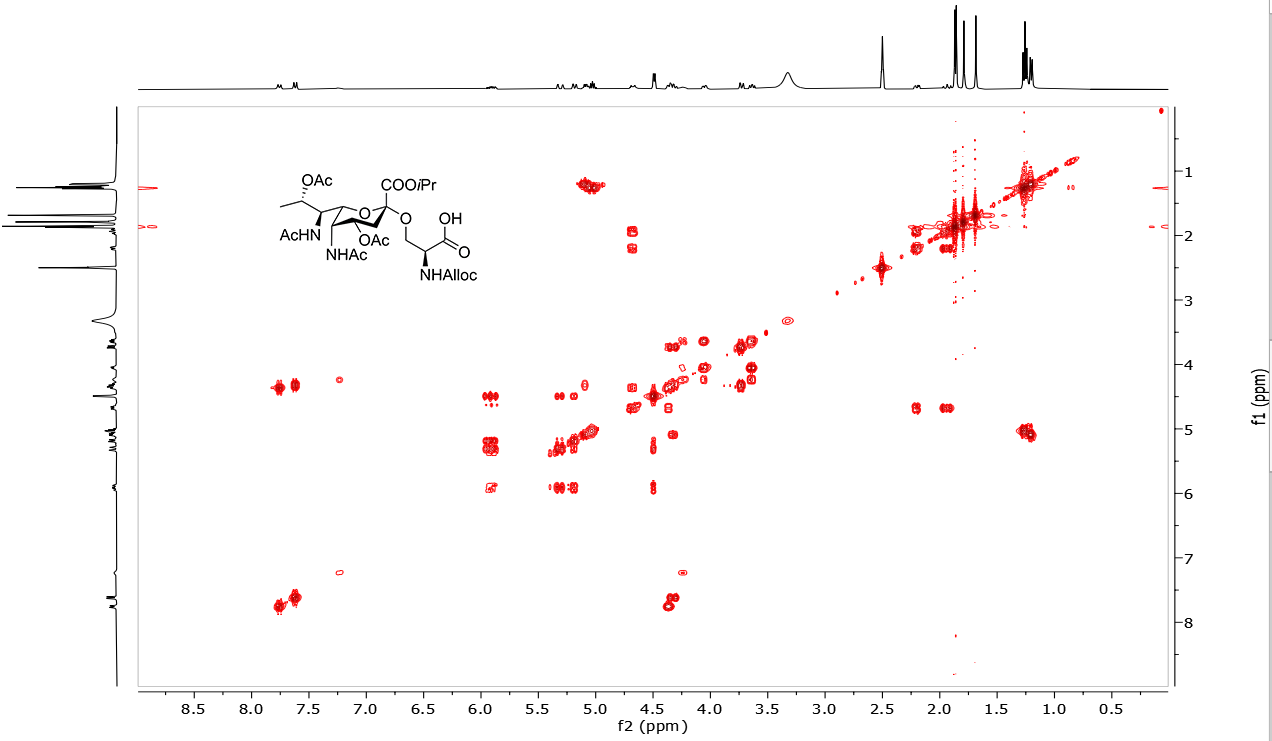

Compound **β‑1 HSQC** (400/101 MHz, DMSO-*d*_6_)

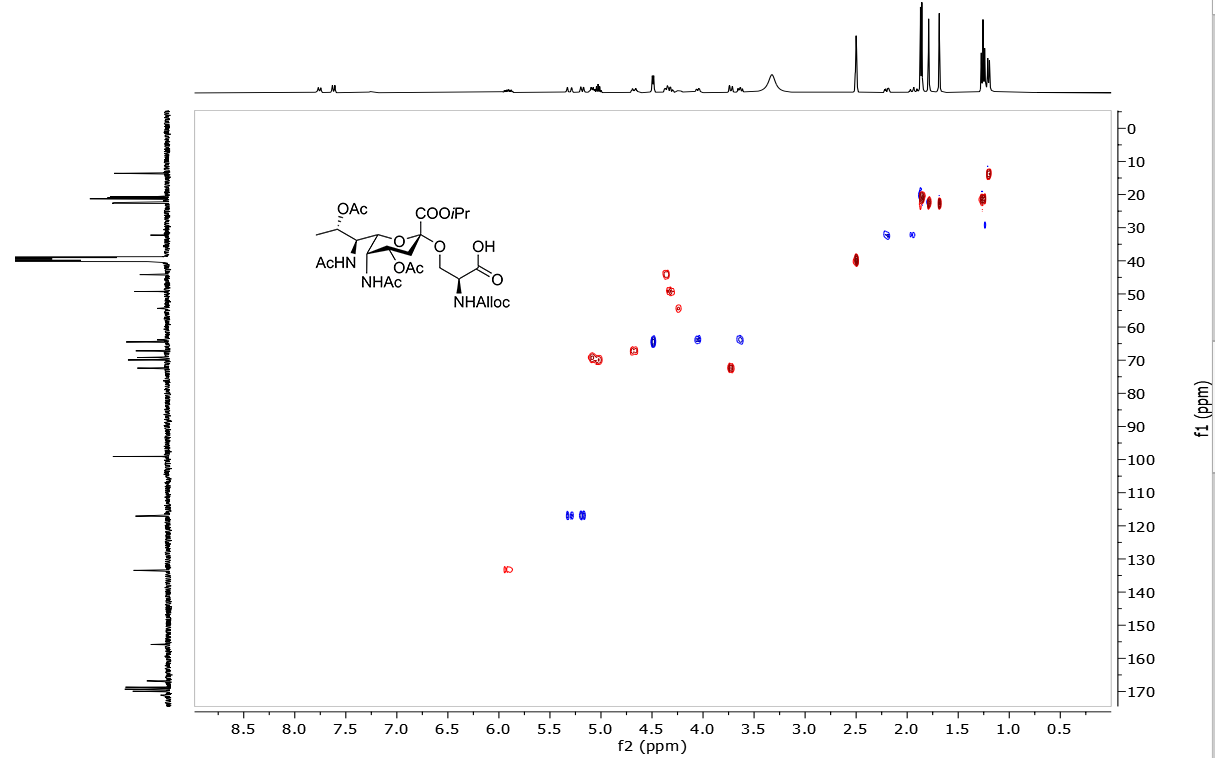

Compound **β‑1 HMBC** (400/101 MHz, DMSO-*d*_6_)

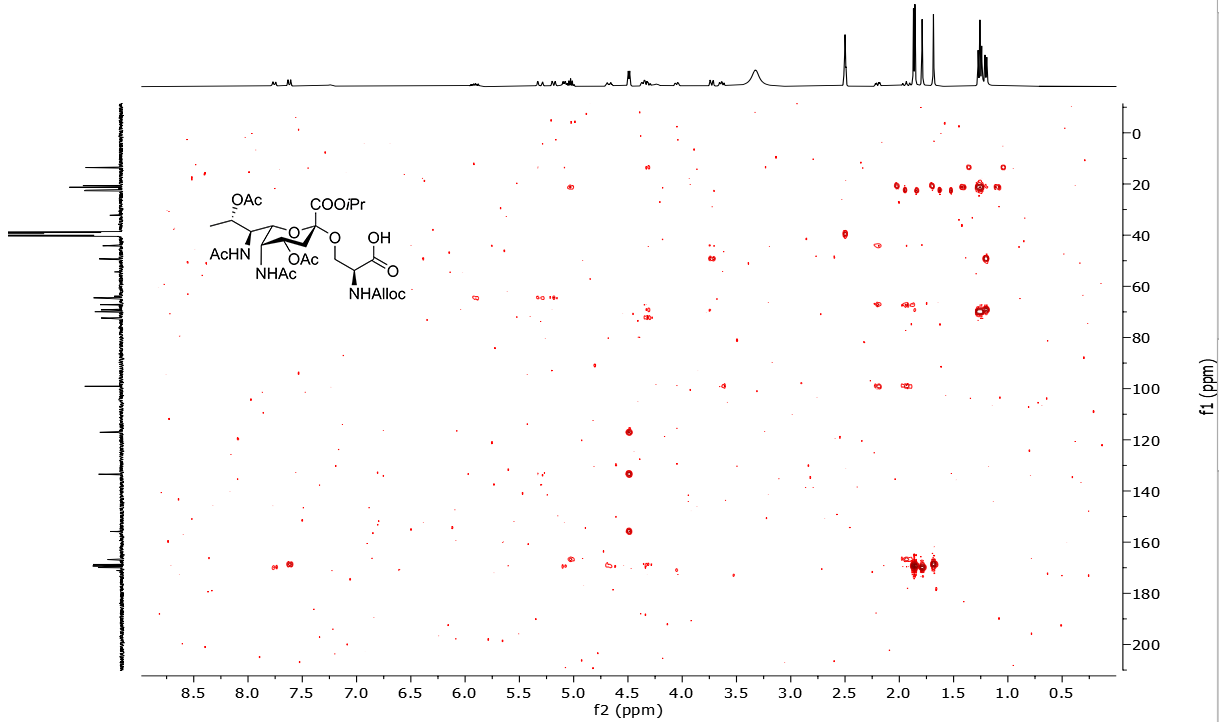

Compound **α‑S6 ^1^H NMR** (400 MHz, DMSO-*d*_6_)

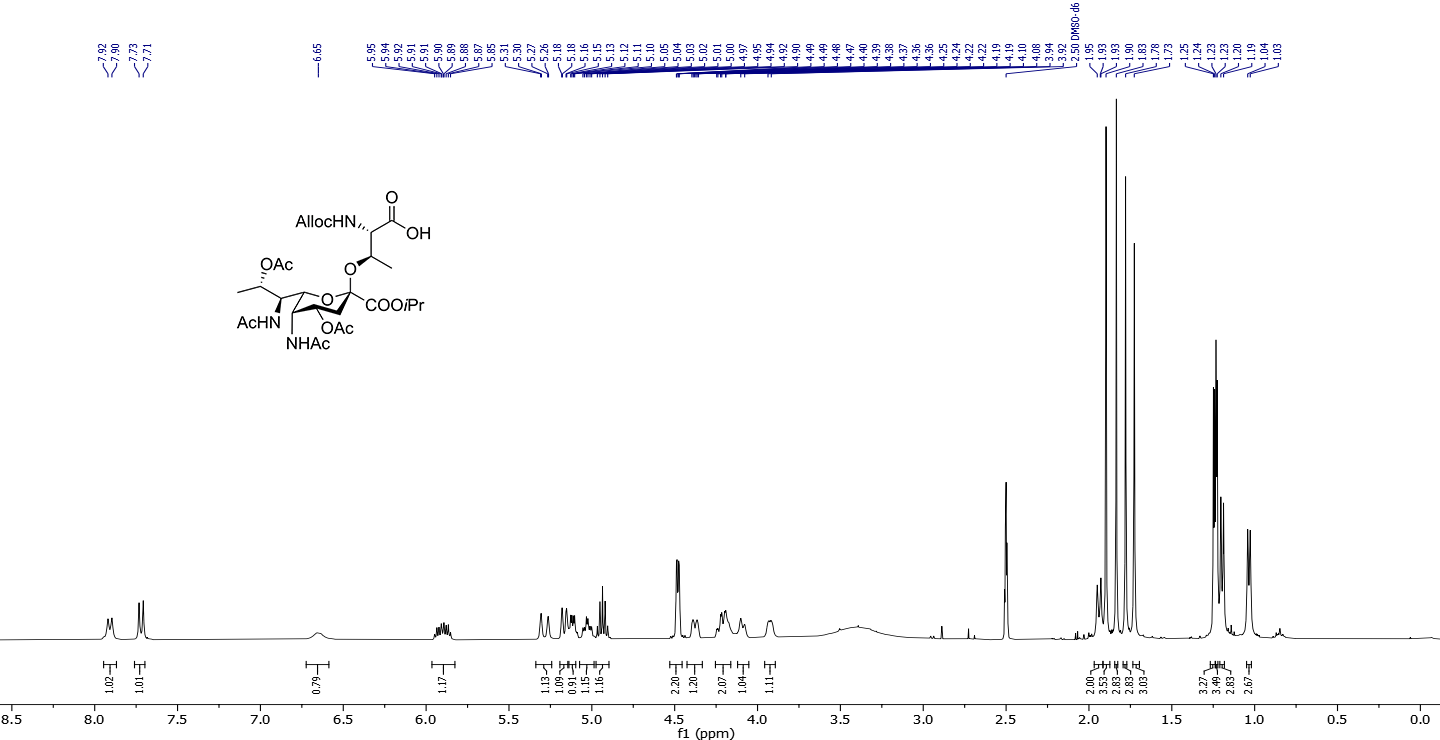

Compound **α‑S6 ^13^C NMR** (101 MHz, DMSO-*d*_6_)

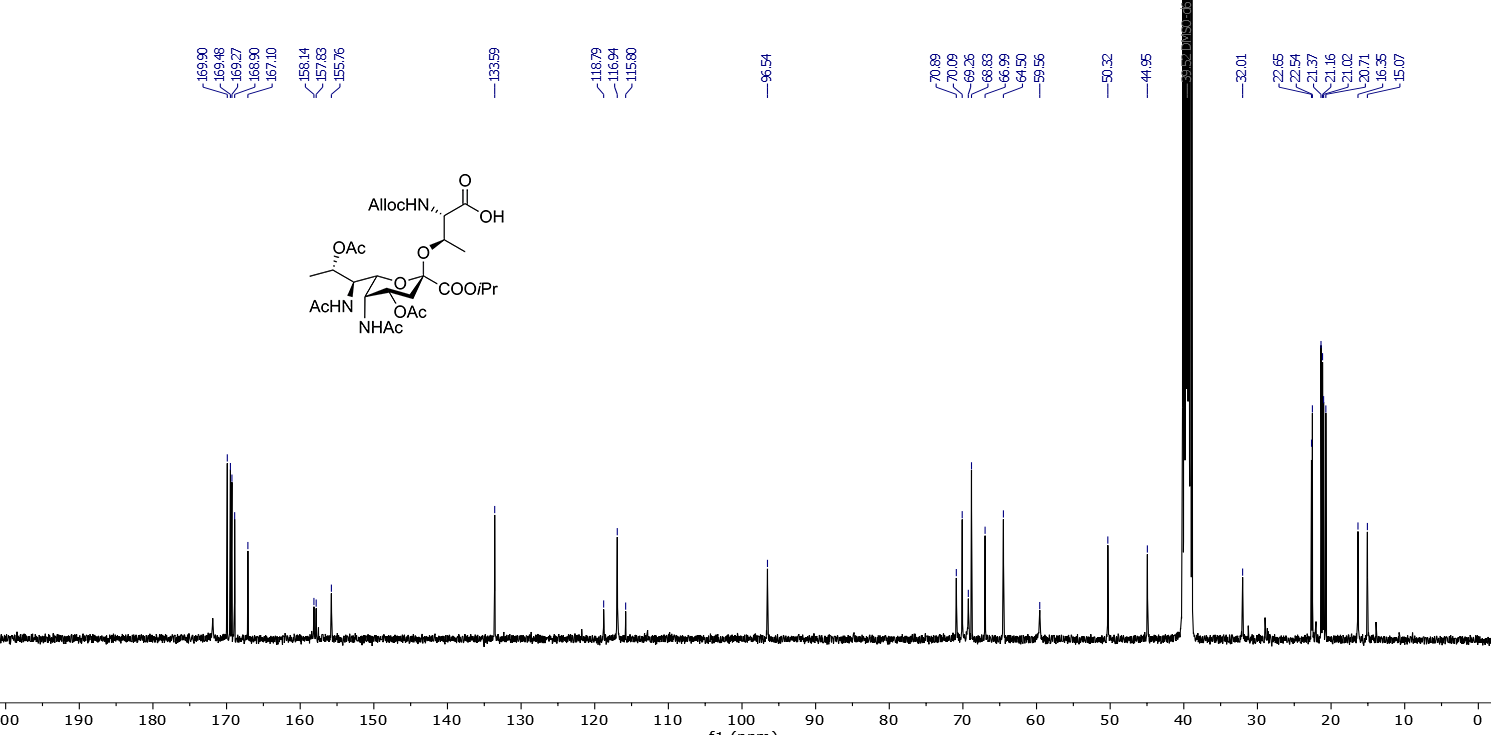

Compound **α‑S6 COSY** (400 MHz, DMSO-*d*_6_)

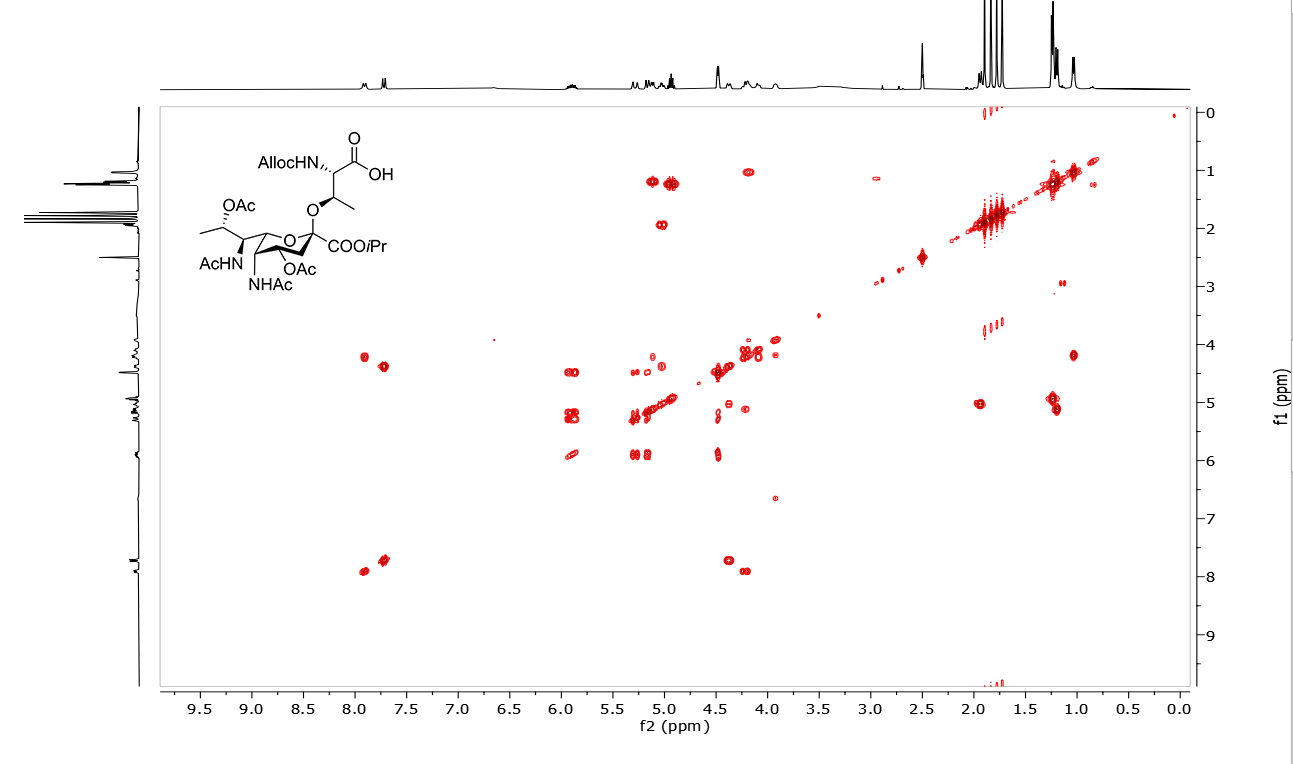

Compound **α‑S6 HSQC** (400/101 MHz, DMSO-*d*_6_)

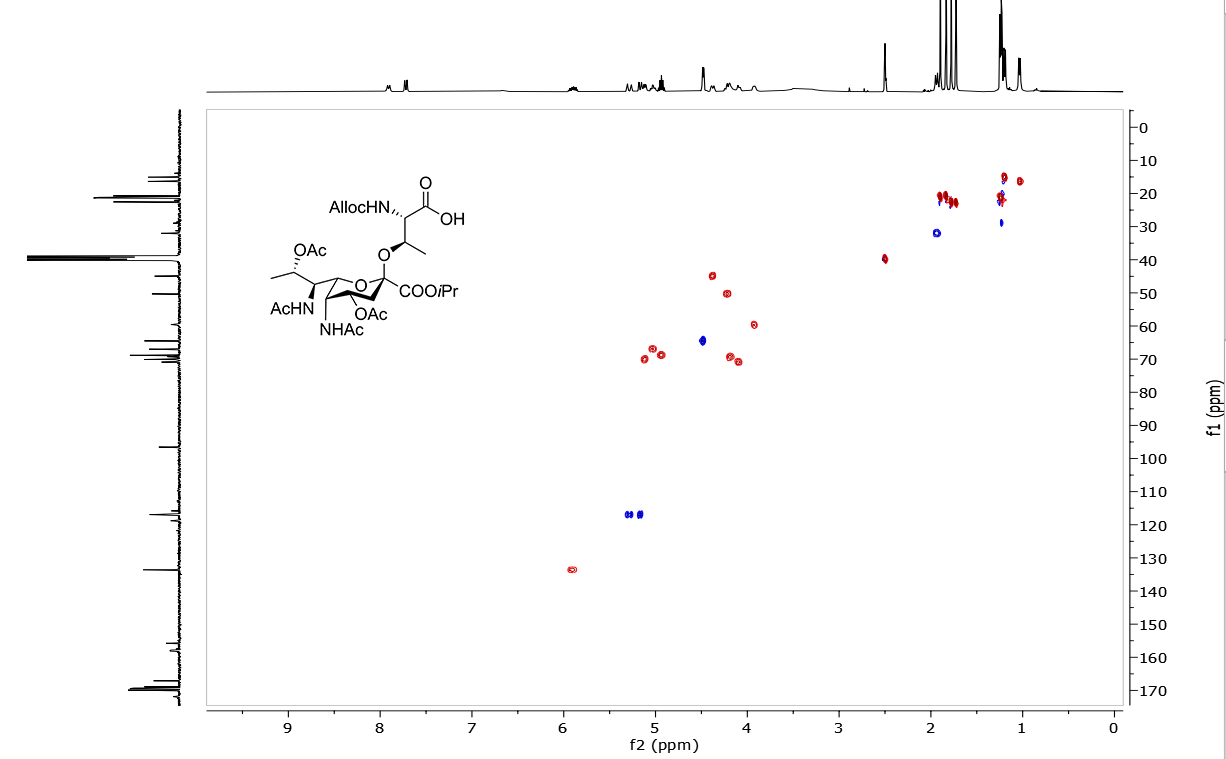

Compound **α‑S6 HMBC** (400/101 MHz, DMSO-*d*_6_)

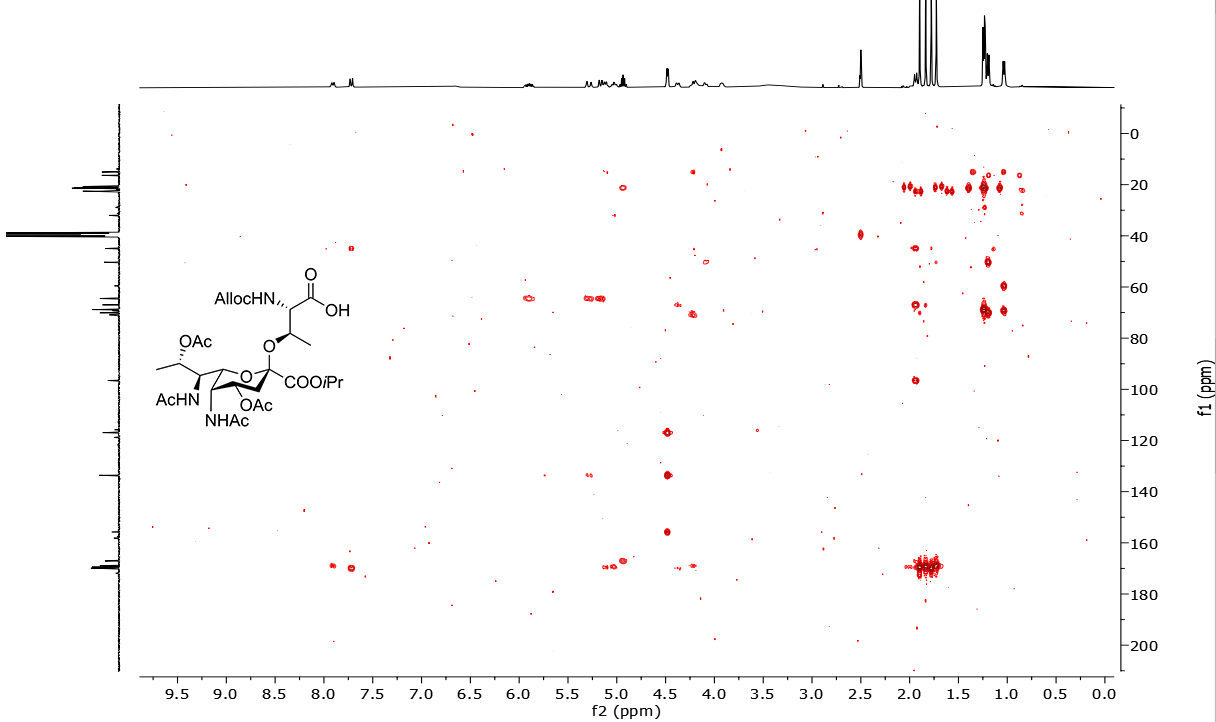

Compound **β‑S6 ^1^H NMR** (500 MHz, DMSO-*d*_6_)

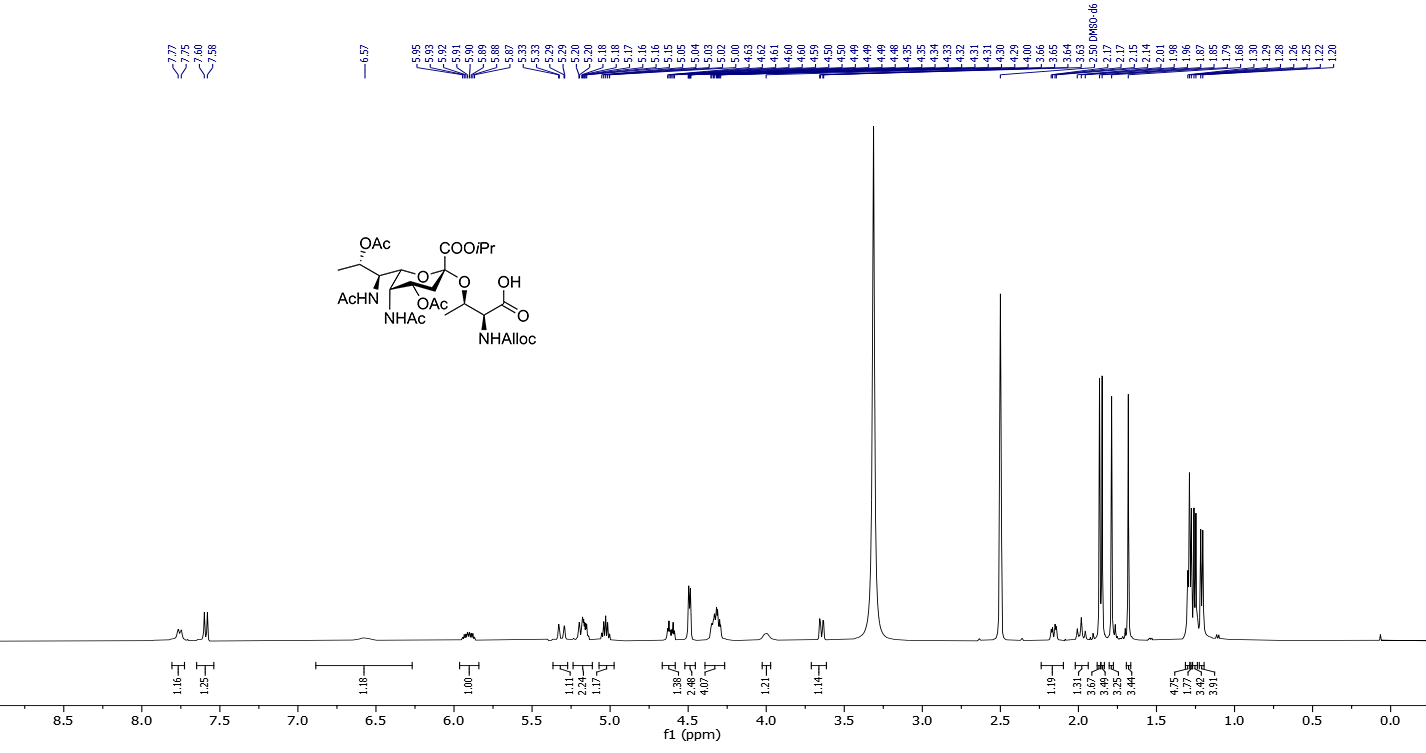

Compound **β‑S6 ^13^C NMR** (126 MHz, DMSO-*d*_6_)

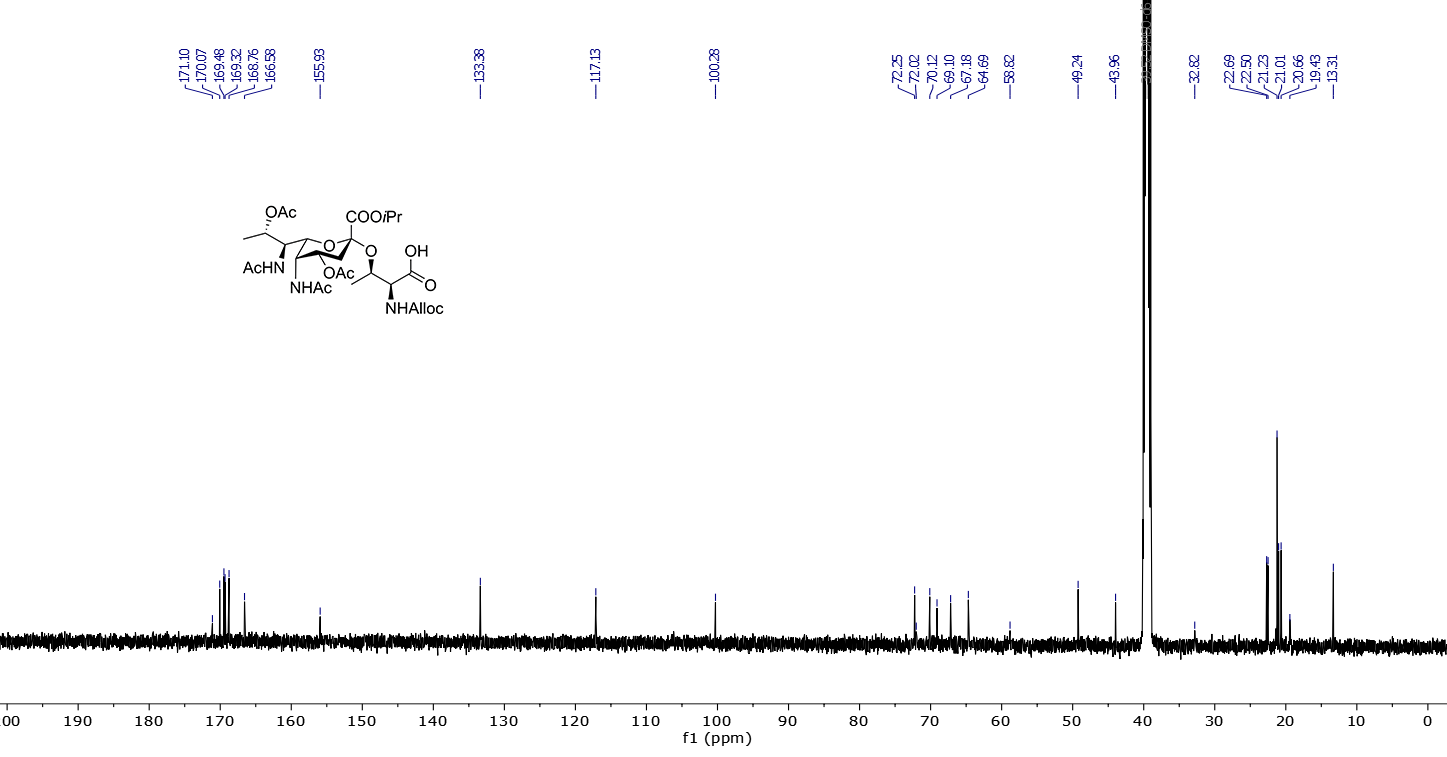

Compound **β‑S6 COSY** (500 MHz, DMSO-*d*_6_)

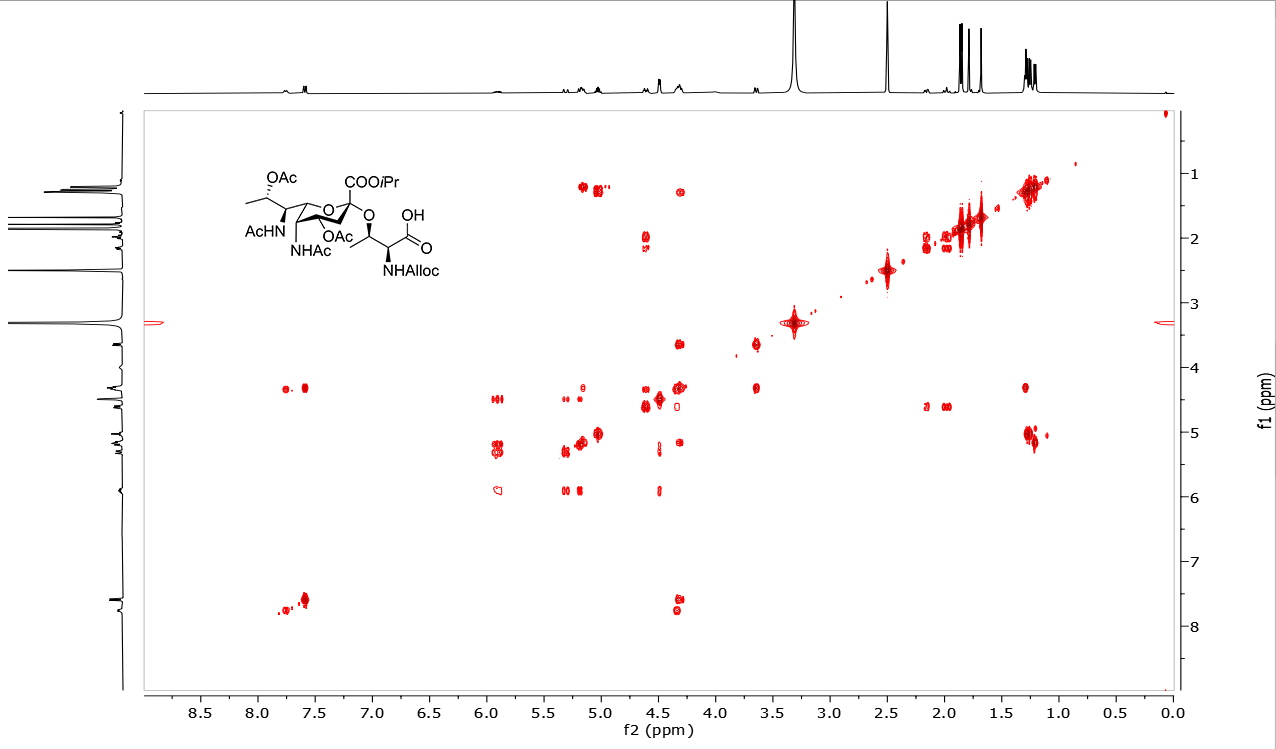

Compound **β‑S6 HSQC** (500/126 MHz, DMSO-*d*_6_)

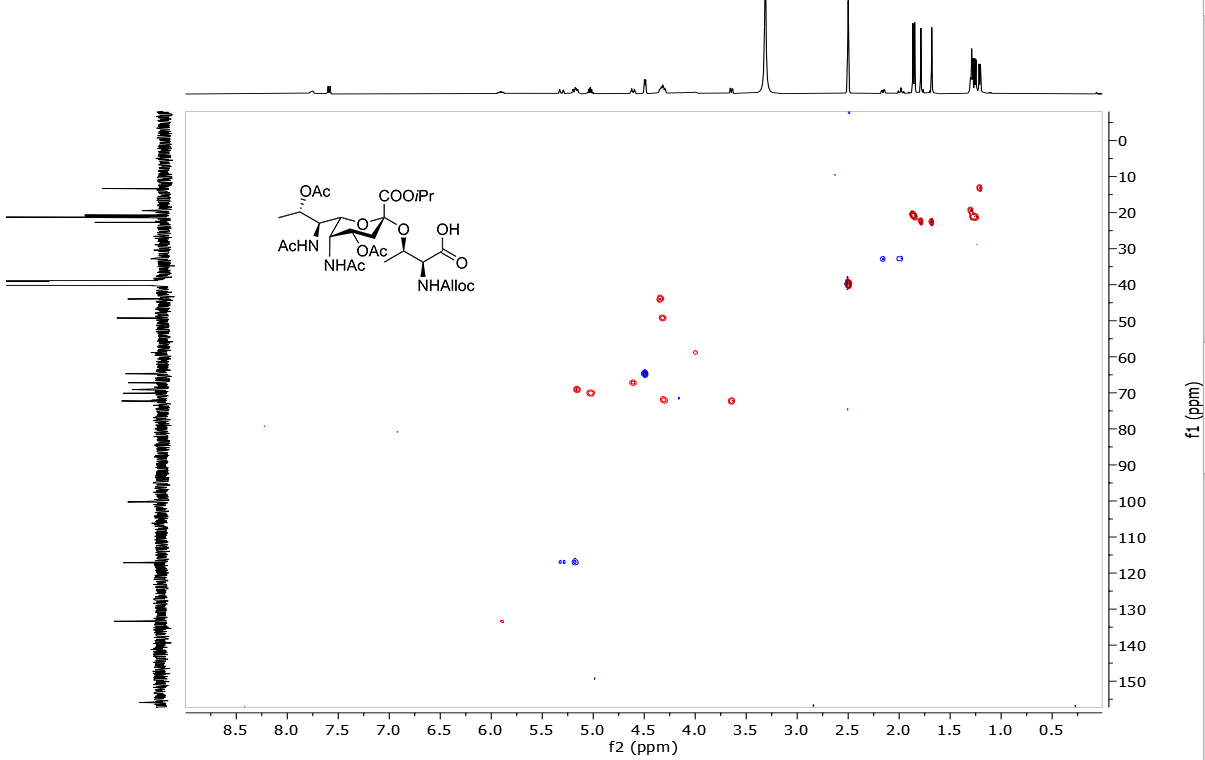

Compound **β‑S6 HMBC** (500/126 MHz, DMSO-*d*_6_)

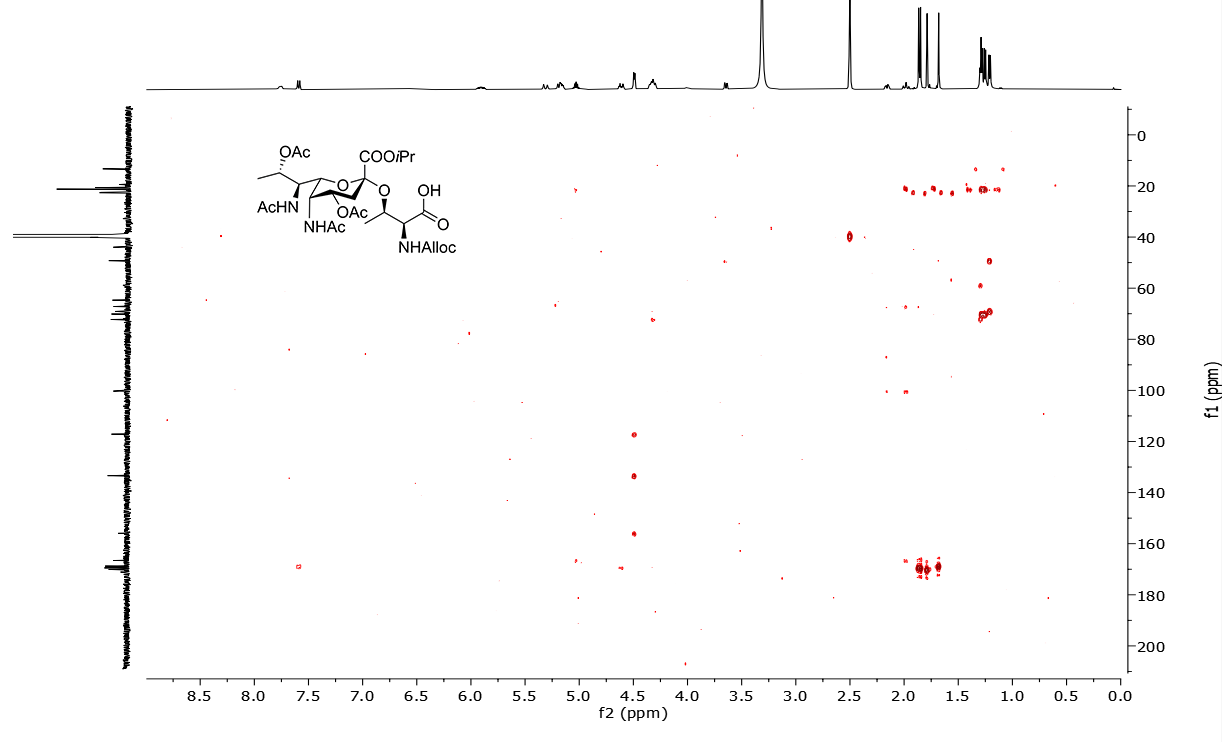
